# Pleiotropic effects of trisomy and pharmacologic modulation on structural, functional, molecular, and genetic systems in a Down syndrome mouse model

**DOI:** 10.1101/2023.07.31.551282

**Authors:** Sergi Llambrich, Birger Tielemans, Ellen Saliën, Marta Atzori, Kaat Wouters, Vicky Van Bulck, Mark Platt, Laure Vanherp, Nuria Gallego Fernandez, Laura Grau de la Fuente, Harish Poptani, Lieve Verlinden, Uwe Himmelreich, Anca Croitor, Catia Attanasio, Zsuzsanna Callaerts-Vegh, Willy Gsell, Neus Martínez-Abadías, Greetje Vande Velde

**Affiliations:** Biomedical MRI, Department of Imaging and Pathology, KU Leuven, Leuven, Belgium; Department of Human Genetics, KU Leuven, Leuven, Belgium; Laboratory of Biological Psychology, KU Leuven, Leuven, Belgium; Centre for Preclinical Imaging, Department of Molecular and Clinical Cancer Medicine, University of Liverpool, Liverpool, UK; Departament de Biologia Evolutiva, Ecologia i Ciències Ambientals (BEECA), Facultat de Biologia, Universitat de Barcelona (UB), Spain; Clinical and Experimental Endocrinology (CEE), KU Leuven, Leuven, Belgium

## Abstract

Down syndrome (DS) is characterized by skeletal and brain structural malformations, cognitive impairment, altered hippocampal metabolite concentration and gene expression imbalance. These alterations were usually investigated separately, and the potential rescuing effects of green tea extracts enriched in epigallocatechin-3-gallate (GTE-EGCG) provided disparate results due to different experimental conditions. We overcame these limitations by conducting the first longitudinal controlled experiment evaluating genotype and GTE-EGCG prenatal chronic treatment effects before and after treatment discontinuation. Our findings revealed that the Ts65Dn mouse model reflected the pleiotropic nature of DS, exhibiting brachycephalic skull, ventriculomegaly, neurodevelopmental delay, hyperactivity, and impaired memory robustness with altered hippocampal metabolite concentration and gene expression. GTE-EGCG treatment modulated most systems simultaneously but did not rescue DS phenotypes. On the contrary, the treatment exacerbated trisomic phenotypes including body weight, tibia microarchitecture, neurodevelopment, adult cognition, and metabolite concentration, not supporting the therapeutic use of GTE-EGCG as a prenatal chronic treatment. Our results highlight the importance of longitudinal experiments assessing the co-modulation of multiple systems throughout development when characterizing preclinical models in complex disorders and evaluating the pleiotropic effects and general safety of pharmacological treatments.

## INTRODUCTION

Down syndrome (DS) is a developmental disorder with structural, functional, molecular, and genetic alterations that show a dynamic onset and severity (Antonarakis et al., 2020; de Moraes et al., 2008; Grieco et al., 2015; Keeling et al., 1997; LaCombe & Roper, 2020; Llambrich, González-Colom, et al., 2022; McCarron et al., 2017; Steingass et al., 2011). The alterations associated with DS simultaneously affect multiple systems that are likely interrelated over development, with changes in one system modulating the others (Llambrich, González, et al., 2022). Many studies have provided evidence of these developmental alterations and have demonstrated the ability of dietary supplements such as epigallocatechin-3-gallate (EGCG) or green tea extracts enriched in EGCG (GTE-EGCG) to modulate these systems separately (Catuara-Solarz et al., 2016; de la Torre et al., 2016; Goodlett et al., 2020; Guedj et al., 2009; Llambrich, González, et al., 2022; Rondal, 2020; Stagni et al., 2015; Starbuck et al., 2021). However, a holistic evaluation of the simultaneous effects of trisomy and GTE-EGCG in those intertwined systems is missing. The variety in experimental setups and readouts obtained in different preclinical studies hinders the comparison and integration of results and, as a result, evidence is usually contradictory, reporting both positive and negative treatment effects. This lack of consistency can lead to biased interpretations about the etiology of the disorder and the potential effect of pharmacological treatments.

Much of this DS research was based on the Ts65Dn mouse model (Davisson et al., 1990; Reeves et al., 1995), as it was one of the first preclinical models available for DS and has been widely used for experimental testing. These mice carry a segment with approximately 120 genes homologous to Hsa21 (starting upstream of Mrpl39 to the telomeric end of Mmu16), translocated to a small centromeric part of Mmu17 (Duchon et al., 2011; Herault et al., 2017; Reinholdt et al., 2011). Ts65Dn mice are trisomic for about two-thirds of the genes orthologous to Hsa21, but also carry genes originating from the Mmu17 that are not related with DS, including about 46 protein-coding genes, 35 nonprotein-coding genes and 35 pseudogenes (Muñiz Moreno et al., 2020). These genetic alterations do not fully represent Down syndrome’s aneuploidy and other mouse and rat models have been developed recently that more faithfully represent the trisomic nature of DS (Kazuki et al., 2020; Kazuki et al., 2022). However, in this study we used the Ts65Dn mouse model because it recapitulates the main skeletal, brain, cognitive, brain metabolite, and genetic alterations associated with DS (Blazek et al., 2011; Costa et al., 2010; Dierssen et al., 2002; Escorihuela et al., 1998; Gupta et al., 2016; Huang et al., 2000; Llambrich, González, et al., 2022; Même, 2014; Starbuck et al., 2021); and the effects of GTE-EGCG pharmacological modulation have been extensively evaluated using this mouse model (Catuara-Solarz et al., 2016; Goodlett et al., 2020; Jamal et al., 2022; Llambrich, González, et al., 2022; McElyea et al., 2016; Stagni et al., 2016; Starbuck et al., 2021; Stringer et al., 2017).

At the structural level, people with DS show skeletal and brain alterations that progress through ontogeny (Antonarakis et al., 2020; de Moraes et al., 2008; Fischer-Brandies et al., 1986; Keeling et al., 1997; LaCombe & Roper, 2020; Pearlson et al., 1998; Starbuck et al., 2021). Children with DS have a decreased buildup of bone mass and a low bone turnover rate, resulting in more osteoclast than osteoblast activity, smaller bone area, lower bone mineral density (BMD), and an increased risk of osteoporosis at adulthood (Carfì et al., 2017; Keeling et al., 1997; LaCombe & Roper, 2020). Individuals with DS also present midfacial hypoplasia and flattened nasal bridge along with skull malformations resulting in a shorter, wider, and rounder skull (brachycephaly) (Blazek et al., 2011; LaCombe & Roper, 2020; Richtsmeier et al., 2000; Suri et al., 2010; Thomas & Roper, 2021). Previous studies have shown the potential of GTE-EGCG to modulate craniofacial and postcranial morphology, as well as the microarchitecture and bone mineral density of the long bones, showing positive, negative or no treatment effects (Abeysekera et al., 2016; J. D. Blazek et al., 2015; Jamal et al., 2022; Llambrich, González-Colom, et al., 2022; McElyea et al., 2016; Starbuck et al., 2021; Stringer et al., 2017). We previously detected dose-, time- and anatomical structure-dependent effects in a study using the same mouse model and the same treatment regime with two different GTE-EGCG doses (Llambrich, González-Colom, et al., 2022). A dose of 100 mg/kg/day of GTE-EGCG exacerbated facial dysmorphologies (Starbuck et al., 2021), modified the skeletal dysmorphologies associated with DS without rescuing the bones shape (Llambrich, González-Colom, et al., 2022), and altered the integration between the skull and the brain (Llambrich, González, et al., 2022). However, a lower dose of 30 mg/kg/day significantly reduced the facial dysmorphologies (Starbuck et al., 2021) but did not show additional rescuing effects in other skeletal traits (Llambrich, González-Colom, et al., 2022).

In DS, the craniofacial size and shape alterations are accompanied with structural brain alterations. People with DS show a reduced overall brain volume from birth, with disproportionately smaller hippocampus and cerebellum, and larger ventricles (Hamner et al., 2018; Movsas et al., 2016; Patkee et al., 2020; Pinter et al., 2001; Rodrigues et al., 2019; Smigielska-Kuzia et al., 2011). Evidence for the effects of GTE-EGCG on brain anatomy is limited. Our own previous study showed that administration of 100 mg/kg/day of GTE-EGCG altered the brain shape of Ts65Dn mice at adulthood (Llambrich, González, et al., 2022), while another study administering green tea polyphenols corresponding to 0.6 - 1 mg EGCG per day in YACtg152F7 mice showed reduced thalamic-hypothalamic volume and reduced overall brain weight and volume (Guedj et al., 2009).

The structural brain alterations in DS are associated with cognitive disabilities, causing functional disability (Guidi et al., 2011; Rodrigues et al., 2019; Stagni et al., 2016). People with DS show relative impairment in executive function, short-term memory, working memory, and explicit long-term memory (Grieco et al., 2015; Lott & Dierssen, 2010; McCarron et al., 2017; Real de Asua et al., 2015; Steingass et al., 2011; Zis & Strydom, 2018); together with a neurodevelopmental delay in the acquisition of both gross and fine motor skills during childhood (Beqaj et al., 2017; Ferreira-Vasques & Lamônica, 2015; Frank & Esbensen, 2015; Kim et al., 2017; Malak et al., 2015). Regarding cognition, the therapeutic results of green tea extracts are also contradictory. A study administering 60 mg/kg GTE-EGCG to 3- to 4-month-old mBACtgDyrk1a and Ts65Dn mice for 4 weeks rescued working memory (Souchet et al., 2015), and interventional studies in humans with DS have shown the ability of (GTE-)EGCG to improve memory recognition, working memory, inhibitory control, and adaptive behavior (de la Torre et al., 2016; De la Torre et al., 2014). However, other studies found no effect or negative effects of (GTE-)EGCG on cognition. In children aged 6-12 years, a randomized phase Ib clinical trial administering 10 mg/kg/day of EGCG for 6 months combined with a dietary supplement did not improve cognition and functionality (Cieuta-Walti et al., 2022). In mice, GTE-EGCG at a dose of 30 mg/kg/day had no effect on visuospatial learning and memory, and even resulted in reduced swimming speed in the Morris water maze and increased thigmotaxic behavior in 5- to 6-month-old Ts65Dn mice (Catuara-Solarz et al., 2015).

Several studies have shown that the cognitive functional alterations observed in DS are related with changes at a molecular level on the concentration of hippocampal metabolites, as people with DS show increased myo-inositol levels when compared to euploid population, and people with DS and dementia show reduced levels of N-acetylaspartate (NAA) when compared to people with DS without dementia (Beacher et al., 2005; Lamar et al., 2011). These findings warrant further investigation into the potential role of hippocampal metabolites in cognitive function, particularly given that few studies to date have investigated the effects of trisomy on the concentration of the main metabolites in the hippocampus of Ts65Dn mice (Huang et al., 2000; Même, 2014; Santin et al., 2014) and there is currently no evidence on the effects of (GTE-)EGCG treatment.

Finally, all these structural, functional and molecular alterations are the result of a complex genetic imbalance involving the triplicated genes in chromosome 21 and the interactions of many genes across the genome (Olson et al., 2004). Indeed, skeletal malformations can be associated with the dysregulation of genes such as *Regulator of Calcineurin 1 (RCAN1), Superoxide Dismutase 1 (SOD1), Engrailed Homeobox 2 (EN2), ETS Proto-Oncogene 2 (ETS2), Sonic Hedgehog (SHH), Dual-Specificity Tyrosine-(Y)-Phosphorylation Regulated Kinase 1A (DYRK1A), SRY-Box Transcription Factor 9 (SOX9), and Orthodenticle Homeobox 2 (OTX2)* (Arron et al., 2006; Billingsley et al., 2013; McElyea et al., 2016; Roper et al., 2009; Thomas & Roper, 2021; Weisfeld-Adams et al., 2016), while cognitive impairment and altered brain development may be associated with *RCAN1, SOD1, Oligodendrocyte Transcription Factor 1 (OLIG1), Oligodendrocyte Transcription Factor2 (OLIG2), SIM BHLH Transcription Factor 2 (SIM2), Down Syndrome Cell Adhesion Molecule (DSCAM), DYRK1A, Down Syndrome Critical Region 1 (DSCR1), Synaptojanin 1 (SYNJ1)* and *Potassium Inwardly Rectifying Channel Subfamily J Member 6 (KCNJ6)* (Chang et al., 2003; Hoeffer et al., 2007; Kazemi et al., 2016; Kleschevnikov et al., 2017; Lana-Elola et al., 2011; Stagni & Bartesaghi, 2022; Stagni et al., 2018). From these genes, *DYRK1A*, a gene involved in both skeletal and neuronal development that is overexpressed by trisomy of chromosome 21 (Joshua D. Blazek et al., 2015; Dierssen & de Lagrán, 2006), has been proposed as a target gene for therapy (Atas-Ozcan et al., 2021; Becker et al., 2014; Dierssen & de Lagrán, 2006; García-Cerro et al., 2014; Jarhad et al., 2018; Ji et al., 2015; Lee et al., 2009; Rondal, 2020). The potential of EGCG to modulate gene expression in the Ts65Dn mouse model for DS has limited evidence, with one study administering 200 mg/kg EGCG on embryonic days 7 and 8 twice daily showing decreases in *Protein patched homolog 1* (*Ptch)* and *Ets2* RNA expression and significant increases in *Rcan1* and *Shh* RNA expression in the first pharyngeal arch of Ts65Dn mice at embryonic day (E) 9.5 (McElyea et al., 2016).

Since a holistic evaluation of these systems is missing, we designed a longitudinal experimental setup to follow the simultaneous development of structural, functional, molecular, and genetic alterations in the Ts65Dn mouse model. In this study, we conducted a multi-modal *in vivo* imaging study using micro computed tomography (µCT), magnetic resonance imaging (MRI) and magnetic resonance spectroscopy (MRS) to investigate the integrated development of craniofacial shape, BMD, brain volumes, and hippocampal metabolites in wildtype and Ts65Dn mice. Additionally, we evaluated the changes in body weight and performed a battery of neurodevelopmental and adult cognitive tests to assess cognitive function from birth to adulthood throughout development. At endpoint, we also evaluated tibia microarchitecture from *ex vivo* µCT scans and used RNAseq to analyze cerebellar gene expression in the same mice at eight months old (Fig. 1). Furthermore, we evaluated the pleiotropic effects of prenatal chronic GTE-EGCG treatment and explored the effects of treatment discontinuation on these systems, providing the first controlled longitudinal study assessing the simultaneous effects of treatment across different systems.

**Figure 1.**
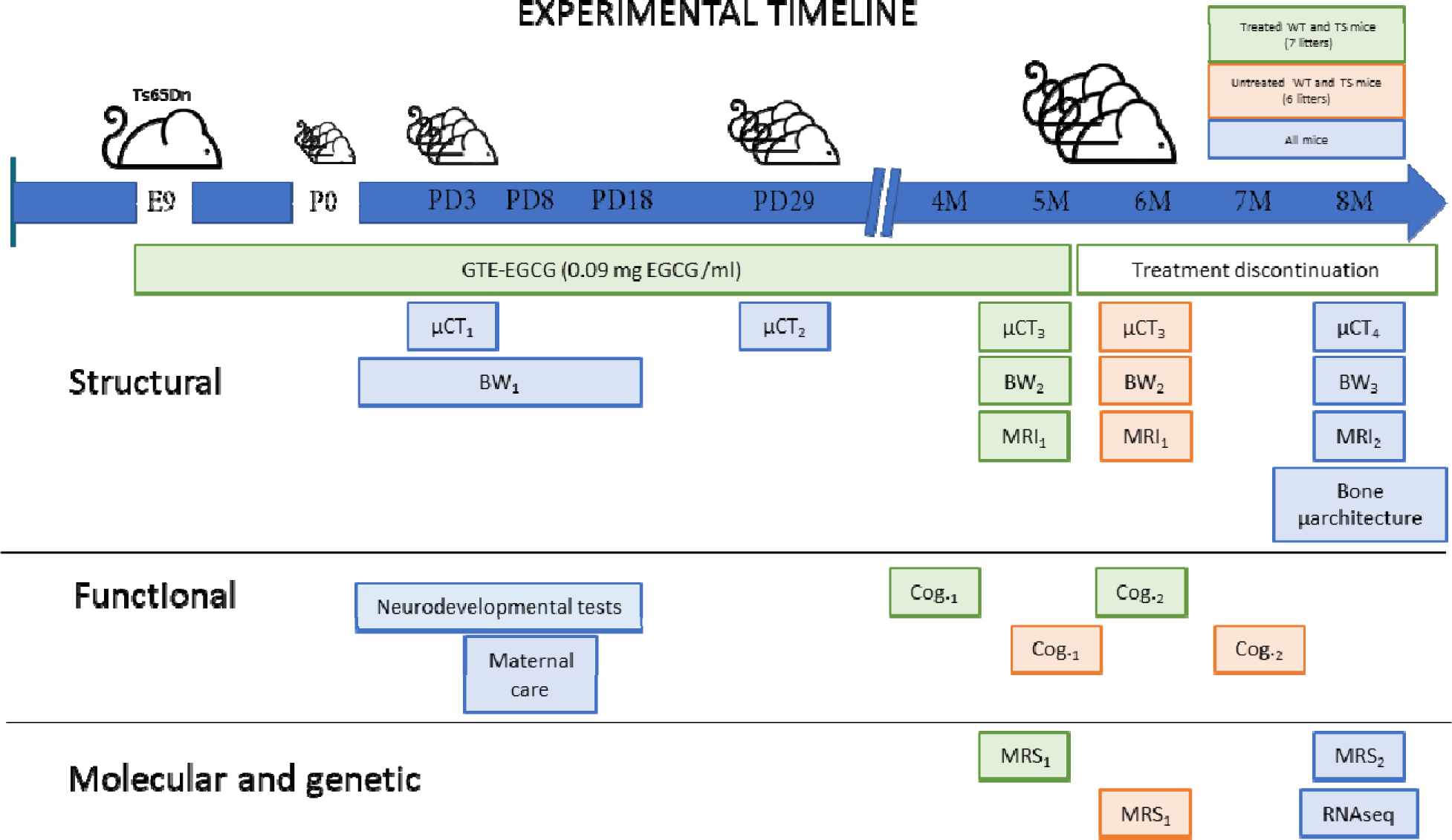
Experimental design comparing structural, functional, molecular, and genetic characteristics of wildtype (WT) and Ts65Dn (TS) mice over development and evaluating the effects of a prenatal chronic GTE-EGCG treatment and its discontinuation. Ts65Dn and WT littermates were treated prenatally at a concentration of 0.09 mg EGCG/mL, which corresponds to a dose of 30 mg/kg/day, from embryonic day 9 (E9) until mice were 5 months old (5M). A battery of neurodevelopmental tests was performed daily from postnatal day (PD) 1 to PD18 to evaluate early cognitive development. To monitor maternal care, mice home cages were recorded at PD8 for 24h. Mice body weight (BW) was recorded daily from PD1 to PD17 (BW_1_), and at two additional times before µCT scanning (BW_2_ and BW_3_). *In vivo* micro-computed tomography (µCT) scans were performed four times over development to follow skeletal development: µCT_1_ at PD3 and µCT_2_ at PD29 in all mice; µCT_3_ at 5M in treated mice, and 6M in untreated mice; and µCT_4_ at 8M in all mice. Additionally, mice were scanned with *in vivo* magnetic resonance imaging (MRI) and magnetic resonance spectroscopy (MRS) before and after treatment discontinuation to quantify brain volumetric changes and metabolite concentrations in the hippocampal region: MRI_1_ and MRS_1_ at 5M in treated mice and 6M in untreated mice; and MRI_2_ and MRS_2_ at 8M in all mice. Two batteries of cognitive tests of one-month duration each were performed to evaluate adult cognition: Cog._1_ at 4M in treated mice, and 5M in untreated mice; and Cog._2_ at 6M in treated mice, and 7M in untreated mice. At endpoint (8M), the tibia of all mice was collected to measure its length using a digital caliper and analyze its microarchitecture using *ex vivo* µCT. At this last stage, cerebellar tissue was also collected to perform RNAseq gene expression analysis.

## RESULTS

Our longitudinal experimental setup allowed us to investigate the pleiotropic effects of trisomy, GTE-EGCG treatment and treatment discontinuation at the structural, functional, molecular, and genetic levels in the Ts65Dn Down syndrome mouse model (Fig. 1). We compared wildtype (WT) and Ts65Dn (TS) untreated mice to evaluate trisomy effects, and WT and TS treated mice to evaluate treatment effects.

## STRUCTURAL CHARACTERIZATION

Structurally, we investigated the longitudinal effects of trisomy, treatment, and treatment discontinuation on body weight, skeletal system, and adult brain volume (Fig. 2).

**Figure 2:**
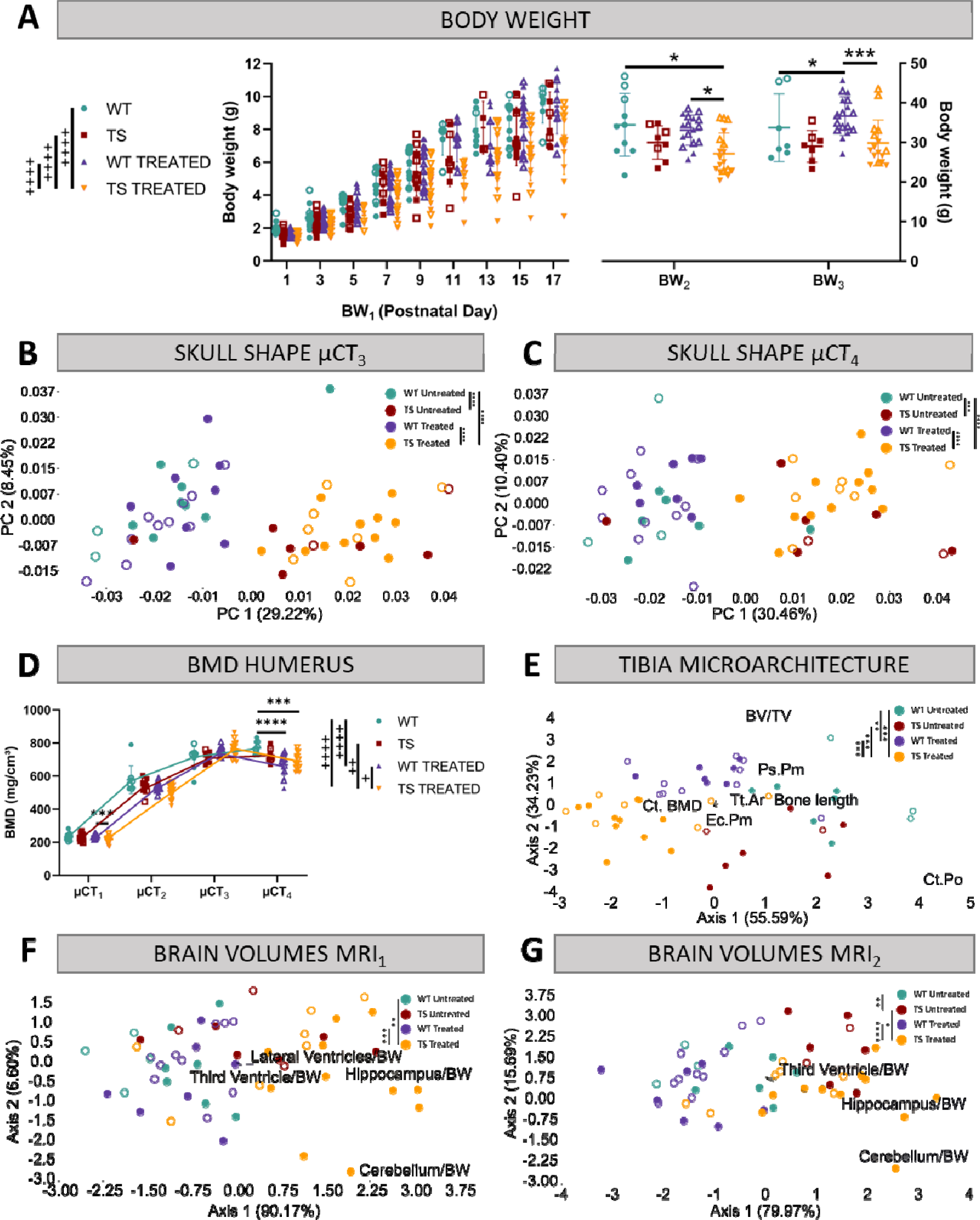
Evaluation of trisomy, prenatal chronic GTE-EGCG treatment and treatment discontinuation effects on structural traits. **(A)** Body weight measurements over postnatal development in untreated and treated WT and TS mice. **(B,C)** Skull shape differences between adult WT and TS mice before (B) and after GTE-EGCG treatment discontinuation (C). Skull shape variation explored by a Principal Component Analysis (PCA) based on the 3D coordinates of landmarks recorded on the surfaces of 3D craniofacial reconstructions from *in vivo* µCT at µCT_3_ (B) and µCT_4_ (C) timepoints. The landmark configuration for each stage is defined in Supplementary Figure S1. Scatter plots are presented along with the morphings associated with the negative and positive extremes of the PC1 axis. Statistical differences between groups were assessed by permutation tests based on the Procrustes distances. **(D)** Bone mineral density of the humerus over postnatal development. The *in vivo* µCT scans of the humerus were used to determine the BMD at µCT_1_, µCT_2_, µCT_3_ and µCT_4_ timepoints. **(E)** Linear discriminant analysis (LDA) based on the results from the tibia microarchitecture tests performed at endpoint (8M), three months after chronic treatment discontinuation. **(F,G)** LDA based on the brain volumes obtained from *in vivo* MRI before (F) and after GTE-EGCG treatment discontinuation (G). The contribution of each variable to separate groups of mice across Axis1 and Axis2 is represented in each LDA as lines pointing in the direction of each axis, with longer lines indicating higher contributions. All data are presented as mean +/-standard deviation. (+) p< 0.05; (++) p < 0.01; (++++) p< 0.0001; Mixed-effects analysis across timepoints; (***) p < 0.001; (****) p< 0.0001; pairwise comparisons. Mice analyzed may differ across stages due to due to uncontrollable technical issues inherent to longitudinal studies, such as scanning failure or mouse death during the experiment but represent overall ontogenetic trajectories. Male mice are indicated with empty symbols. Sample sizes used in each test are provided in Supplementary Table S1.

### Body Weight

First, we investigated mouse body growth by monitoring body weight over development. TS untreated mice tended to present lower body weights than WT untreated mice over development from birth to adulthood, but these genotype differences did not reach significance at any developmental stage (Fig. 2A; *P*_BW1_=0.7630; *P*_BW2_=0.3688; *P_BW3_*=0.1094).

Chronic GTE-EGCG treatment, which started prenatally and was maintained until 5 months, significantly reduced the body growth of TS treated mice from PD1 to PD17 (Fig. 2A at BW_1_) in comparison with WT untreated (*P*<0.0001), WT treated (*P*<0.0001), and TS untreated mice (*P*<0.0001). At adulthood before treatment discontinuation (Fig. 2A at BW_2_), these differences were maintained, but only reached significance when compared to WT untreated (*P*= 0.0322) and WT treated mice (*P*= 0.0103). After three months of treatment discontinuation, both WT and TS treated mice showed a mild increase in body weight, making TS treated mice not significantly different from WT untreated (Fig. 2A; *P_BW3_*=0.9284), and causing WT treated mice to be significantly different from WT untreated (Fig. 2A; *P_BW3_*=0.0259).

### Skeletal Development

Since people with DS show craniofacial abnormalities, reduced bone mineral density (BMD), and altered bone microarchitecture (Carfì et al., 2017; Kao et al., 1992; LaCombe & Roper, 2020; Thomas & Roper, 2021; Vicente et al., 2020), we investigated the effects of genotype, treatment, and treatment discontinuation in these systems by performing *in vivo* µCT scans throughout development and *ex vivo* µCT scans at endpoint as indicated in Figure 1.

#### Craniofacial morphology

We first performed geometric morphometric analysis on the craniofacial 3D models obtained from *in vivo* µCT scans to investigate craniofacial shape throughout development. The results confirmed our previous findings (Llambrich, González-Colom, et al., 2022; Llambrich, González, et al., 2022), showing that TS untreated mice presented a significantly different craniofacial shape than WT untreated mice at µCT_1_ (*P*<0.0001) and µCT_2_ (*P*=0.0011) (Supplementary Table S2). In this study, we further detected that the craniofacial differences induced by genotype persisted until adulthood (Fig. 2B, *P_µCT3_*=0.0009; Fig. 2C*, P_µCT4_*=0.0055).

Prenatal chronic GTE-EGCG treatment only showed a significant effect in the craniofacial shape at PD3, where both WT and TS treated mice were different from their untreated counterparts (*P*_WT_=0.0418; *P*_TS_=0.0005). However, the treatment never rescued the craniofacial shape in TS treated mice, as these mice were significantly different from WT untreated mice at all stages (Fig. 2 B,C; Supplementary Table S2). Discontinuing the treatment for three months did not induce any effects (Fig. 2C), maintaining the craniofacial dysmorphologies already observed at µCT_3_ (Fig. 2B).

#### Bone Mineral Density

The BMD of the humerus was estimated throughout development from the *in vivo* µCT scans. Although we observed that TS untreated mice tended to present lower BMD than WT untreated mice over development and adulthood, the differences did not reach significance at any developmental stage (Fig. 2D, Supplementary Table S3).

Prenatal GTE-EGCG chronic treatment significantly modified the developmental trajectory of humerus BMD in WT and TS treated mice (*P*_WT_<0.0001; *P*_TS_=0.0023), but never rescued the trisomic phenotype, as TS treated mice remained different from WT untreated (*P*<0.0001). At PD3, WT and TS treated mice showed similar levels of BMD as compared to untreated mice (Fig. 2D at µCT_1_) but tended to show lower BMD at PD29 (Fig. 2D at µCT_2_). This tendency for reduced BMD was rescued at adulthood (Fig. 2D at µCT_3_), but after three months of treatment discontinuation, both WT and TS treated groups showed a decline in BMD (Fig. 2D at µCT_4_), which caused WT and TS treated mice to be different than WT untreated control mice (*P*_WT_=0.0001; *P*_TS_=0.0002), suggesting a rebound effect following treatment retrieval.

#### Tibia Development & Microarchitecture

Finally, we complemented our structural skeletal analysis by evaluating the length and microarchitecture of the tibia from *ex vivo* µCT scans at endpoint, as indicated in Figure 1.

To obtain a global view of the genotype and treatment effects, we performed a linear discriminant analysis (LDA) combining the results of all tibia microarchitecture tests into a single analysis that maximized the differences among groups of mice (Fig. 2E). The LDA showed limited overlap between groups, as mice separated by treatment along the first axis, and by genotype along the second axis. These results suggested genotype differences and treatment effects that did not rescue the trisomic phenotype (Fig. 2E). The multivariate pairwise comparisons after one-way PERMANOVA using Mahalanobis distances detected significant differences between all groups of mice except between WT untreated and TS untreated mice (Supplementary Table S4), confirming the treatment effects in both genotypes.

To explore the overall differences detected by the LDA in finer detail, we grouped the results of the tibia microarchitecture tests according to the structural domain they evaluated (cortical bone strength, cortical bone size and trabecular bone), and repeated the multivariate pairwise comparisons (Supplementary Table S4). Even though we did not detect any significant difference between WT untreated and TS untreated mice in any structural domain (Supplementary Table S4), TS untreated mice tended to show specific reductions in cortical thickness, polar moment of inertia, cross sectional area, and bone area (Figure 2-figure supplement 1 C,E,F,G); and the univariate pairwise comparison tests revealed significant differences in the tibia length (Figure 2-figure supplement 1A; *P*=0.0170) and periosteal perimeter (Figure 2-figure supplement 1I; *P*=0.0154).

After GTE-EGCG chronic treatment and discontinuation, WT treated mice were not significantly different than WT untreated mice for any specific structural domain (Supplementary Table S4), but the univariate pairwise tests indicated significantly reduced tibia length (Figure 2-figure supplement 1A; *P*=0.0006) and cortical porosity (Figure 2-figure supplement 1D; *P*=0.0120), together with increased cortical bone mineral density (Figure 2-figure supplement 1B; *P*=0.0120). TS treated mice were significantly different than TS untreated mice for the cortical bone strength and cortical bone size domains (Supplementary Table S4), showing significantly more mineralized (Figure 2-figure supplement 1B; *P*=0.0039), thicker (Figure 2-figure supplement 1C; *P*=0.0400), and less porous (Figure 2-figure supplement 1D; *P*=0.0005) cortical bone. However, they appeared different from WT untreated mice in both cortical domains (Supplementary Table S4) as well as in the cortical BMD (Figure 2-figure supplement 1B; *P*=0.0097) and cortical porosity tests (Figure 2-figure supplement 1D; *P*=0.0107). These results could be explained either by adverse effects of the treatment during development that were persistent after discontinuation, or by a negative effect following treatment withdrawal.

**Figure 2-figure supplement 1.**
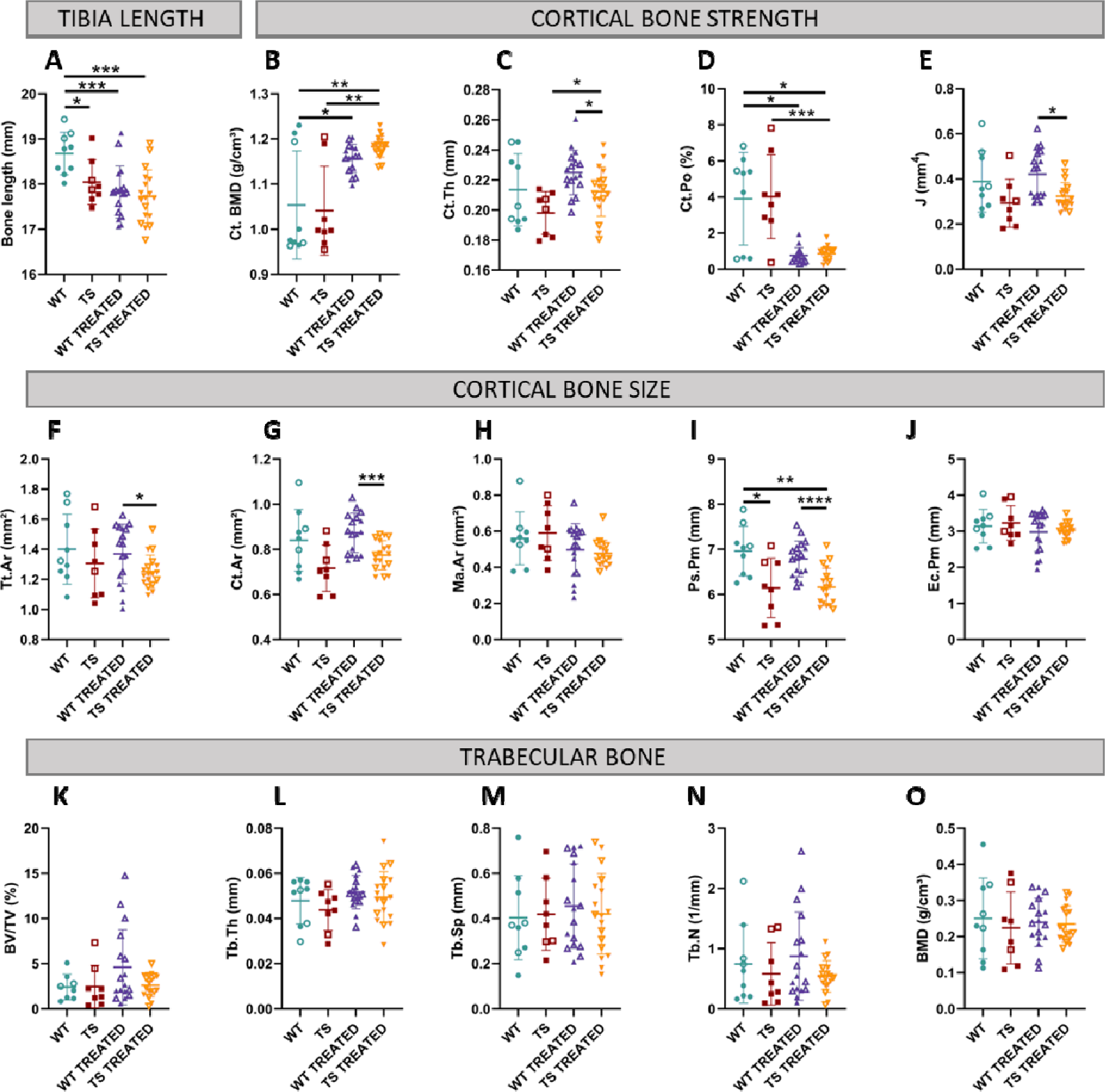
Univariate analysis of tibia microarchitecture in wildtype and trisomic mice at adulthood and effects of prenatal chronic GTE-EGCG treatment and three months of discontinuation. **(A)** The tibia length was measured using a digital caliper at endpoint. The measurements assessing the microarchitecture of the tibia were obtained from *ex vivo* µCT scans and grouped in three categories according to structural bone domains: **(1) Cortical bone strength: (B)** cortical bone mineral density, **(C)** cortical thickness, **(D)** cortical porosity, and **(E)** polar moment of inertia; **(2) Cortical bone size: (F)** cross sectional area, **(G)** bone area, **(H)** medullary area, **(I)** periosteal perimeter, and **(J)** endocortical perimeter. **(3) Trabecular bone: (K)** percentage of bone volume, **(L)** trabecular thickness, **(M)** trabecular separation, **(N)** trabecular number, and **(O)** bone mineral density. Data are presented as mean +/− standard deviation. (*) p < 0.05; (**) p < 0.01; (***) p< 0.001; (****) p< 0.0001; pairwise tests. Male mice are indicated with empty symbols. Sample sizes for each test are provided in Supplementary Table S1.

### Brain Development

Finally, as it was reported that people with DS show altered brain volumes (Aylward et al., 1997; Hamner et al., 2018; Movsas et al., 2016; Patkee et al., 2020; Pinter et al., 2001; Rodrigues et al., 2019; Smigielska-Kuzia et al., 2011), we completed our structural evaluation by assessing the volume of the whole brain, along with the volumes of the hippocampal region, the cerebellum and the ventricles using the *in vivo* MRI scans performed before and after treatment discontinuation (Fig. 1).

The LDA including the volumes of the subcortical brain regions mentioned above showed genotype separation across Axis1 at MRI_1_, which further increased at MRI_2_, with WT mice occupying the negative extreme of Axis1 and TS mice occupying the positive extreme (Fig. 2 F,G). The multivariate pairwise comparisons after one-way PERMANOVA using Mahalanobis distances (Supplementary Tables S5 & S6) confirmed genotype differences only at MRI_2_, when WT untreated mice were significantly different than TS untreated (Fig. 2G; *P*=0.0065). Contrary to previous evidence in Ts65Dn mice (Aldridge et al., 2007; Duchon et al., 2021; Holtzman et al., 1996; Insausti et al., 1998), adult TS untreated mice tended to present larger volumes than WT untreated mice in all brain regions at both MRI_1_ and MRI_2_ (Figure 2-figure supplement 2). The univariate pairwise tests performed at MRI_1_ confirmed that TS mice presented a significantly larger brain (Figure 2-figure supplement 2A, top; *P*=0.0066), hippocampal region (Figure 2-figure supplement 2B, top; *P*=0.0254) and whole ventricles volume (Figure 2-figure supplement 2H, top; *P*=0.0427) than WT untreated mice. At MRI_2_, these differences did no longer reach significance and significant increases emerged in the volumes of the third ventricle (Figure 2-figure supplement 2E, bottom; *P*=0.0479) and fourth ventricle (Figure 2-figure supplement 2F, bottom; *P*=0.0014).

Regarding treatment effects, the LDA at MRI_1_ showed separation between TS treated mice and the rest of the groups (Fig. 2F), and the multivariate pairwise comparisons after one-way PERMANOVA confirmed significant differences between WT untreated and TS treated mice (*P*=0.0047), but not between TS untreated and TS treated mice (*P*=0.4921). Indeed, TS treated mice tended to show larger volumes than TS untreated mice in all brain regions (Figure 2-figure supplement 2), but the univariate pairwise comparison tests only detected a significant difference in the hippocampal region (Figure 2-figure supplement 2B, top; *P*=0.0413). These effects did not rescue the trisomic phenotype, as TS treated mice were significantly different than WT untreated mice in all brain regions (Figure 2-figure supplement 2, top).

After three months of treatment discontinuation at MRI_2_, some TS treated mice tended to occupy the same space as WT untreated mice in the LDA (Fig. 2G), and although the multivariate pairwise comparisons after one-way PERMANOVA did not detect significant differences between WT untreated and TS treated mice (*P*=0.0609), TS treated mice remained significantly different than WT untreated mice in all univariate brain regions except for the volume of the third ventricle (Figure 2-figure supplement 2E, bottom; *P*=0.0562), indicating limited rescuing effects at this last stage after treatment discontinuation.

**Figure 2-figure supplement 2.**
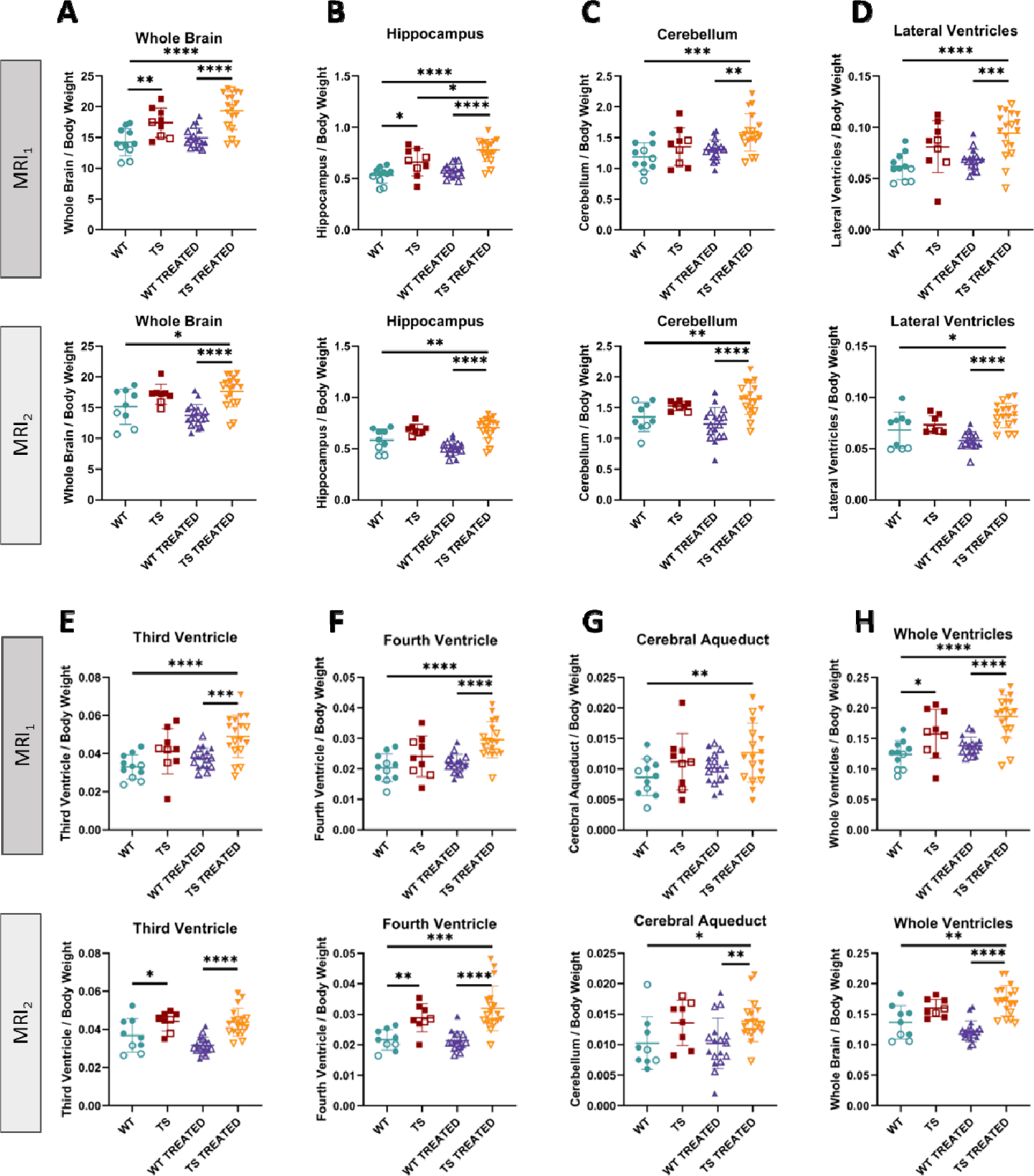
Univariate analysis of brain volumes in wildtype and trisomic mice at adulthood and effects of prenatal chronic GTE-EGCG treatment and its discontinuation. The *in vivo* MRI scans performed at MRS_1_ and MRS_2_ were used to quantify the volumes of **(A)** the whole brain, **(B)** hippocampal region, **(C)** cerebellum, **(D)** lateral ventricles, **(E)** third ventricle, **(F)** the fourth ventricle, **(G)** cerebral aqueduct, and **(H)** whole ventricular system. All volumes were normalized to body weight. For each test, the results at MRI_1_ are presented on top and at MRI_2_ are presented on the bottom. Data are presented as mean +/− standard deviation. (*) p < 0.05; (**) p < 0.01; (***) p< 0.001; (****) p< 0.0001; pairwise tests. Male mice are indicated with empty symbols. Sample sizes for each test are provided in Supplementary Table S1.

## FUNCTIONAL CHARACTERIZATION

After evaluating the effects of trisomy and chronic prenatal GTE-EGCG treatment and its discontinuation at a structural level, we investigated its simultaneous effects at a functional level. We performed a battery of cognitive tests to evaluate early neurodevelopment and adult cognition before and after treatment discontinuation (Fig. 3), at the timepoints specified in Figure 1.

**Figure 3:**
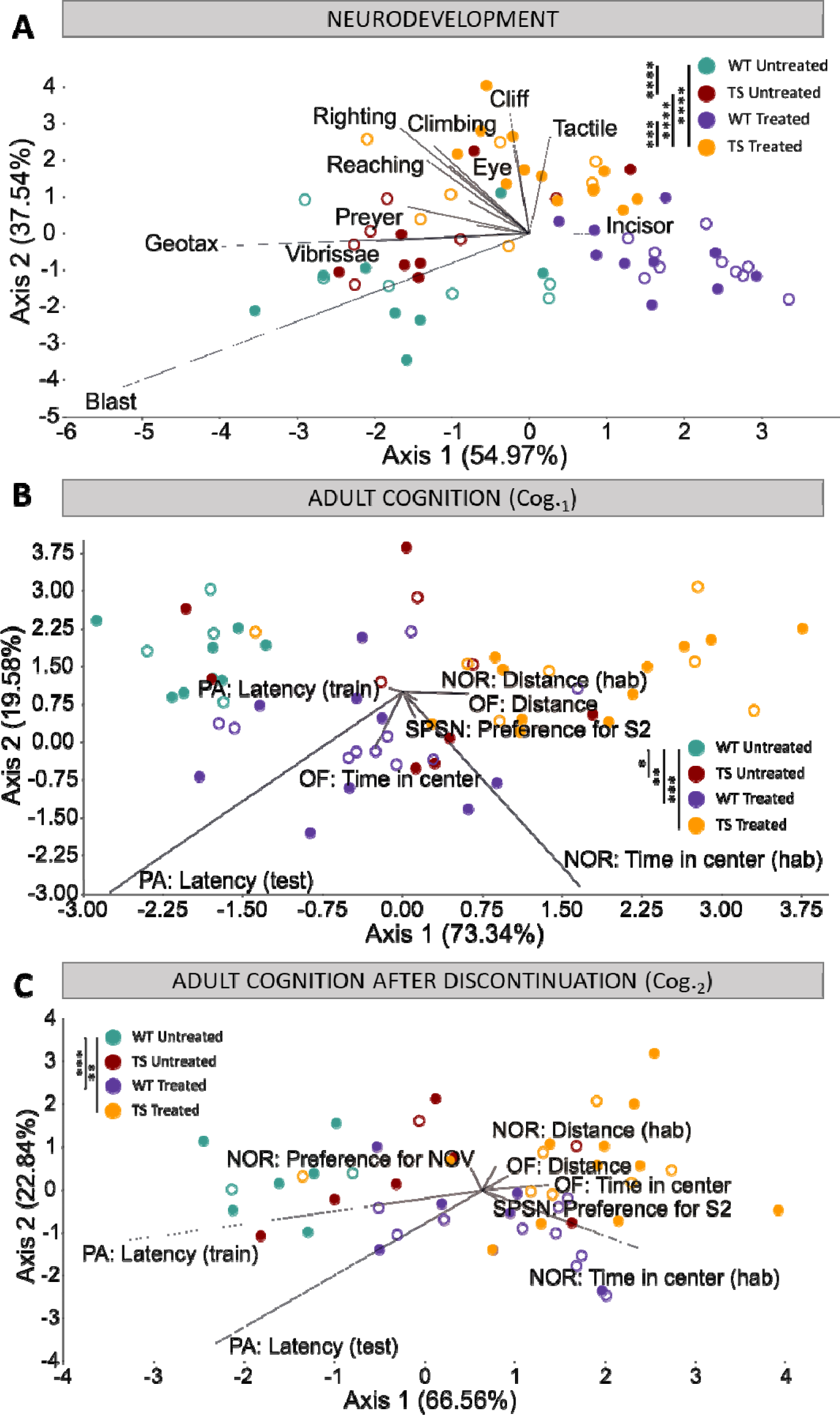
Evaluation of trisomy, prenatal chronic GTE-EGCG treatment and treatment discontinuation effects on cognitive function during development and at adulthood before and after treatment discontinuation. **(A)** Linear discriminant analysis (LDA) based on the results from the neurodevelopmental tests performed from PD1 until PD18. **(B,C)** LDA based on the results from the cognitive tests performed at Cog._1_ (B) and Cog._2_ (C). The contribution of each variable to separate groups of mice across Axis1 and Axis2 is represented in each LDA as lines pointing in the direction of each axis, with longer lines indicating higher contributions. Male mice are indicated with empty symbols. Sample sizes for each test are provided in Supplementary Table S1.

### Neurodevelopment

We first evaluated early brain function and neurodevelopment over the first weeks after birth by performing a battery of neurodevelopmental tests from PD1 until PD18 (Fig. 3A). The LDA with the results from all neurodevelopmental tests combined showed that the treatment was the main differentiating factor among groups of mice, as Axis 1 separated treated and untreated mice (Fig. 3A). Despite some overlap, Axis2 separated WT and TS mice. These results suggested mild genotype differences and treatment effects that did not rescue the trisomic phenotype (Fig. 3A). The multivariate pairwise comparisons after one-way PERMANOVA using Mahalanobis distances (Supplementary Table S7) detected significant differences between TS untreated and WT untreated mice (*P*=0.0001), and between TS treated mice and both TS untreated (*P*=0.0001) and WT untreated mice (*P*=0.0001), confirming these findings.

When the neurodevelopmental tests were grouped into functional domains (developmental landmarks, neuromotor tests and reflexes) (Supplementary Table S7), the multivariate pairwise comparisons only indicated significant differences between WT untreated and TS untreated mice for the reflexes (*P*=0.0423). TS untreated mice tended to be delayed in most individual tests (Figure 3-figure supplement 1), and the univariate pairwise comparison tests confirmed significant delays in the acquisition rate of the vertical climbing (Figure 3-figure supplement 1E; *P*=0.0256) and tactile response (Figure 3-figure supplement 1I; *P*=0.0064), together with the average day of acquisition of the surface righting response (Figure 3-figure supplement 1C; *P*=0.0320). However, we also observed that TS untreated mice tended to be anticipated in the vibrissae placing, blast response, and Preyer reflex (Figure 3-figure supplement 1 J,M,N), even though these results did not reach significance.

Prenatal chronic treatment with GTE-EGCG caused TS treated mice to be significantly different than TS untreated mice for the domains neuromotor tests and reflexes, and WT treated mice to be different from WT untreated mice for the domain reflexes (Supplementary Table S7). The univariate tests indicated that treatment significantly modulated the average day of acquisition and acquisition rate of multiple neurodevelopmental tests in both WT and TS treated mice, causing either a delay or an anticipation in most tests (Figure 3-figure supplement 1). However, the treatment never rescued the trisomic phenotype, as TS treated mice remained significantly different than WT untreated mice for the three functional domains and most of the individual tests (Supplementary Table S7) (Figure 3-figure supplement 1).

Finally, we investigated the mothers’ behavior during 24h at PD8 to evaluate maternal care and observed that, despite being trisomic, all Ts65Dn mothers performed similarly and there was no large intra-group variation for any of the behaviors analyzed, confirming that maternal care did not influence the results of the neurodevelopmental tests (Figure 3-figure supplement 2). Furthermore, no significant differences were detected between treated and untreated mothers in any readout (Figure 3-figure supplement 2), confirming that the GTE-EGCG treatment did not modulate maternal care.

**Figure 3-figure supplement 1.**
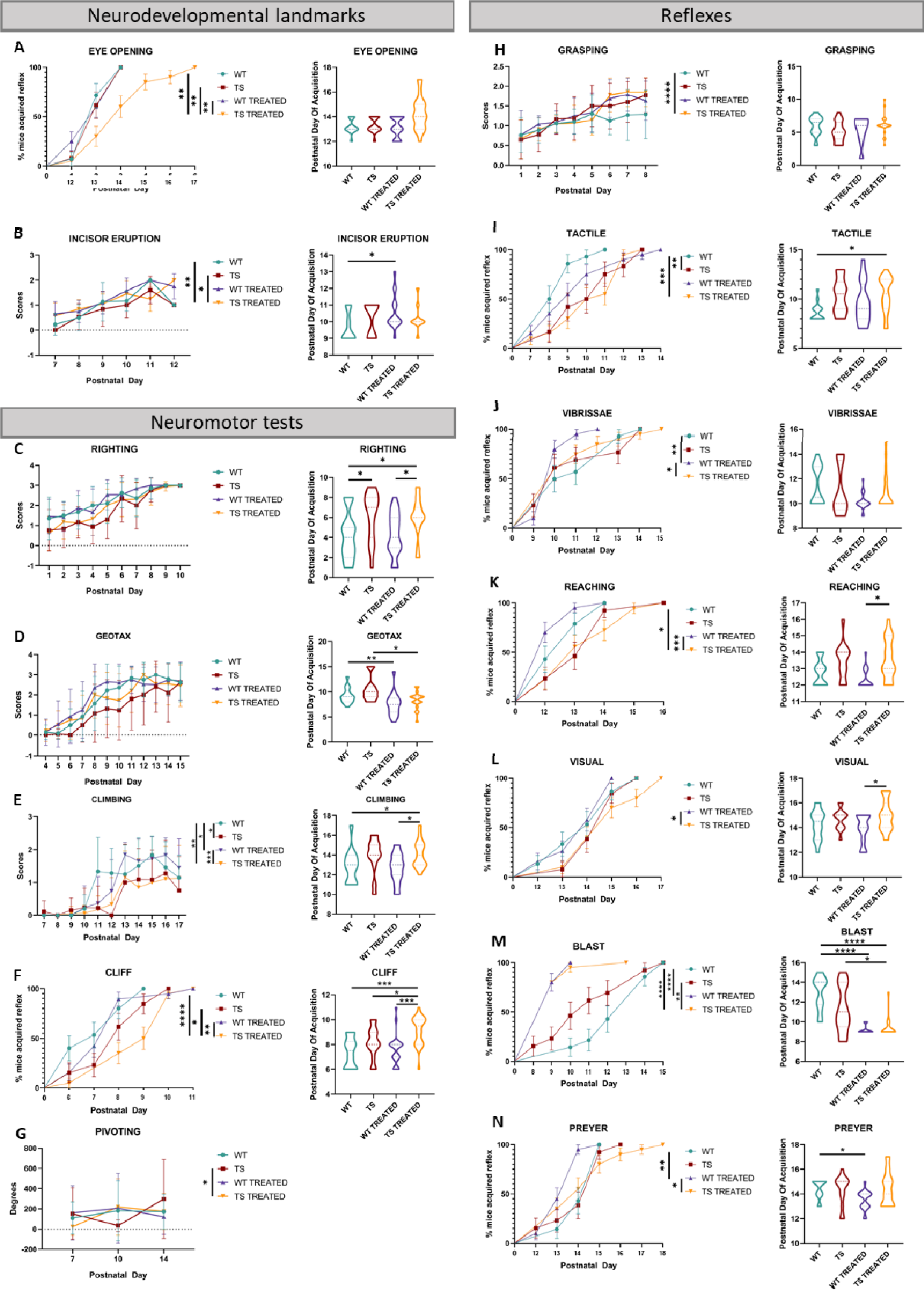
Univariate analysis of early postnatal neurodevelopment in WT and TS mice and effects of pre- and postnatal GTE-EGCG treatment. Neurodevelopment tests were performed from PD1 to PD18 and grouped into three categories according to functional domains. **(1) Neurodevelopmental landmarks: (A)** eye opening, **(B)** incisor eruption, and pinna detachment (results not shown as no significant differences were detected). **(2) Neuromotor development: (C)** surface righting response, **(D)** negative geotaxis, **(E)** vertical climbing, **(F)** cliff drop aversion, **(G)** pivoting, homing and walking (results not shown as no significant differences were detected). **(3) Reflexes: (H)** grasping, **(I)** tactile orientation, **(J)** vibrissae placing, **(K)** reaching response, **(L)** visual placing response, **(M)** blast response and **(N)** Preyer reflex. For each test, developmental rate is presented on the le t as mean +/− standard deviation and average day of successful test acquisition is presented on the right as violin plots. (*) p < 0.05; (**) p < 0.01; (***) p< 0.001; (****) p< 0.0001. Sample sizes for each test are provided in Supplementary Table S1.

**Figure 3-figure supplement 2.**
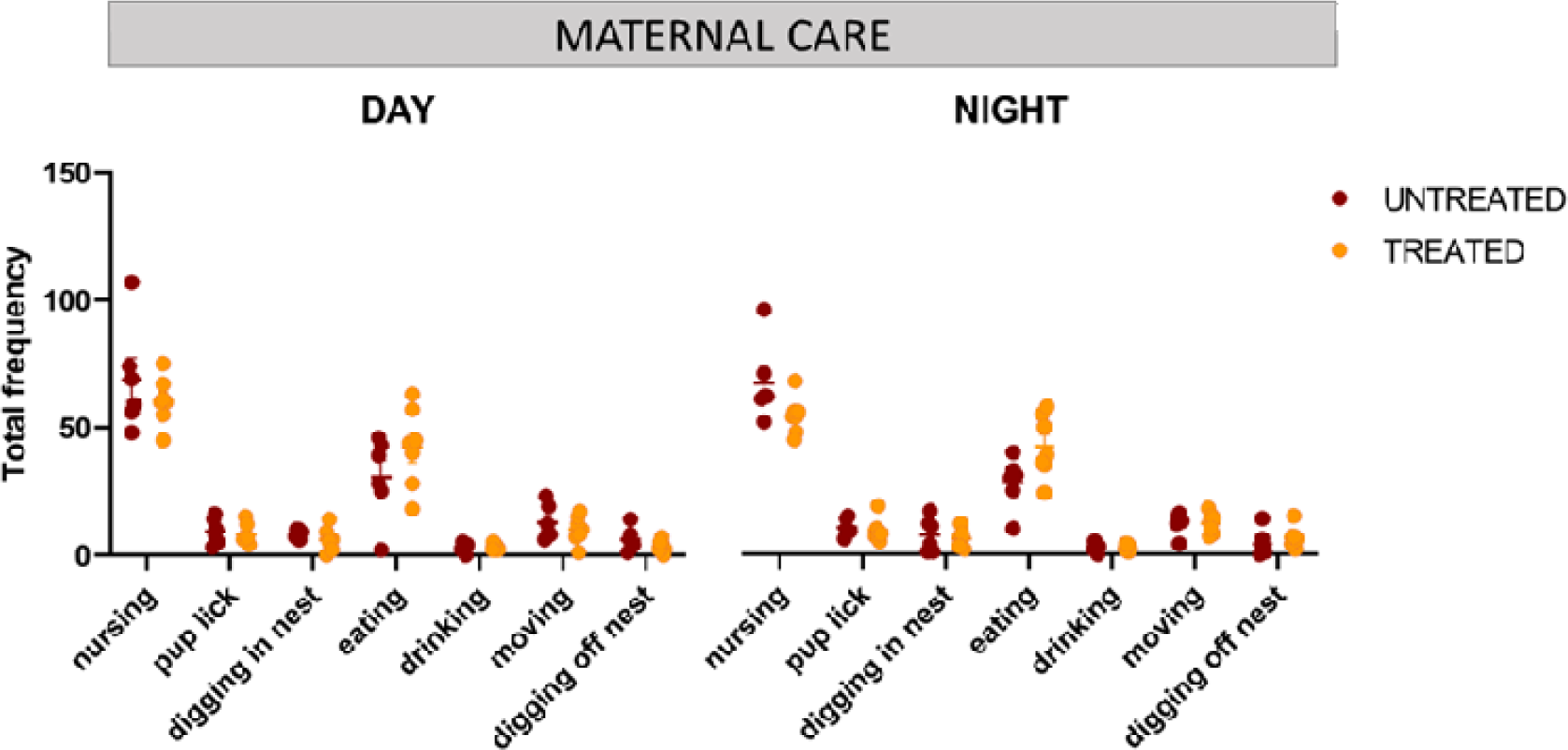
Maternal care evaluation and effects of GTE-EGCG treatment. Frequency of nursing, pup grooming, digging in nest, eating, drinking, moving, and digging off nest behaviors scored manually fro 24h videotapes at PD8. Sample sizes for each test are provided in Supplementary Table S1.

### Adult Cognition

Then, we evaluated adult cognitive performance by means of open field (OF), elevated plus maze (EPM), sociability/preference for social novelty (SPSN), novel object recognition (NOR) and passive avoidance (PA) testing at two timepoints, Cog._1_ and Cog._2_, before and after treatment discontinuation (Fig. 3 B,C), as specified in Figure 1.

The LDA with the results from all tests combined showed minor overlap between groups at both Cog._1_ and Cog._2_ (Fig. 3 B,C), with WT untreated mice occupying the negative extreme of Axis1, WT treated and TS untreated mice presenting an intermediate position, and TS treated mice falling on the positive extreme of Axis1, opposite to WT untreated mice (Fig. 3 B,C). These results suggested genotype differences between WT and TS untreated mice that were confirmed as significant by the multivariate pairwise comparisons after one-way PERMANOVA at Cog._1_ (Supplementary Table S8; *P*=0.0125) but not at Cog._2_ (Supplementary Table S9; *P*=0.4596).

When the cognitive tests were grouped into cognitive domains (anxiety, arousal, and memory), the multivariate pairwise comparisons only detected a significant difference between WT untreated and TS untreated mice for the arousal tests at Cog._1_ (*P*=0.0028) (Supplementary Tables S8 and S9). Indeed, TS untreated mice showed a tendency to cover longer distances at higher speed in all arousal tests when compared with WT untreated mice (Figure 3-figure supplement 3 E-J). These differences reached significance for the distance and speed during OF, as well as for the distance and speed during NOR at Cog._1_ (Figure 3-figure supplement 3 E,F,I,J, top). TS untreated mice also tended to show increased exploratory and risk-taking behavior, as they usually presented a tendency of increased time spent in the center of the arena during habituation for NOR at Cog._1_ (Figure 3-figure supplement 3C, top), increased percentage of time spent in the open arm during EPM at Cog._1_ (Figure 3-figure supplement 3D, top), increased time in the center during OF at Cog._2_ (Figure 3-figure supplement 3A, bottom), and reduced time to enter the dark chamber during the training session of PA at Cog._1_ (Figure 3-figure supplement 3M, top), although these differences did not reach statistical significance. Finally, TS untreated mice showed impaired memory robustness and formation at the latest stage, with significantly less preference for the novel object during NOR when compared with WT untreated mice (Figure 3-figure supplement 3K, bottom; *P*=0.0257), and a tendency for reduced latency to enter the dark chamber during training and testing sessions for PA at Cog._2_ (Figure 3-figure supplement 3 M,N, bottom).

Regarding treatment effects, the results from the LDA at Cog._1_ suggested that the prenatal chronic treatment induced changes in both WT and TS treated mice (Fig. 3 B) that exacerbated the differences in TS treated mice and induced cognitive changes in WT treated mice that resulted in a cognitive performance similar to TS untreated mice (Fig. 3 B). Indeed, WT treated mice were significantly different than WT untreated mice when all cognitive tests were considered together as well as for the anxiety and arousal domains (Supplementary Table S8). In the case of TS treated mice, the multivariate pairwise comparisons revealed significant differences with WT untreated mice when considering all groups together as well as for the anxiety and arousal domains (Supplementary Table S8), confirming the lack of rescuing treatment effects. Both WT and TS treated mice showed significantly increased time in the center during NOR, and increased time in the open arm during EPM at Cog._1_ when compared with WT untreated mice (Figure 3-figure supplement 3 C,D, top), indicating increased exploratory and risk-taking behavior. Similarly, both WT and TS treated mice showed increased locomotor activity, as they covered significantly more distance at more speed than WT untreated mice for the distance and speed during OF, as well as the distance and speed during NOR at Cog._1_ (Figure 3-figure supplement 3 E,F,I,J, top). However, in the memory tests (Figure 3-figure supplement 3 K-N), the pairwise comparison tests revealed only a significant difference between TS untreated and TS treated mice for the preference for subject 2 (S2) at Cog._1_, where treated mice had greater preference for the novel stranger (Figure 3-figure supplement 3L, top; *P*=0.0137).

After treatment discontinuation at Cog._2_, the LDA (Fig.3C) presented a similar pattern as the LDA at Cog._1_ (Fig.3B), suggesting that discontinuing the treatment for one month did not substantially change the cognitive patterns induced after chronic GTE-EGCG treatment. However, new significant differences emerged between WT untreated and both WT treated and TS treated mice when evaluating the memory domain (Supplementary Table S9), suggesting that the treatment discontinuation altered the memory of both WT and TS mice. Furthermore, TS treated mice remained significantly different than WT untreated mice for all tests combined and all cognitive domains (Supplementary Table S9), confirming that the effects of treatment discontinuation did not rescue the trisomic cognitive phenotype. Indeed, significant differences between WT untreated and TS treated mice remained for the percentage of time spent in the open arm during EPM at Cog._2_ (Figure 3-figure supplement 3D, bottom; *P*=0.0476), and new significant differences emerged between WT untreated mice and both WT treated and TS treated mice for the time spent in the center and in the periphery during OF (Figure 3-figure supplement 3 A,B, bottom), indicating that treated mice still presented a more explorative and risk-taking behavior. A similar trend was observed for the arousal tests, as even though no significant differences were detected between TS treated and TS untreated mice (Figure 3-figure supplement 3 E-J, bottom), both treated groups showed hyperactivity as the univariate pairwise comparison tests confirmed significant differences between TS treated mice and WT untreated mice for all arousal tests (Figure 3-figure supplement 3 E-J, bottom); and significant differences were detected between WT treated and WT untreated mice for the distance and speed during SPSN and the distance and speed during NOR (Figure 3-figure supplement 3 G-J, bottom). In the memory tests, TS treated mice were significantly different than TS untreated (*P*=0.0046) but not than WT untreated mice for the preference for the novel object during NOR (Figure 3-figure supplement 3K, bottom; *P*=0.3566), which could suggest a beneficial treatment effect. Contrary to the other behavioral tests, the results for PA testing at Cog._2_ are influenced by the testing at Cog._1_ as mice may present a robust contextual fear memory trace created during the testing at Cog._1_, preventing them from entering the dark chamber even in the training phase of the test at Cog._2_. Indeed, WT untreated mice did not enter the dark box during training at Cog._2_, while both WT and TS treated mice showed a reduced tendency to enter the dark chamber which reached significance when comparing TS treated mice with WT untreated mice (Figure 3-figure supplement 3M, bottom; *P*<0.0001), indicating less robust contextual memory retention. After re-exposure to the fear conditioning in the training phase of PA at Cog._2_, all WT mice avoided the dark chamber while a few TS treated mice still showed a reduced tendency to enter the dark chamber (Figure 3-figure supplement 3N, bottom), reflecting altered memory robustness.

**Figure 3-figure supplement 3.**
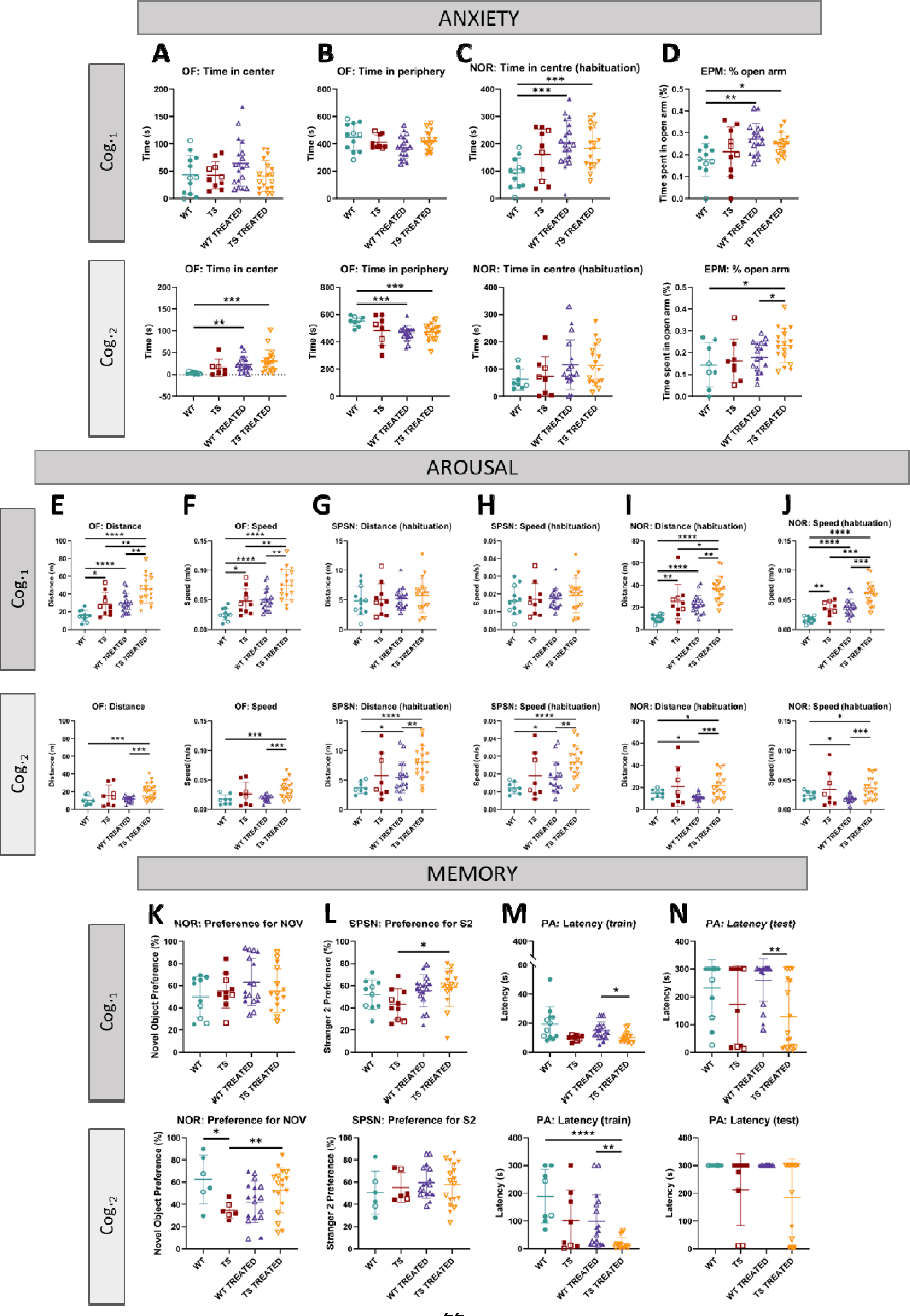
Univariate analysis of adult cognition in WT and TS mice and effects of prenatal chronic GTE-EGCG treatment and its discontinuation. Results for open field (OF), novel object recognition (NOR), elevated plus maze (EPM), sociability/preference for social novelty (SPSN) and passive avoidance (PA) tests performed at Cog._1_ (top) and Cog._2_ (bottom) were grouped into three categories according to functional domains. **(1) Anxiety tests: (A)** time in center during OF, **(B)** time in periphery during OF, **(C)** time in center during habituation for NOR, and **(D)** percentage of time in open arm during EPM. **(2) Arousal tests: (E)** total distance covered during OF, **(F)** mean speed during OF, **(G)** total distance covered during SPSN, **(H)** mean speed during SPSN, **(I)** total distance covered during habituation for NOR, and **(J)** mean speed during habituation for NOR. **(3) Memory tests: (K)** preference for novel object during testing for NOR, **(L)** preference for second stranger during SNS2, **(M)** latency to enter dark chamber during training for PA, and **(N)** latency to enter dark chamber during testing for PA. For each test, the results at Cog._1_ are presented on top and the results at Cog._2_ are presented on the bottom. Data are presented as mean +/− standard deviation. (*) p < 0.05; (**) p< 0.001; (***) p< 0.001; (****) p< 0.0001. Male mice are indicated with empty symbols. Sample sizes for each test are provided in Supplementary Table S1.

## MOLECULAR AND GENETIC CHARACTERIZATION

Finally, to complement the structural and functional evaluation and to obtain a complete holistic characterization of the alterations associated with DS, we performed *in vivo* MRS and *ex vivo* RNAseq at the timepoints specified in Figure 1 to investigate changes in hippocampal region metabolite concentration and cerebellar gene expression respectively (Fig.4).

**Figure 4.**
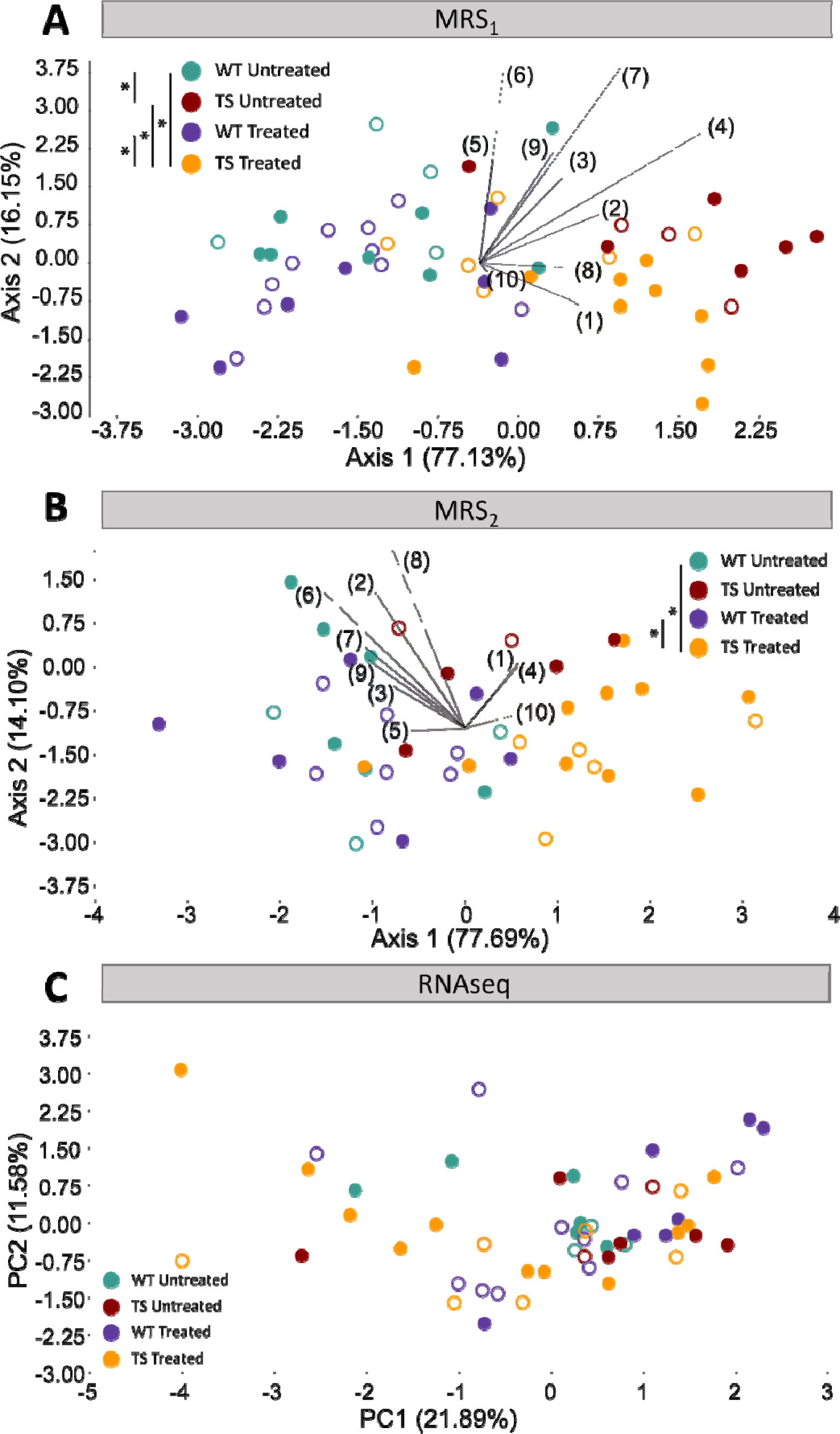
Evaluation of trisomy, prenatal chronic GTE-EGCG treatment and treatment discontinuation effects on molecular and gene expression parameters. **(A,B)** Linear discriminant analysis (LDA) based on the relative integrals of 10 spectral regions obtained from MRS performed in the hippocampal region at MRS_1_ (A) and MRS_2_ (B). The contribution of each variable to separate groups of mice across Axis1 and Axis2 is represented in each LDA as lines pointing in the direction of each axis, with longer lines indicating higher contributions. **(C)** Principal component analysis (PCA) based on the normalized expression of the 125 triplicated genes obtained from RNAseq at 8M. Male mice are indicated with empty symbols. Sample sizes for each test are provided in Supplementary Table S1.

### Hippocampal Region Metabolites

We performed *in vivo* MRS on a voxel placed in the hippocampal region at two timepoints, MRS_1_ and MRS_2_, as specified in Figure 1.

We first analyzed global differences in the metabolite spectra by performing an LDA containing the relative integrals of 10 spectral regions between 0.0 ppm and 4.3 ppm (Supplementary Figure S2 and Supplementary Table S10). The LDA at MRS_1_ separated WT untreated and TS untreated mice across Axis1 (Fig. 4A), and the multivariate pairwise comparisons after one-way PERMANOVA using Mahalanobis distances confirmed significant genotype differences (Supplementary Table S11; *P*=0.0136). After treatment discontinuation at MRS_2_, the LDA showed milder genotype separation (Fig. 4B) and the multivariate pairwise comparisons did not detect any significant difference between WT untreated and TS untreated mice (Supplementary Table S12; *P*=0.5790), confirming these findings.

When evaluating the concentrations of specific main metabolites like N-acetylaspartate (NAA), creatine + phosphocreatine (Cr+PCr), choline + phosphocholine + glycerophosphocoline (tCho), myo-inositol (myo-Inos) and taurine (Tau) (Figure 4-figure supplement 1), TS untreated mice tended to show lower levels of NAA and tCho at MRS_1_ when compared with WT untreated mice (Figure 4-figure supplement 1 A,C), but the univariate pairwise comparison tests only revealed a significant increase in the concentration of Cr+PCr (Figure 4-figure supplement 1B, top; *P*=0.0235). At MRS_2_, the same pattern was observed but the significant differences between WT untreated and TS untreated mice disappeared (Figure 4-figure supplement 1, bottom).

Regarding treatment effects, the LDA at MRS_1_ separated treated and untreated mice across Axis2 (Fig. 4A), suggesting treatment effects that did not rescue the trisomic phenotype. The multivariate pairwise comparisons after one-way PERMANOVA (Supplementary Table S11) showed significant differences between TS treated and both WT untreated (*P*=0.0496) and TS untreated mice (*P*=0.0361), confirming these findings. When evaluating the specific concentrations of NAA, Cr+PCr, tCho, myo-Inos and Tau, both WT treated and TS treated mice tended to show increased levels of Cr+PCr, which reached significance when comparing TS treated mice with WT untreated mice (Figure 4-figure supplement 1B, top; *P*= 0.0317). Conversely, both WT and TS treated mice tended to show reduced levels of myo-inositol and tCho (Figure 4-figure supplement 1D, top), causing TS treated mice to have significantly reduced tCho when compared with WT untreated mice (Figure 4-figure supplement 1C, top; *P*= 0.0296).

After treatment discontinuation at MRS_2_, the LDA showed milder treatment separation across Axis2 (Fig. 4B) and the multivariate pairwise comparisons did not detect any significant difference between TS untreated and TS treated mice (Supplementary Table S12; *P*=0.2111). However, the differences between TS treated and WT untreated mice remained significant (*P*=0.0347), indicating that discontinuing the treatment for three months did not rescue the trisomic phenotype. When evaluating the specific concentrations of NAA, Cr+PCr, tCho, myo-Inos and Tau, the same pattern was observed and the significant differences between TS treated and WT untreated mice disappeared (Figure 4-figure supplement 1, bottom). However, significant differences emerged between WT untreated mice and WT treated mice for the concentration of tCho and myo-inositol (Figure 4-figure supplement 1 C,D, bottom; *P_tCho_* = 0.0479, *P_myo-inositol_*= 0.0261).

**Figure 4-figure supplement 1.**
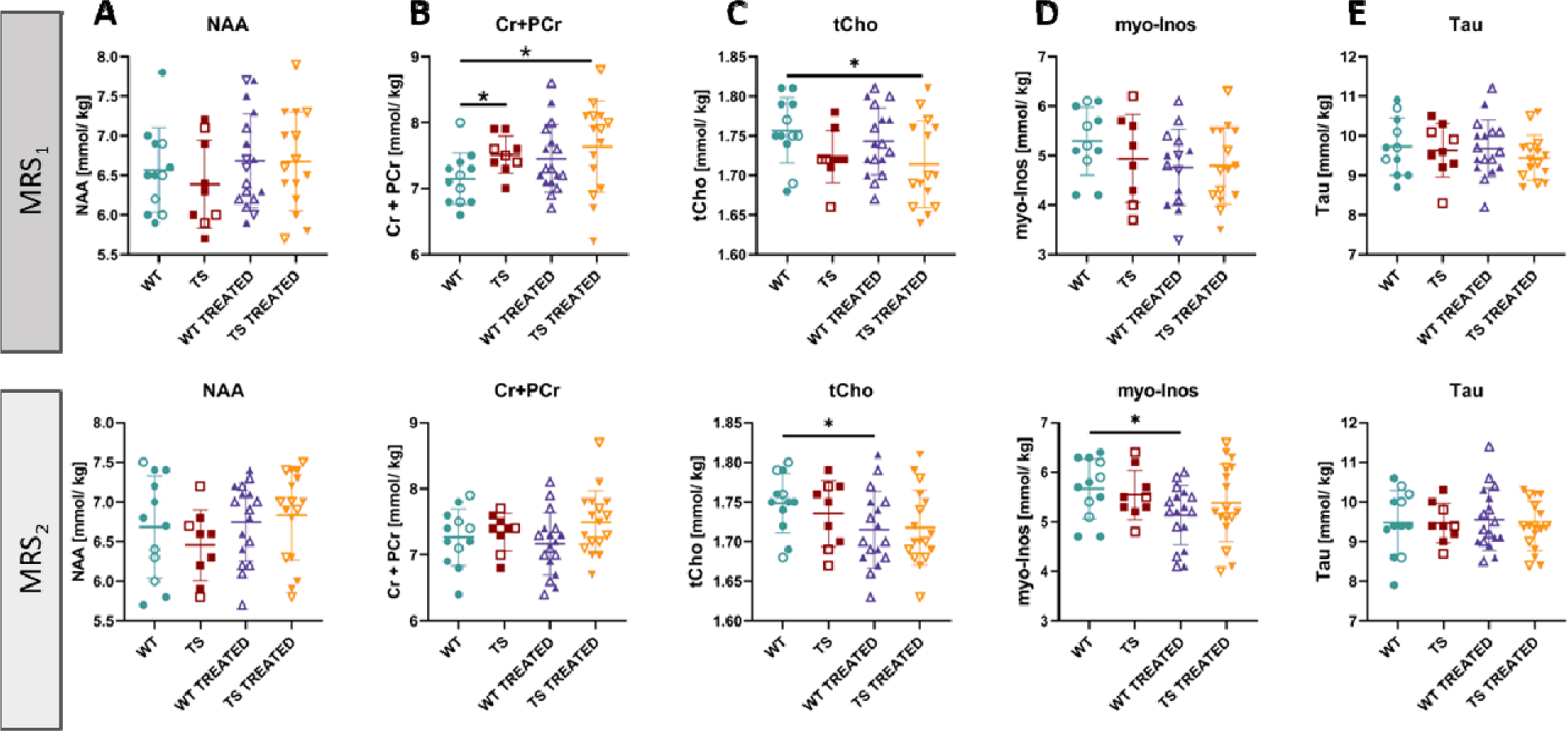
Univariate analysis of hippocampal metabolites in wildtype and trisomic mice at adulthood and effects of prenatal chronic GTE-EGCG treatment and its discontinuation. The *in vivo* MRS scans performed in the hippocampal region at MRS_1_ (top) and MRS_2_ (bottom) were used to quantify the concentrations of **(A)** N-acetylaspartate (NAA), **(B)** creatine + phosphocreatine (Cr+PCr), **(C)** choline + phosphocholine + GPC (tCho), **(D)** myo-Inositol (myo-Inos) and **(E)** taurine (Tau). For each test, the results at MRS_1_ are presented on top and results at MRS_2_ are presented on the bottom. Data are presented as mean +/− standard deviation. (*) p < 0.05; pairwise tests. Male mice are indicated with empty symbols. Sample sizes for each test are provided in Supplementary Table S1.

### Gene Expression

Finally, since people with DS show impaired motor skills (Quinzi et al., 2022), we investigated the effects of genotype and chronic treatment after three months of treatment discontinuation in the cerebellar gene expression by performing RNAseq at endpoint (8M).

We first performed a PCA using the normalized expression of the 125 genes that are triplicated in the Ts65Dn mouse model and included in our dataset. The PCA showed major overlap between all groups, even though TS treated mice showed a larger range of variation (Fig. 4C). This result suggested that the differences in gene expression of the triplicated genes were minimal and that GTE-EGCG prenatal chronic treatment and three months of discontinuation did not induce major permanent effects when considering all triplicated genes. The multivariate pairwise comparison tests after one-way PERMANOVA using Euclidean distances did not detect any significant difference between groups, confirming these findings (Supplementary Table S13).

Then, we analyzed the differentially expressed genes (DEGs) between each pair of experimental groups across the entire transcriptome (Supplementary Table S14). When analyzing the entire transcriptome, we only detected 24 DEGs between WT untreated and TS untreated mice even when applying a loose p-adjusted value of 0.1, indicating small differences in cerebellar gene expression (Supplementary Table S14). Interestingly, we detected 12 DEGs between WT untreated and TS treated mice, and only 9 of these genes were also found to be differentially expressed between WT untreated and TS untreated mice, which could indicate mild treatment effects in TS mice (Supplementary Table S14). However, we did not detect any DEGs between TS untreated and TS treated mice, and only 5 DEGs were detected between WT untreated and WT treated mice, indicating that either the prenatal chronic GTE-EGCG treatment did not largely modulate gene expression or that the treatment had transient effects that were mostly not detected after three months of treatment discontinuation and were not rescuing the trisomic phenotype (Supplementary Table S14).

## DISCUSSION

We investigated for the first time the simultaneous pleiotropic effects of trisomy and a chronic prenatal and postnatal treatment with GTE-EGCG on several organ systems and levels throughout development in the Ts65Dn mouse model for DS. Our longitudinal and holistic approach overcomes a main limitation of previous research, in which systems were analyzed independently at single time points using different experimental setups and delivered disparate and contradictory results that could not be generalized. Here, we followed the simultaneous development of structural, functional, molecular, and genetic systems on Ts65Dn mice and provided comparable results of genotype and prenatal chronic GTE-EGCG treatment effects before and after treatment discontinuation.

### Detailed comparison of genotype and GTE-EGCG treatment effects across studies

We here summarized and compared our results with previous studies by domain to discuss whether the Ts65Dn mouse model was representative of DS in the human condition, and whether GTE-EGCG treatment produced any effect that could be translated into clinical practice.

#### Structural domain

Starting with body size, we observed that TS mice tended to weigh less than WT mice. However, as in a recent study (Tallino et al., 2022), this difference did not achieve statistical significance, and contrasts with previous studies reporting significantly smaller weight of TS mice (Costa et al., 2010; Goodlett et al., 2020; Heinen et al., 2012; Jamal et al., 2022; Roper et al., 2006). The prenatal chronic GTE-EGCG treatment further reduced the body weight of TS treated mice from PD1 to PD17, but not at adulthood, which is consistent with some studies (Goodlett et al., 2020; Xicota et al., 2020), but not with others (Jamal et al., 2022; Noll et al., 2022). Moreover, we observed that WT treated mice showed normal body weight during development, but increased weight after treatment discontinuation, suggesting a possible rebound effect after a prolonged time of treatment administration.

Regarding skeletal morphology, we detected craniofacial alterations that were consistent with previous findings and replicated the human condition (Llambrich, González-Colom, et al., 2022; Llambrich, González, et al., 2022; McElyea et al., 2016; Suri et al., 2010). We also detected modulatory treatment effects on the craniofacial morphology in treated WT and TS mice at PD3, but these did not remain significant at adulthood, which was an unexpected result considering that our previous findings indicated that GTE-EGCG had modulatory effects from birth that were exacerbated later in development at PD29 (Llambrich, González-Colom, et al., 2022; Llambrich, González, et al., 2022).

Our multivariate analyses did not detect global tibia microarchitecture alterations associated with genotype and the univariate tests only showed a significant reduction in the length and periosteal perimeter of the proximal tibia, not showing other skeletal alterations previously described (Abeysekera et al., 2016; J. D. Blazek et al., 2015; Blazek et al., 2011; Joshua D.Blazek et al., 2015; Carfì et al., 2017; LaCombe & Roper, 2020; Llambrich, González-Colom, et al., 2022; Thomas et al., 2020; Thomas et al., 2021). The prenatal chronic treatment and three months of discontinuation did not modulate the trabecular bone and caused general but mild adverse effects in the cortical bone of WT and TS mice, not replicating the modulatory effects of green tea polyphenols previously observed on bone microarchitecture (J. D. Blazek et al., 2015; Goodlett et al., 2020; Huang et al., 2020; Jamal et al., 2022). Moreover, the GTE-EGCG treatment modified the developmental trajectory of humerus BMD in both WT and TS treated mice with disparate effects throughout development. These results did not match our previous findings (Llambrich, González-Colom, et al., 2022) and highlight the variability of treatment responses depending on developmental timing.

In the brain, we detected increased ventricular volumes, which is consistent with the trend observed in our previous study at PD29 (Llambrich, González, et al., 2022) and the ventriculomegaly typically observed in humans and mouse models for DS (Ishihara et al., 2010; Movsas et al., 2016; Pearlson et al., 1998; Raveau et al., 2017). However, we also detected increased hippocampal, cerebellar, and whole brain volumes in TS untreated mice that do not match clinical reports indicating reduced brain, hippocampal and cerebellar volumes in humans with DS (Aylward et al., 1997; Hamner et al., 2018; Patkee et al., 2020; Pearlson et al., 1998; Pinter et al., 2001; Rodrigues et al., 2019; Smigielska-Kuzia et al., 2011), and other preclinical studies that did not find brain volume differences in Ts65Dn mice (Aldridge et al., 2007; Duchon et al., 2021; Holtzman et al., 1996; Insausti et al., 1998). The prenatal chronic treatment did not have major effects, but generally increased the volume of all brain regions in TS treated mice, which is contrary to a previous study indicating that administering a green tea infusion corresponding to 0.6–1 mg EGCG per day from mating until adulthood significantly reduced the brain weight and volume in YACtg152F7 mice (Guedj et al., 2009).

#### Functional domain

Regarding cognitive function, our results indicated that TS mice showed cognitive delay during early stages of development. As suggested in clinical investigations (Locatelli et al., 2021; J. Luis Olmos-Serrano et al., 2016), the cognitive alterations continued with development, and adult TS mice showed hyperactivity, impaired memory robustness, and more explorative behavior, in line with previous reports (Aziz et al., 2018; Costa et al., 2010; Coussons-Read & Crnic, 1996; Dierssen et al., 2002; Escorihuela et al., 1995; Escorihuela et al., 1998; Holtzman et al., 1996; Llambrich, González, et al., 2022; J. Luis Olmos-Serrano et al., 2016). Regarding treatment effects, the combined results from all tests suggested that even though the treatment could have had a positive effect reducing the anxiety of treated mice, it likely altered memory robustness and induced a more hyperactive, explorative, and risk-taking behavior. After treatment discontinuation, when the tests were repeated, all groups of mice presented less locomotor activity and explorative behavior. It is not unlikely that this reduction in activity was due to the stress and anxiety induced by the manipulation and testing during the first round of cognitive evaluation. However, TS untreated mice and both WT and TS treated mice still tended to show increased locomotor activity and explorative behavior compared with WT untreated mice, suggesting that the prenatal chronic treatment could have had permanent adverse effects after one month of discontinuation, or could have reduced the anxiety produced by the first round of testing while mice were treated. These results are in line with previous articles indicating no or negative effects of EGCG on cognition in humans and mice (Cieuta-Walti et al., 2022; Goodlett et al., 2020; Stringer et al., 2015; Stringer et al., 2017), but are contrary to other articles showing rescuing effects (Catuara-Solarz et al., 2016; de la Torre et al., 2016; De la Torre et al., 2014; Souchet et al., 2019; Stagni et al., 2016; Yin et al., 2017).

#### Molecular domain

Regarding hippocampal metabolite concentration, we observed differences between untreated WT and TS mice when evaluating the entire spectra using the LDA, which reflects the analysis of the whole MR spectra rather than selected metabolites, but we only detected a specific significant difference in the levels of Cr+PCr at MRS_1_, suggesting that the genotype produced mild general molecular differences rather than large specific disruptions in any metabolite. Our results did not reveal any difference in NAA or myo-inositol, which is consistent with a human study showing no differences in the levels of NAA in the hippocampal region of adults with DS without dementia (Lamar et al., 2011), but do not replicate the results from another human study indicating increased myo-inositol levels (Beacher et al., 2005). Other studies in humans with DS reported decreased NAA levels relative to creatine and myo-inositol in other brain regions, such as the frontal lobes and posterior cingulate cortex (Lin et al., 2016; Smigielska-Kuzia & Sobaniec, 2007). Studies performed in the Ts65Dn model indicated reduced NAA levels relative to creatine in the hippocampal region (Huang et al., 2000; Même, 2014), which is contrary to our findings, but another study indicated no differences in NAA absolute concentrations (Santin et al., 2014). Furthermore, all these mouse studies indicated increased values of inositol or myo-inositol, which we did not detect in this study. These results reflect the large variability of phenotypes associated with DS depending on the developmental stage and region analyzed, as well as the disparity in experimental setups and data analyses between studies, as some studies reported the absolute metabolite concentrations while others reported their relative values normalized to creatine, which could explain the discrepancy among some of the results.

Regarding treatment effects, we detected significant differences between TS treated mice and both TS untreated and WT untreated mice when analyzing the entire spectra using the LDA, but no significant differences between TS untreated and TS treated mice in any specific metabolite. However, TS treated mice presented significantly increased Cr+PCr and reduced tCho levels as compared with WT untreated mice. After treatment discontinuation, these differences were no longer significant, but general differences remained in the LDA between TS treated and WT untreated mice. These results indicated that prenatal chronic GTE-EGCG treatment had mild modulating effects altering the entire hippocampal metabolite profile in TS mice.

#### Genetic domain

Regarding cerebellar gene expression, we did not detect large differences in the PCA that compared the expression of triplicated genes across groups of mice. Even though these genes presented an extra copy number, their expression was not significantly altered, which is consistent with previous articles (Aït Yahya-Graison et al., 2007; Lyle et al., 2004; J. L. Olmos-Serrano et al., 2016; Saran et al., 2003).

However, we only detected 24 DEGs between untreated WT and TS mice when analyzing the entire genome, which is a lower number in comparison to other studies (Aziz et al., 2018; Chrast et al., 2000; De Toma et al., 2021; J. L. Olmos-Serrano et al., 2016; Saran et al., 2003; Vilardell et al., 2011). Contrary to previous findings indicating a transcriptome-wide deregulation (De Toma et al., 2021; Letourneau et al., 2014; Vilardell et al., 2011), in our study all the 24 DEGs overexpressed in TS untreated mice were mapped to the mouse chromosomes (Mmu) 16 or 17, except for *Ccdc23,* which mapped to Mmu4. Within these 24 DEGs we detected genes typically related with DS and cognitive disability, such as *App*, *Dyrk1A*, *Dscr3*, *Synj1*, *Ttc3*, *Hmgn1*, *Brwd1* and *Usp16*; but we did not detect other genes such as *Sod1, Sim2, Dscam, Rcan1, Olig1, Olig2* or *Kcnj6* (Aziz et al., 2018; Chrast et al., 2000; De Toma et al., 2021; Ruparelia et al., 2012; Vilardell et al., 2011). These discrepancies with previous results could be explained by methodological reasons, as previous studies investigated other tissues, pooled data from different mouse models, contained only male mice, investigated different developmental timepoints, or used different RNA quantification techniques.

Even though the prenatal chronic GTE-EGCG treatment used in this study was administered continuously from embryonic day 9 until adulthood, the treatment did not cause any large permanent effects in the gene expression of adult TS mice after three months of discontinuation, as we did not detect any DEG between TS untreated and TS treated mice. These results suggest that even though the treatment could have potentially modulated gene expression during a critical window for brain development in TS mice, these effects may have been reverted after treatment withdrawal. In WT mice, five genes were found to be differentially expressed between WT untreated and WT treated mice, of which *Hspa1b* and *Hspa1a* mapped to Mmu17, *Fosl2* mapped to Mmu5, *Actn1* to mapped Mmu12, and *Tm6sf2* mapped to Mmu8; which could indicate permanent genome-wide treatment effects in a few genes in WT mice. Interestingly, from the list of DEGs, only *Hspa1b* was found to interact with *Dyrk1A* (Rouillard et al., 2016), suggesting that GTE-EGCG may have other mechanisms of action than DYRK1A kinase activity inhibition.

### The pleiotropic nature of genotype and treatment effects

The combination of multi- and univariate tests performed in our study allowed us to explore in a more comprehensive way the integrated effects of genotype and treatment. Overall, we detected more significant differences when performing multivariate analysis combining the results of different tests together than when we evaluated each test individually, which highlights the pleiotropic nature of DS and GTE-EGCG, as both genotype and treatment caused mild effects in multiple readouts rather than large specific effects in single variables.

For example, we observed that even though TS untreated mice presented mild gene expression alterations in the cerebellum, these mice still showed hyperactivity and increased cerebellar volume. Similarly, TS treated mice presented a larger cerebellum and hyperactivity when compared to WT untreated mice after treatment discontinuation, but only 12 genes were found to be differentially expressed between WT untreated and TS treated mice at that stage. These results can be interpreted as that the altered expression of a few genes was sufficient to alter the brain and cognitive phenotypes, or that the alteration of gene expression during development was no longer detected at adulthood but was sufficient to permanently alter brain volumetry and cognition, which would be in line with previous reports indicating that the altered phenotype in DS was due to small contributions of multiple genes rather than strong effects of a few selected genes (Antonarakis et al., 2004; Chang et al., 2020; Chrast et al., 2000; De Toma et al., 2021). Interestingly, trisomy altered the expression of genes related with hypotonia and movement impairment such as *Son*, *Synj1*, *Atp5o*, *Jam2* and *Paxbp1*; but TS mice still showed increased locomotor activity.

Furthermore, we observed that even though there were no significant differences in the concentration of NAA in the hippocampal region of TS untreated mice, these mice showed altered hippocampal region metabolite spectra, increased hippocampal region volume, and altered memory robustness. Similarly, the treatment did not alter the concentration of NAA in the hippocampal region of TS treated mice, but induced general differences in hippocampal region metabolite spectra, mildly increased hippocampal region volume, and altered cognition. These results suggested that the cognitive differences induced by both the genotype and treatment were not related with the concentration of NAA in the hippocampal region but rather with general differences in hippocampal region volume and metabolite spectra.

### The advantages of a longitudinal holistic approach

Single holistic and longitudinal experiments evaluating multiple systems simultaneously overcome the limitations of individual analyses and the lack of consistency across studies. The differences between study results highlighted in this study could be due to a large variety of factors, including experimental differences in the mouse model, developmental stage analyzed, sex distribution of the sample, experimental setup, technical differences in data acquisition and analysis, as well as differences in treatment dose, timing, and route of administration. As a result, evidence from different studies is challenging to interpret, and can lead to misinterpretations about the course of the disorder and the effects of pharmacological treatments. With our approach, the differences between experimental setups could be accounted for, analyzing the simultaneous development of different systems in the same mice in a controlled experimental setup, and investigating the related effects of trisomy and treatment in multiple related systems.

Our results support that, overall, the Ts65Dn model reflects the multisystemic nature of DS and recapitulates many characteristics of the trisomic phenotype. As compared to WT mice, Ts65Dn mice presented a trend for reduced body weight over development together with a brachycephalic skull with facial flatness and a trend for reduced BMD in the humerus. These skeletal alterations co-occurred with cognitive delay during early stages of development, hyperactivity, impaired long-term memory, and increased explorative and risk-taking behavior at adulthood. At the molecular and genetic level, Ts65Dn mice presented alterations in the hippocampal metabolite spectra and differential gene expression in the cerebellum. However, our results also revealed phenotypes that did not match the human condition, as Ts65Dn mice did not show altered tibia microarchitecture and showed increased hippocampal, cerebellar, ventricular, and whole brain volumes. Overall, our longitudinal analyses confirm the validity of the model for the structural, functional, molecular and genetic phenotypes associated with DS, but also support the hypothesis of genotypic and phenotypic drift within the Ts65Dn mouse model (Shaw et al., 2020), as we generally observed a decrease in the magnitude of alterations induced by the genotype compared to previous articles including our earlier work.

Regarding the treatment effects, our results confirmed that GTE-EGCG modulated most of these systems simultaneously along development. However, our holistic approach revealed that, in general, the treatment did not rescue the trisomic phenotype and even exacerbated some phenotypes over time, such as body weight, tibia microarchitecture, neurodevelopment, adult cognition, and hippocampal metabolite concentration. Although discontinuing the GTE-EGCG administration reduced the treatment effects, it did not rescue the trisomic phenotype, as TS treated mice remained different from WT untreated mice in most systems. Discontinuing the GTE-EGCG treatment for three months only rescued the body weight and brain volume of a few TS mice but increased the weight of WT mice and reduced the BMD of the humerus in both WT and TS mice. Summing up, our preclinical results warn against a chronic GTE-EGCG treatment initiated prenatally and maintained until adulthood with a dosage of 30 mg/kg/day.

## FUTURE DIRECTIONS

Performing specific experiments to evaluate the effects of a potential therapeutic compound on one structure at one timepoint are important first steps to screen for novel therapeutic agents. However, the pleiotropic effects on all involved organ systems over a relevant time course should not be ignored when evaluating its general safety and efficacy, especially in complex disorders like DS.

The experimental pipeline used in this article, from the imaging techniques to the data analysis strategy, are applicable to any rodent model and potential therapeutic compound, allowing the integrated investigation of other therapeutic compounds and DS models that more faithfully replicate the genetic condition of DS, such as the TcHSA21rat and TcMAC21 mouse model (Kazuki et al., 2020; Kazuki et al., 2022). Furthermore, the simultaneous direct and indirect effects of potential therapeutic agents could be investigated in an integrated manner for other disorders and syndromes with multi-systemic alterations such as Apert, Pfeiffer and Crouzon craniosynostosis syndromes (Caputo et al., 2016; Fernandes et al., 2016; Monteagudo, 2020; Nopoulos et al., 2007; Pirozzi et al., 2018; Roberts et al., 2012; Treit et al., 2016; Vogels & Fryns, 2006; Wilhoit et al., 2017; Wozniak et al., 2019), considering the associated effects in different systems.

With this holistic approach, the preclinical biomedical research field will be able to solve new challenges and answer new questions, understanding the complexities of systemic diseases in a generalized manner and providing a global context to the contributions of specific genes, proteins, molecules, and compounds.

## MATERIALS AND METHODS

### Animals, housing, treatment, and experimental design

Ts65Dn (B6EiC3Sn-a/A-Ts (1716)65Dn) females and B6EiC3Sn.BLiAF1/J males (refs. 005252 and 003647, the Jackson Laboratory Bar Harbor, ME, USA) were obtained from the Jackson Laboratory and crossed within six months to obtain F1 trisomic Ts65Dn (TS) mice and euploid wildtype littermates (WT) that were used throughout the experiment. Mice were housed at the animal facility of KU Leuven in individually ventilated cages (IVC cages, 40 cm long x 25 cm wide x 20 cm high) under a 12h light/dark schedule in controlled environmental conditions of humidity (50% - 70%) and temperature (22 ± 2°C) with food and water supplied *ad libitum*. Date of conception (E0) was determined as the day in which a vaginal plug was present. After birth, all pups were labeled with a non-toxic tattoo ink (Ketchum Animal Tattoo Ink, Green Paste) for identification throughout the longitudinal experiments, as the same mice were used throughout the entire experiment. All procedures complied with all local, national, and European regulations and ARRIVE guidelines and were authorized by the Animal Ethics Committee of KU Leuven (ECD approval number P120/2019).

Mice were genotyped at PD1 by PCR from tail snips adapting the protocol in (Shaw et al., 2020). Trisomic primers, Chr17fwd-5′-GTGGCAAGAGACTCAAATTCAAC-3′ and Chr16rev-5′- TGGCTTATTATTATCAGGGCATTT-3′; and positive control primers, IMR8545-5′- AAAGTCGCTCTGAGTTGTTAT-3′ and IMG8546-5′- GAGCGGGAGAAATGGATATG-3′ were used. The following PCR cycle conditions were used: step 1: 94°C for 2 min; step 2: 94°C for 30 s; step 3: 55°C for 45 s; step 4: 72°C for 1 min (steps 2-4 repeated for 40 cycles); step 5: 72°C for 7 min, and a 4°C hold. PCR products were separated on a 1% agarose gel.

We bred a total of 13 litters. Six litters were left untreated and seven litters were treated via the drinking water with GTE-EGCG (Mega Green Tea Extract, Life Extension, USA) at a concentration of 0.09 mg EGCG/mL, as calculated based on the label concentration (45% EGCG per capsule). As EGCG crosses the placental barrier and reaches the embryo (K.O. Chu et al., 2006), GTE-EGCG treatment started prenatally at embryonic day 9 (E9) via the drinking water of the pregnant dams. After weaning at postnatal day (PD) 21, GTE-EGCG dissolved in water at the same concentration was provided to the young mice *ad libitum* until 5 months (5M), when the treatment was discontinued (Fig. 1). The treatment was prepared freshly every day and water intake was monitored in each cage. The calculated dosage of EGCG received by an adult mouse would be approximately 30 mg/kg/day considering that, on average, early adult mice weigh 20 g and drink 6 mL of water per day according to our measurements. In developing embryos and pups before weaning, the received dosage was lower since previous studies indicate that maternal plasma concentrations of catechins are about 10 times higher than in placenta and 50–100 times higher than in the fetal brain (K. O. Chu et al., 2006) and EGCG in milk and plasma of PD1 to PD7 pups was detected at low concentrations (Souchet et al., 2019).

Mice were allocated to groups according to their genotype and pharmacological intervention: WT and TS mice untreated or treated with GTE-EGCG (Fig. 1). Investigators were blinded to genotype during animal experimentation, and to genotype and treatment during data analysis. The same mice were longitudinally used throughout the experiment. Sample sizes varied across groups and developmental stages due to uncontrollable technical issues inherent to longitudinal studies, such as scanning failure or mouse death during the experiment. The litter information containing litter number, treatment administration and sex for each mouse is described in Supplementary Table S15. Detailed information regarding sample sizes for each experiment and analysis is provided in Supplementary Table S1.

### Structural assessment

#### Body weight

Mice body weight was recorded daily from PD1 to PD17 and before each µCT scanning (Fig. 1).

#### Skeletal development

##### In vivo µCT

We performed high resolution longitudinal *in vivo* µCT at four timepoints from after birth until adulthood to monitor skeletal development (Fig. 1). Mice were anesthetized by inhalation of 1.5-2% of isoflurane (Piramal Healthycare, Morpeth, Northumberland, United Kingdom) in pure oxygen and scanned *in vivo* with the SkyScan 1278 (Bruker Micro-CT, Kontich, Belgium) for three minutes using the optimized parameters specified in Supplementary Table S16. *In vivo* µCT data was reconstructed using a beam hardening correction of 10% (NRecon software, Bruker Micro-CT, Kontich, Belgium).

###### • Skull shape analysis

Skull 3D models were automatically generated from reconstructed *in vivo* µCT scans by creating an isosurface based on specific threshold for bone using Amira 2019.2 (Thermo Fisher Scientific, Waltham, MA, USA). We compared craniofacial morphology in WT and TS mice with and without GTE-EGCG treatment using Geometric Morphometric quantitative shape analyses (Dryden & Mardia, 1998; Hallgrimsson et al., 2015; James Rohlf & Marcus, 1993; Klingenberg, 2010). The analysis was based on the 3D coordinates of anatomical homologous landmarks recorded over the skull and face at each developmental stage as described before (Llambrich, González-Colom, et al., 2022). The landmark configuration for each stage is defined in Supplementary Figure S1 and Supplementary Table S17. Landmarks were acquired using Amira 2019.2.

###### • Humerus bone mineral density (BMD)

To calculate humerus BMD from the µCT data, we first computed the humerus mean grey value of each mouse by delimiting a volume of interest of ten slices that was placed right below the deltoid protuberance of the humerus using the CTAn software (Bruker Micro-CT, Kontich, Belgium). Then, we scanned two phantoms with different known densities of hydroxyapatite (100 mg/cm3 and 500 mg/cm3) using the same settings as in the *in vivo* scans (Supplementary Table S16). A calibration line was obtained between the known hydroxyapatite densities and their corresponding grey values. The resulting equation was applied to calculate the BMD of the humerus of each mouse from their mean grey value.

##### Tibia length

After sacrifice, the length of the right tibia was measured in all mice using a digital caliper.

##### Ex vivo µCT for tibia

The proximal region of the tibia was scanned *ex vivo* using the Skyscan 1272 high-resolution µCT scanner (Bruker Micro-CT, Kontich, Belgium) with the optimized parameters specified in Supplementary Table S16. After scanning the bones, the raw 2D images were reconstructed using NRecon (version 1.7.3.1, Bruker Micro-CT, Kontich, Belgium) and rotated to a standard position using DataViewer (version 1.5.6.2, Bruker Micro-CT, Kontich, Belgium). The reconstructed images were then analyzed using the CTAn software (version 1.17.8.0, Bruker Micro-CT, Kontich, Belgium) as follows.

###### • Trabecular analysis

For trabecular bone, a section of 300 slices (1.5 mm) was selected starting 100 slices (0.5 mm) underneath the point in the proximal tibia where the articular condyles met. Then, a region of interest (ROI) was manually defined including the trabecular bone inside the thin cortical outer layer. The descriptions, abbreviations and parameter units are provided in Supplementary Table S18.

###### • Cortical analysis

For cortical bone, a section of 100 slices (0.5 mm) was selected in CTAn starting 600 slices (3 mm) underneath the reference point in the proximal tibia. The tissue inside the medullary canal was excluded from the ROI. The descriptions, abbreviations and parameter units are provided in Supplementary Table S19.

#### Brain volume

##### In vivo MRI scanning

Mice were also MR scanned *in vivo* under the same anesthesia (1.5-2% isoflurane, Piramal Healthycare, Morpeth, Northumberland, United Kingdom) at MRI_1_ and MRI_2_ (Fig. 1) with a 9.4 T Bruker Biospec 94/20 small animal μMR scanner (Bruker Biospin, Ettlingen, Germany; 20 cm horizontal bore) equipped with actively shielded gradients (maximum gradient strength 600 mT m^−1^). Axial, coronal, and sagittal images were acquired using a 2D T2 weighted Rapid Acquisition with Relaxation Enhancement (RARE) sequence (repetition time (TR)/ echo time (TE): 3781 / 33 ms; RARE factor: 8; averages: 6; field of view (FOV): 20 × 20 mm; matrix 128 × 128; slice number: 35; slice thickness: 0.4 mm; slice gap: 0.1 mm; acquisition time 6 min). A quadrature radiofrequency resonator (inner diameter 7.2 cm, Bruker Biospin) was used for transmission of radiofrequency pulses in combination with and actively decoupled mouse brain surface coil for reception (Bruker Biospin).

##### Segmentation of brain regions of interest

After MR image acquisition, brain masks were manually delineated on the axial plane for each mouse using 3D Slicer v5.0.2. (http://www.slicer.org) (Fedorov et al., 2012). Then, the masks were fed to the Atlas-Based Imaging Data Analysis (AIDA) pipeline described previously (Pallast et al., 2019). In brief, the pipeline consisted of a series of preprocessing steps including skull stripping and bias field correction of the MR images before registration with the Allen Mouse Brain Reference Atlas (Sunkin et al., 2013) through a series of affine and non-linear transformations. The volume of the whole brain was extracted from the manually delineated masks and the volumes of the hippocampal region, cerebellum, and ventricles were extracted from the AIDA segmentations. All brain volumes were normalized to body weight to account for the differences in overall body size between WT and TS mice.

### Functional assessment

#### Neurobehavioral development

Neurobehavioral developmental tests were carried out daily from PD1 to PD18, as previously described (Dierssen et al., 2002; Llambrich, González, et al., 2022). Mothers were separated from their pups before testing. Pups were then taken out one at a time from their home cage for testing, and mothers were returned into the cage after all pups were evaluated. For each neurobehavioral test, we evaluated the acquisition rate and the average day of successful test completion.

In neurodevelopment tests with a presence/absence binary outcome, such as eye opening, pinna detachment, walking, cliff drop aversion, Preyer reflex, blast response, visual placing, reaching response, vibrissae placing and tactile response; the acquisition rate was scored as the percentage of mice that successfully acquired the landmark or response behavior on each day. The day of successful test completion was considered as the day when there was a positive response.

In those neurodevelopment tests measured with categorical non-binary scores, such as incisor eruption, surface righting response, negative geotaxis, vertical climbing, and grasping; the acquisition rate was scored as the daily average score of each group of mice. The day of successful test completion was considered as the day when the highest score was achieved.

#### Maternal care

For maternal care monitoring, home IVC cages were transferred to a separate light cycle-controlled room with food and water supplied *ad libitum*. Cages were videotaped from the top during 24h using the Foscam C1 camera with night vision and a transparent Plexiglas cover with holes for ventilation. The recordings were manually inspected every 6 minutes for 10 seconds, and maternal behavior was categorized as nursing, pup grooming, digging in nest, eating, drinking, moving, or digging off nest to evaluate the frequency of each maternal behavior.

#### Adult cognition

Open Field (OF), Elevated Plus Maze (EPM), Sociability/Preference for Social Novelty (SPSN), Novel Object Recognition (NOR) and Passive Avoidance (PA) tests were performed in this order before and after treatment discontinuation (Fig. 1).

The OF test was performed in a brightly illuminated Plexiglas arena (50 x 50cm) with transparent walls. Dark habituated (30 min) mice were placed in the left bottom corner facing the walls and were left free to explore the arena for 10 minutes. Movements were recorded using a camera and the tracking software ANY-mazeTM Video Tracking System software (Stoelting Co., IL, USA). Exploration towards the center of the field was considered a readout for reduced anxiety.

The EPM test took place on a plus shaped maze with 2 open and 2 closed arms (5 cmx30 cm) that was elevated 35 cm from the tabletop. Each mouse was placed in the left closed arm, with the snout pointing away from the crossing. After a 1-minute habituation time, the trial was initiated manually, letting the mouse spend 10 minutes in the arena. Four infrared (IR) beams connected to an activity logger recorded the arm entries of the mouse and one beam recorded the percentage of time that the mouse spent in the open arms per minute.

The SPSN set-up consisted of a Plexiglas box (60 x 15cm) with three compartments separated with perforated Plexiglas walls. The SPSN test involved three trials. At the first stage of habituation, the test mouse was left to explore the middle chamber for 300 seconds, while the left and right chambers were empty and visible from the middle chamber. Next, in the Social Preference stage with Subject 1 (S1), the test mouse was placed in the middle chamber for 300 seconds while one stranger mouse (STR1) was placed in either the left or right chamber, and the other chamber was left empty. Social approach was recorded as time spent close to STR1, and a preference ratio was calculated (Pref=100* Time close to STR1/ (Time close to STR1 + Time close to Empty)). Finally, in the Social Novelty stage with Subject 2 (S2) (300 s), a second stranger mouse (STR2) was placed in the previously empty chamber. Social recognition memory was scored as preference towards STR2 (calculated as ratio: pref =100* (time close to STR2) / (Time close to STR1 + Time close to STR2)). The two stranger mice were C57BL/6 wildtype mice of the same sex as the test mouse and had served before as stranger mice in previous SPSN experiments. Explorative social behavior towards stranger mice was measured using ANY-mazeTM Video Tracking System software (Stoelting Co., IL, USA).

NOR testing started with a habituation phase where the animals were placed during 15 minutes in a dimly lit open field arena (wooden box 40X40cm painted white). Twenty-four hours later, the animals were reintroduced to the same arena for 10 minutes with two identical falcon tubes filled with colored liquid that were placed in opposite corners equally distant from the mouse. Exploration time involving sniffing in close proximity (<2cm) was recorded by an overhead camera and tracking software. Total exploration time was set at minimally 15 seconds to ensure that object characteristics were encoded. Sixty minutes later, the animals were placed in the arena for 10 minutes, with one object replaced by a novel object, and exploration time was recorded. Preference for novel object was calculated as ratio (Pref=100* Time novel object / (Time familiar object + novel object)). Objects were randomly assigned as familiar or novel for each mouse, and the left or right position of the novel object was counterbalanced between trials. Exploration time was measured by a camera and ANY-mazeTM Video Tracking System software (Stoelting Co., IL, USA).

The PA experimental set-up consisted of a transparent box illuminated with an LED lamp leading to a dark box with an electrifiable grid connected to a shocker (LE 100-26, Panlab Bioseb, Spain) and a lid. Dark habituated mice were placed in the light box and when they entered the dark box (CS), the latency to enter the dark box was recorded, and a mild foot shock was delivered (US 0,5mA, 2s, scrambled). The next day, the trial was repeated (without foot shock presentation) and latency to enter the dark compartment was recorded (maximum 300s). Animals with good memory retention would display a higher latency to enter on day 2. This test was repeated in the same animals after treatment cessation. We noticed that upon re-exposure to the same setup, some animals still remembered the CS-US presentation, and refused to enter the dark box. Therefore, this second testing session reflected the stability of long-term memory.

### Molecular and genetic assessment

#### In vivo MRS

MR spectra were acquired as previously reported using a Bruker Biospec 94/20 MR scanner (Vanherp et al., 2021; Weerasekera et al., 2018). After MRI scanning, MR spectra were acquired from a 2.5 × 1.25 × 1.5 mm voxel placed in the hippocampal region of the brain, using a PRESS sequence with TR/TE 2000/20 ms, 320 averages, and localized shimming with no margin. Water suppression was optimized using VAPOR (Griffey & P. Flamig, 1990). An unsuppressed water MR spectrum was acquired before each water-suppressed 1H-MRS spectrum for quantification/referencing. Shimming was performed using FASTMAP, resulting in a final water line width at half height < 20 Hz.

For the multivariate analysis of the spectra, the signals were truncated to retain only the region of interest between 0.0 and 4.3ppm. The msbackadj function in Matlab (Inc., 2022) was applied for baseline and offset correction. The baseline was estimated within multiple shifted windows of width 150 separation units and extracted from the original signal. The baseline corrected signals were further analyzed by segmenting it into 12 spectral regions, which were integrated and normalized to the total integral using peak integration methodology for metabolite quantification (Supplementary Figure S2). The two spectral regions that represent contaminations from macromolecules were excluded (number 11 and 12 in Supplementary Figure S2). Main metabolites present in the respective spectral regions are listed in Supplementary Table S10.

For quantification of absolute metabolite concentrations, a similar approach was taken as previously reported (Weerasekera et al., 2018). In brief, spectra were processed using jMRUI v6.0 (Stefan et al., 2009). Spectra were phase corrected and an HLSVD (Hankel Lanczos Singular Values Decomposition) filter was applied to remove the residual water signal (van den Boogaart, 1994). Metabolites were quantified with the QUEST algorithm (Ratiney et al., 2004) in jMRUI using a simulated (NMRScopeB) basis set (Starčuk Jr et al., 2009). Results were reported in reference to the non-suppressed water signal. A metabolite data base was used as in (Weerasekera et al., 2018).

#### Gene expression

Gene expression analysis was performed on a cerebellar tissue homogenate at endpoint (Fig. 1). Each cerebellum was dissected and processed independently. Homogenates were obtained with a gentleMACS dissociator (Miltenyi Biotech). Total RNA was then extracted with QIAzol according to the manufacturer’s instructions. RNA purity and concentration were assessed by NanoDrop ND-1000 Spectrophotometer and RNA integrity was evaluated by Fragment Analyzer analysis (RIN≥8). Illumina TruSeq stranded mRNA kit was used for library preparation, samples were pooled and sequenced on a HiSeq4000, single end, 50bp reads. A minimum of one million reads were obtained per sample. Quality control of raw reads was performed with FastQC v0.11.7 (Andrews, 2010). Adapters were filtered with ea-utils fastq-mcf v1.05 (Aronesty, 2011). Splice-aware alignment was performed with HISat2 (Kim et al., 2019), against the mouse reference genome mm10 using default parameters. Reads mapping to multiple loci in the reference genome were discarded. Resulting Binary Alignment Map (BAM) files were handled with Samtools v1.5 (Li et al., 2009). Quantification of reads per gene was performed with HT-seq Count v0.10.0, Python v2.7.14 (Anders et al., 2014). Count-based differential expression analysis was performed with R-based (The R Foundation for Statistical Computing, Vienna, Austria) Bioconductor package DESeq2 (Love et al., 2014), normalizing absolute counts. Pairwise comparison of the entire genome for all groups was done with default settings and the reported P-values were adjusted for multiple testing with the Benjamini-Hochberg procedure controlling for false discovery rate (FDR) (Supplementary Table S14). Multivariate evaluation of the subset of 125 triplicated genes present in the Ts65Dn mouse model was performed as described below. All mice that survived until endpoint were included in both analyses (Supplementary Table S1).

### Statistics

#### Univariate evaluation

The developmental trajectories of the body weight, BMD, and acquisition rate of neurodevelopment tests with categorical non-binary scores were longitudinally analyzed by fitting a mixed-effects model as implemented in GraphPad Prism 8.0, using the Geisser-Greenhouse correction and Restricted Maximum Likelihood (REML) fit as described before (Llambrich, González, et al., 2022). The acquisition rate of neurodevelopment tests with a presence/absence binary outcome was longitudinally analyzed using a log-rank test (Mantel-Haenszel approach), considering the day of appearance of the landmark or response as an event using GraphPad Prism 8.0.

We made five pairwise comparisons for all univariate non-longitudinal data: the body weight at adulthood, the BMD at each developmental timepoint, the variables evaluating tibia microarchitecture, the brain volume of the different regions, the average day of acquisition of the neurodevelopmental tests, the variables evaluating adult cognition, and the concentration of the hippocampal metabolites. We compared WT vs. TS untreated mice to evaluate the genotype effect (1), WT untreated vs. WT treated mice to evaluate the treatment effect in the WT background (2), TS untreated vs. TS treated mice to evaluate the treatment effect in the trisomic background (3), WT untreated vs. TS treated mice to determine whether the treatment had a rescuing effect in trisomic mice (4), and WT treated vs. TS treated mice to evaluate whether the treatment showed different effects in the WT and trisomic background (5). We determined statistical significance for each comparison using univariate statistical tests as previously described (Llambrich, González-Colom, et al., 2022). For the BMD, P-values at each developmental stage were adjusted for multiple comparisons using the Benjamini-Hochberg (Q = 5%) test.

The results for normality, homoscedasticity and statistical tests performed for each variable can be found in Supplementary Table S20. Mice identified as outliers by the ROUT test (Motulsky & Brown, 2006) with a Q (maximum desired False Discovery Rate) of 1% were excluded from the analysis. All univariate statistical analysis were performed using GraphPad Prism (v8.02, GraphPad Software, San Diego, California USA).

#### Multivariate evaluation

We performed multivariate statistics in all tests with multiple variables: craniofacial shape, tibia microarchitecture, brain volumes, neurodevelopmental tests, adult cognitive tests, brain metabolite concentration and gene expression.

We performed a principal component analysis (PCA) for the craniofacial shape and gene expression analysis. The PCA for the craniofacial shape analysis was based on the 3D coordinates of the set of 27 landmarks recorded to capture craniofacial shape. To extract shape information from the 3D landmark configurations, we performed a Generalized Procrustes Analysis (GPA) followed by a PCA at each stage as described before (Llambrich, González, et al., 2022) using MorphoJ v1.06d (Klingenberg, 2011). For the gene expression data, the PCA was based on the rlog normalized expression data of the 125 triplicated genes in the Ts65Dn mouse model according to the MGI-Mouse Genome Informatics Database that were present in our dataset and was performed using PAST v4.1 (Hammer et al., 2001).

As we aimed to maximize differences between groups, we performed a linear discriminant analysis (LDA) for the tibia microarchitecture analysis, brain volumetric analysis, neurodevelopmental tests, adult cognitive tests, and hippocampal metabolite concentration analysis using the results obtained from each test. The variables included in each LDA are shown in Supplementary Table S21. We performed an LDA for each domain considering genotype+treatment as the grouping variable using PAST v4.1. As we compared four groups of mice, the LDA created three new axes that were independent among them and explained 100% of variation across groups. In the LDA, the separation among groups was determined by Mahalanobis distances, which account for correlations between standardized variables, and allow to combine measurements with different units in the same analysis.

If slight or no differences were associated with DS or treatment, the mice groups overlapped in the PCA or LDA scatterplot, showing similar phenotypes. If there were differences, the different groups of mice separated from each other.

To statistically quantify differences between WT, TS, WT treated and TS treated mice and answer the five scientific questions formulated above, we performed pairwise permutation tests with 10.000 rounds following the PCAs and LDAs. For the craniofacial shape analysis, we obtained the P-values from the permutation tests based on the Procrustes distances between the average shape of pairs of groups at each developmental stage using MorphoJ v1.06d (Klingenberg, 2011). For the gene expression analysis, we obtained the P-values from the pairwise comparisons after a one-way PERMANOVA based on Euclidean distances using PAST v4.1 (Hammer et al., 2001). For the tibia microarchitecture tests, brain volumetric tests, neurodevelopmental tests, adult cognitive tests, and hippocampal metabolite concentration tests we obtained the P-values from the pairwise comparisons after a one-way PERMANOVA based on Mahalanobis distances using PAST v4.1 (Hammer et al., 2001).

Finally, to better understand the differences between groups, we selected the variables that were testing for a certain domain and grouped them into categories for the tibia microarchitecture tests, neurodevelopmental tests and adult cognitive tests. We then repeated the multivariate pairwise permutation analysis. The categories were cortical bone strength, cortical bone size and trabecular bone for the tibia microarchitecture; developmental landmarks, neuromotor tests and reflexes for the neurodevelopmental tests; and anxiety, arousal, and memory for the adult cognitive tests.

## SUPPLEMENTARY MATERIAL

**Supplementary Figure S1.**
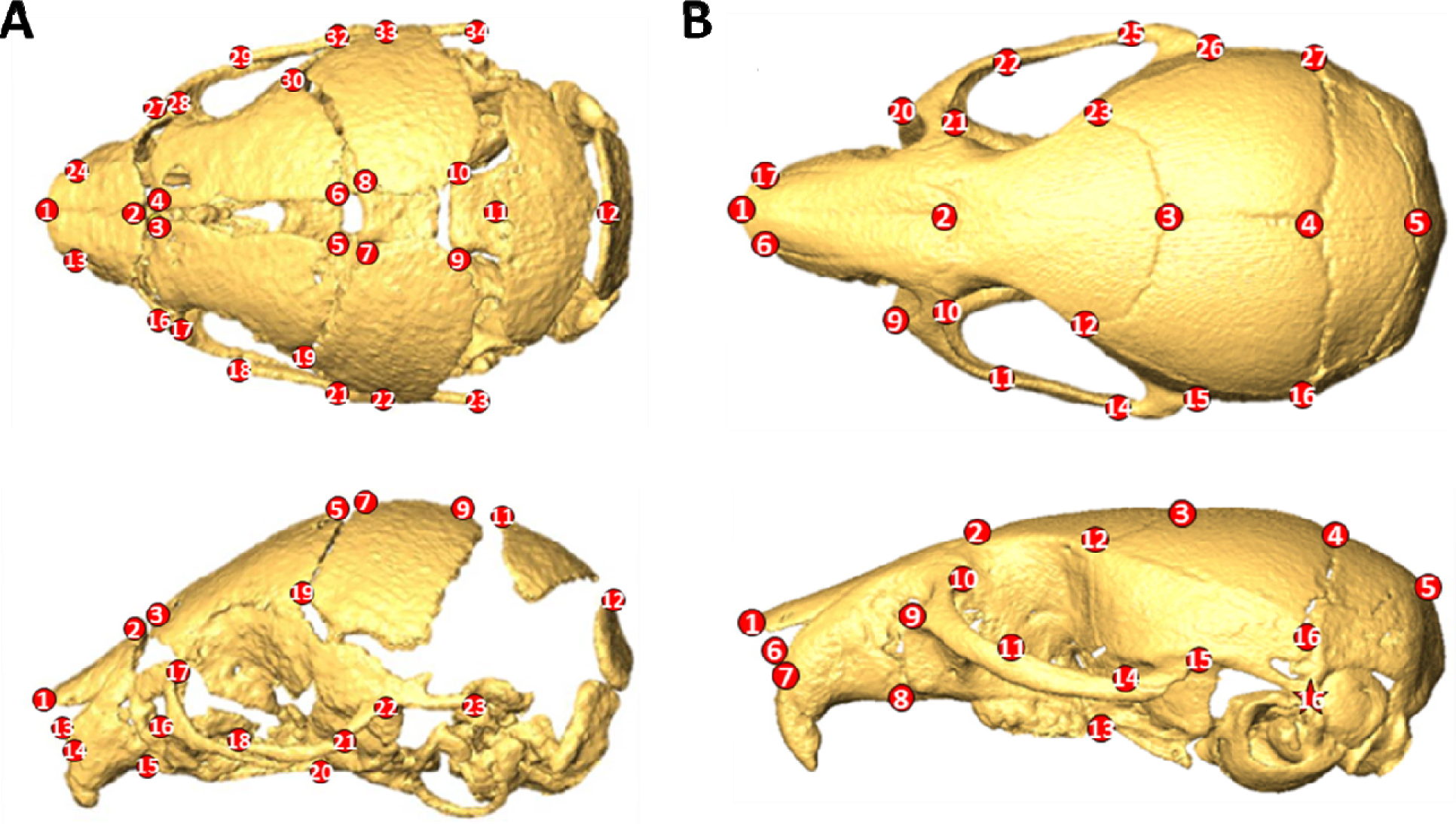
Set of anatomical landmarks used to characterize the shape of the skull and face. **(A)** Set of 34 landmarks characterizing craniofacial shape at µCT_1_ from a 3D reconstruction of a µCT scan. **(B)** Set of 27 landmarks characterizing craniofacial shape at µCT_2_, µCT_3_ and µCT_4_ from a 3 reconstruction of a µCT scan. See Supplementary Table S17 for precise anatomical definitions.

**Supplementary Figure S2.**
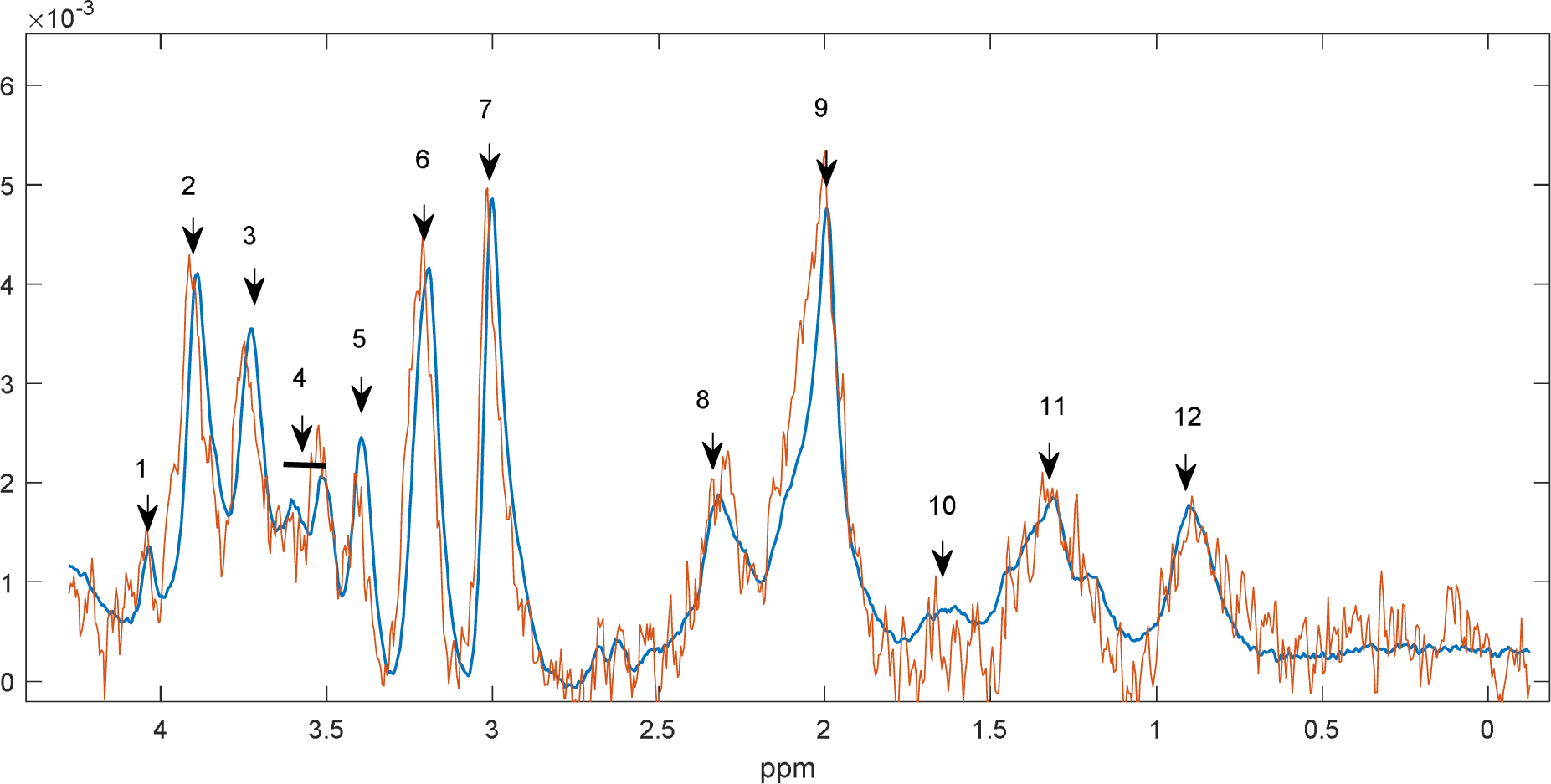
MR spectra and integrated spectral regions for multivariate MRS analysis. The region from 0 to 4.3 ppm of a representative MR spectrum is shown. The integrated regions and the most concentrated metabolites contributing to those regions are indicated in Supplementary Table S10. Regions 11 and 12, representing contaminations from macromolecules, were not included in the analysis as they showed large variability.

**Supplementary Table S1.** Sample size for each analysis, experiment, and developmental stage. Differences in sample size between stages are due to technical reasons such as micro-CT or MR scanner not operating on the scanning day, movement scanning artifacts, or mouse death during the experiment. For the tibia microarchitecture, differences in sample size are due to mice detected as outliers. For the MRS, differences in sample size are due to signal artifacts. For the adult cognitive tests, differences in sample size are due to artifacts during tracking.

|  | WT |  | TS |  | WT Treated |  | TS Treated |  |
| --- | --- | --- | --- | --- | --- | --- | --- | --- |
|  | Female | Male | Female | Male | Female | Male | Female | Male |
| BW1 | 11 | 7 | 11 | 7 | 13 | 10 | 14 | 8 |
| BW2 | 7 | 3 | 5 | 3 | 7 | 10 | 11 | 7 |
| BW3 | 5 | 2 | 5 | 2 | 7 | 10 | 9 | 7 |
| μCT1 | 11 | 7 | 10 | 7 | 9 | 13 | 8 | 8 |
| μCT2 | 8 | 5 | 6 | 4 | 10 | 9 | 9 | 7 |
| μCT3 | 8 | 3 | 6 | 2 | 8 | 10 | 11 | 7 |
| μCT4 | 6 | 3 | 6 | 2 | 7 | 10 | 11 | 7 |
| BMD humerus μCT1 | 11 | 7 | 10 | 7 | 9 | 13 | 14 | 8 |
| BMD humerus μCT2 | 4 | 5 | 6 | 4 | 9 | 9 | 11 | 7 |
| BMD humerus μCT3 | 8 | 3 | 6 | 2 | 7 | 10 | 11 | 7 |
| BMD humerus μCT4 | 5 | 4 | 6 | 2 | 8 | 8 | 11 | 7 |
| Tibia length | 6 | 3 | 6 | 2 | 7 | 10 | 11 | 7 |
| Percentage Bone Volume | 6 | 2 | 6 | 2 | 7 | 10 | 11 | 7 |
| Trabecular Thickness | 6 | 3 | 6 | 2 | 7 | 10 | 11 | 7 |
| Trabecular Separation | 6 | 3 | 6 | 2 | 7 | 10 | 11 | 7 |
| Trabecular Number | 6 | 3 | 6 | 2 | 7 | 10 | 11 | 7 |
| Cross Sectional Area | 6 | 3 | 6 | 2 | 7 | 10 | 11 | 7 |
| Periosteal Perimeter | 6 | 3 | 6 | 2 | 7 | 10 | 11 | 7 |
| Endocortical Perimeter | 6 | 3 | 6 | 2 | 7 | 10 | 11 | 7 |
| Bone Area | 6 | 3 | 6 | 2 | 7 | 10 | 11 | 7 |
| Medullary Area | 6 | 3 | 6 | 2 | 7 | 10 | 11 | 7 |
| Cortical Thickness | 6 | 3 | 6 | 2 | 7 | 10 | 11 | 7 |
| Cortical Porosity | 6 | 3 | 6 | 2 | 7 | 9 | 10 | 7 |
| Polar Moment Of Inertia | 6 | 3 | 6 | 2 | 7 | 10 | 11 | 7 |
| BMD | 6 | 3 | 6 | 2 | 7 | 10 | 11 | 7 |
| Cortical bone mineral density | 6 | 3 | 6 | 2 | 7 | 9 | 11 | 7 |
| Whole Brain (MRI1) | 7 | 4 | 6 | 3 | 7 | 10 | 11 | 7 |
| Hippocampus (MRI1) | 7 | 4 | 6 | 3 | 7 | 10 | 11 | 6 |
| Cerebellum (MRI1) | 7 | 4 | 6 | 3 | 7 | 10 | 11 | 7 |
| Lateral Ventricles (MRI1) | 7 | 4 | 6 | 3 | 7 | 10 | 11 | 7 |
| Third Ventricle (MRI1) | 7 | 4 | 6 | 3 | 7 | 10 | 11 | 7 |
| Fourth Ventricle (MRI1) | 7 | 4 | 6 | 3 | 7 | 10 | 11 | 7 |
| Cerebral Aqueduct (MRI1) | 7 | 4 | 6 | 3 | 7 | 10 | 11 | 7 |
| Whole Ventricles (MRI1) | 7 | 4 | 6 | 3 | 7 | 10 | 11 | 7 |
| Whole Brain (MRI2) | 6 | 3 | 6 | 2 | 7 | 10 | 11 | 7 |
| Hippocampus (MRI2) | 6 | 3 | 6 | 2 | 7 | 10 | 11 | 7 |
| Cerebellum (MRI2) | 6 | 3 | 6 | 2 | 7 | 10 | 11 | 7 |
| Lateral Ventricles (MRI2) | 6 | 3 | 6 | 2 | 6 | 10 | 11 | 7 |
| Third Ventricle (MRI2) | 6 | 3 | 6 | 2 | 7 | 10 | 11 | 7 |
| Fourth Ventricle (MRI2) | 6 | 3 | 6 | 2 | 6 | 10 | 11 | 7 |
| Cerebral Aqueduct (MRI2) | 6 | 3 | 6 | 2 | 7 | 10 | 11 | 7 |
| Whole Ventricles (MRI2) | 6 | 3 | 6 | 2 | 7 | 10 | 11 | 7 |
| Eye Opening | 9 | 5 | 6 | 7 | 10 | 10 | 12 | 8 |
| Vibrissae | 9 | 5 | 6 | 7 | 10 | 10 | 12 | 8 |
| Blast | 9 | 5 | 6 | 7 | 10 | 10 | 12 | 8 |
| Cliff | 9 | 5 | 6 | 7 | 10 | 10 | 12 | 8 |
| Incisor | 5 | 3 | 2 | 3 | 10 | 7 | 7 | 2 |
| Righting | 9 | 6 | 7 | 8 | 10 | 11 | 11 | 8 |
| Negative Geotaxis | 9 | 5 | 6 | 6 | 10 | 10 | 11 | 8 |
| Climbing | 8 | 5 | 6 | 6 | 9 | 10 | 11 | 4 |
| Grasping | 6 | 4 | 8 | 9 | 10 | 9 | 12 | 8 |
| Tactile | 9 | 5 | 6 | 6 | 10 | 10 | 12 | 8 |
| Reaching | 9 | 5 | 6 | 7 | 10 | 10 | 11 | 7 |
| Visual | 9 | 5 | 6 | 7 | 10 | 10 | 12 | 8 |
| Preyer | 9 | 5 | 6 | 7 | 10 | 10 | 12 | 8 |
| Pinna | 11 | 7 | 9 | 9 | 10 | 13 | 14 | 8 |
| Walking | 5 | 4 | 4 | 5 | 6 | 6 | 8 | 7 |
| Homing | 6 | 5 | 6 | 6 | 10 | 10 | 12 | 8 |
| Pivoting | 9 | 5 | 6 | 7 | 10 | 10 | 12 | 8 |
| Nursing | N/A | N/A | 6 | 0 | N/A | N/A | 7 | 0 |
| Pup licking | N/A | N/A | 6 | 0 | N/A | N/A | 7 | 0 |
| Digging in nest | N/A | N/A | 6 | 0 | N/A | N/A | 7 | 0 |
| Eating | N/A | N/A | 6 | 0 | N/A | N/A | 7 | 0 |
| Drinking | N/A | N/A | 6 | 0 | N/A | N/A | 7 | 0 |
| Moving | N/A | N/A | 6 | 0 | N/A | N/A | 7 | 0 |
| Digging off nest | N/A | N/A | 6 | 0 | N/A | N/A | 7 | 0 |
| OF: Time in center (Cog1) | 8 | 4 | 7 | 3 | 9 | 10 | 11 | 7 |
| OF: Time in periphery (Cog1) | 8 | 4 | 7 | 3 | 9 | 10 | 11 | 7 |
| NOR: Time in center hab (Cog1) | 8 | 4 | 7 | 3 | 9 | 10 | 11 | 7 |
| EPM: % open arm (Cog1) | 7 | 4 | 7 | 3 | 8 | 10 | 9 | 7 |
| OF: Distance (Cog1) | 8 | 3 | 7 | 3 | 9 | 10 | 11 | 7 |
| OF: Speed (Cog1) | 7 | 4 | 7 | 3 | 9 | 10 | 11 | 7 |
| SPSN: Distance hab (Cog1) | 8 | 4 | 7 | 3 | 9 | 10 | 11 | 7 |
| SPSN: Speed hab (Cog1) | 8 | 4 | 7 | 3 | 9 | 10 | 11 | 7 |
| NOR: Distance hab (Cog1) | 8 | 4 | 7 | 3 | 9 | 10 | 11 | 7 |
| NOR: Speed hab (Cog1) | 8 | 4 | 6 | 3 | 9 | 10 | 11 | 7 |
| NOR: Preference for NOV (Cog1) | 7 | 3 | 7 | 3 | 7 | 10 | 8 | 7 |
| SPSN: Preference for S2 (Cog1) | 7 | 3 | 7 | 3 | 7 | 10 | 8 | 7 |
| PA: Latency train (Cog1) | 8 | 4 | 5 | 3 | 8 | 10 | 10 | 6 |
| PA: Latency test (Cog1) | 8 | 4 | 7 | 3 | 8 | 10 | 11 | 7 |
| OF: Time in center (Cog2) | 6 | 2 | 5 | 2 | 7 | 10 | 11 | 7 |
| OF: Time in periphery (Cog2) | 6 | 2 | 6 | 2 | 7 | 10 | 11 | 7 |
| NOR: Time in center hab (Cog2) | 6 | 2 | 6 | 2 | 7 | 10 | 11 | 7 |
| EPM: % open arm (Cog2) | 6 | 2 | 6 | 2 | 7 | 10 | 11 | 7 |
| OF: Distance (Cog2) | 6 | 2 | 6 | 2 | 7 | 9 | 11 | 7 |
| OF: Speed (Cog2) | 6 | 2 | 6 | 2 | 7 | 9 | 11 | 7 |
| SPSN: Distance hab (Cog2) | 6 | 2 | 6 | 2 | 7 | 10 | 11 | 7 |
| SPSN: Speed hab (Cog2) | 6 | 2 | 6 | 2 | 7 | 10 | 11 | 7 |
| NOR: Distance hab (Cog2) | 6 | 2 | 6 | 2 | 7 | 9 | 11 | 7 |
| NOR: Speed hab (Cog2) | 6 | 2 | 6 | 2 | 7 | 9 | 11 | 7 |
| NOR: Preference for NOV (Cog2) | 4 | 2 | 4 | 2 | 7 | 10 | 11 | 7 |
| SPSN: Preference for S2 (Cog2) | 4 | 2 | 4 | 2 | 7 | 10 | 11 | 7 |
| PA: Latency train (Cog2) | 6 | 2 | 6 | 2 | 7 | 10 | 10 | 5 |
| PA: Latency test (Cog2) | 6 | 2 | 6 | 2 | 5 | 7 | 11 | 7 |
| NAA (MRS1) | 8 | 4 | 6 | 3 | 7 | 10 | 9 | 6 |
| Cr/PCr (MRS1) | 8 | 4 | 6 | 3 | 7 | 10 | 9 | 6 |
| Chol/Pchol/GPC (MRS1) | 8 | 4 | 6 | 3 | 7 | 10 | 9 | 6 |
| myo-Inos (MRS1) | 7 | 4 | 5 | 3 | 5 | 9 | 9 | 6 |
| Tau (MRS1) | 7 | 4 | 6 | 3 | 7 | 10 | 9 | 6 |
| NAA (MRS2) | 8 | 4 | 6 | 3 | 7 | 10 | 10 | 7 |
| Cr/PCr (MRS2) | 8 | 4 | 6 | 3 | 7 | 10 | 10 | 7 |
| Chol/Pchol/GPC (MRS2) | 8 | 4 | 6 | 3 | 7 | 10 | 10 | 7 |
| myo-Inos (MRS2) | 8 | 4 | 6 | 3 | 7 | 10 | 10 | 7 |
| Tau (MRS2) | 8 | 4 | 6 | 3 | 7 | 10 | 10 | 7 |
| RNAseq | 6 | 3 | 6 | 2 | 7 | 10 | 11 | 7 |

**Supplementary Table S2.** *P*-values resulting from permutation tests (10,000 permutation rounds) based on Procrustes distances among groups for craniofacial shape. Bold font indicates statistically significant values. Pairwise comparisons marked as N/A were not calculated since they did not evaluate any relevant scientific question.

|  |  |  |  |
| --- | --- | --- | --- |
| **μCT<sub>1</sub>** | TS Treated | TS Untreated | WT Treated |
| TS Untreated | **0.0005** |  |  |
| WT Treated | **<0.0001** | N/A |  |
| WT Untreated | **<0.0001** | **<0.0001** | **0.0418** |
| **μCT<sub>2</sub>** | TS Treated | TS Untreated | WT Treated |
| TS Untreated | 0.3358 |  |  |
| WT Treated | **<0.0001** | N/A |  |
| WT Untreated | **<0.0001** | **0.0011** | 0.2719 |
| **μCT<sub>3</sub>** | TS Treated | TS Untreated | WT Treated |
| TS Untreated | 0.1237 |  |  |
| WT Treated | **<0.0001** | N/A |  |
| WT Untreated | **<0.0001** | **0.0009** | 0.6242 |
| **μCT<sub>4</sub>** | TS Treated | TS Untreated | WT Treated |
| TS Untreated | 0.2151 |  |  |
| WT Treated | **<0.0001** | N/A |  |
| WT Untreated | **<0.0001** | **0.0055** | 0.5590 |

**Supplementary Table S3.** *P*-values resulting from the mixed-effects analysis and pairwise tests for humerus BMD. Bold font indicates statistically significant values after Benjamini–Hochberg correction. Pairwise comparisons marked as N/A were not calculated since they did not evaluate any relevant scientific question.

| **Mixed-effects analysis** | TS Treated | TS Untreated | WT Treated |
| --- | --- | --- | --- |
| TS Untreated | 0.0023 |  |  |
| WT Treated | 0.0320 | N/A |  |
| WT Untreated | <0.0001 | 0.2397 | <0.0001 |
| **μCT<sub>1</sub>** | TS Treated | TS Untreated | WT Treated |
| TS Untreated | 0.0897 |  |  |
| WT Treated | **0.0002** | N/A |  |
| WT Untreated | 0.0225 | 0.6717 | 0.3034 |
| **μCT<sub>2</sub>** | TS Treated | TS Untreated | WT Treated |
| TS Untreated | 0.5168 |  |  |
| WT Treated | 0.4910 | N/A |  |
| WT Untreated | 0.0996 | 0.4346 | 0.3243 |
| **μCT<sub>3</sub>** | TS Treated | TS Untreated | WT Treated |
| TS Untreated | 0.3721 |  |  |
| WT Treated | 0.5098 | N/A |  |
| WT Untreated | 0.2033 | 0.9097 | 0.0735 |
| **μCT<sub>4</sub>** | TS Treated | TS Untreated | WT Treated |
| TS Untreated | 0.0573 |  |  |
| WT Treated | 0.2961 | N/A |  |
| WT Untreated | **0.0002** | 0.0430 | **0.0001** |

**Supplementary Table S4.** *P*-values resulting from the pairwise tests after a one-way PERMANOVA (9,999 permutation rounds) based on Mahalanobis distances for tibia microarchitecture parameters. Bold font indicates statistically significant values. Pairwise comparisons marked as N/A were not calculated since they did not evaluate any relevant scientific question.

| **ALL TESTS** | TS Treated | TS Untreated | WT Treated |
| --- | --- | --- | --- |
| TS Untreated | **0.0001** |  |  |
| WT Treated | **0.0002** | N/A |  |
| WT Untreated | **0.0008** | 0.5589 | **0.0089** |
| **TRABECULAR BONE** | TS Treated | TS Untreated | WT Treated |
| TS Untreated | 0.7034 |  |  |
| WT Treated | 0.0792 | N/A |  |
| WT Untreated | 0.8793 | 0.8944 | 0.5405 |
| **CORTICAL BONE STRENGTH** | TS Treated | TS Untreated | WT Treated |
| TS Untreated | **0.0001** |  |  |
| WT Treated | **0.0005** | N/A |  |
| WT Untreated | **0.0002** | 0.4536 | 0.0563 |
| **CORTICAL BONE SIZE** | TS Treated | TS Untreated | WT Treated |
| TS Untreated | **0.0011** |  |  |
| WT Treated | **0.0003** | N/A |  |
| WT Untreated | **0.0007** | 0.0975 | 0.3042 |

**Supplementary Table S5.** *P*-values resulting from the pairwise tests after a one-way PERMANOVA (9,999 permutation rounds) based on Mahalanobis distances for brain volumes before treatment discontinuation. Bold font indicates statistically significant values. Pairwise comparisons marked as N/A were not calculated since they did not evaluate any relevant scientific question.

|  |  |  |  |
| --- | --- | --- | --- |
| **ALL TESTS** | TS Treated | TS Untreated | WT Treated |
| TS Untreated | 0.4921 |  |  |
| WT Treated | **0.0008** | N/A |  |
| WT Untreated | **0.0047** | 0.1064 | 0.4467 |

**Supplementary Table S6.** *P*-values resulting from the pairwise tests after a one-way PERMANOVA (9,999 permutation rounds) based on Mahalanobis distances for brain volumes after treatment discontinuation. Bold font indicates statistically significant values. Pairwise comparisons marked as N/A were not calculated since they did not evaluate any relevant scientific question.

|  |  |  |  |
| --- | --- | --- | --- |
| **ALL TESTS** | TS Treated | TS Untreated | WT Treated |
| TS Untreated | **0.0472** |  |  |
| WT Treated | **0.0001** | N/A |  |
| WT Untreated | 0.0609 | **0.0065** | 0.2808 |

**Supplementary Table S7.** *P*-values resulting from the pairwise tests after a one-way PERMANOVA (9,999 permutation rounds) based on Mahalanobis distances for early neurodevelopmental tests. Bold font indicates statistically significant values. Pairwise comparisons marked as N/A were not calculated since they did not evaluate any relevant scientific question.

| **ALL TESTS** | TS Treated | TS Untreated | WT Treated |
| --- | --- | --- | --- |
| TS Untreated | **0.0001** |  |  |
| WT Treated | **0.0014** | N/A |  |
| WT Untreated | **0.0001** | **0.0001** | 0.1252 |
| **DEV. LANDMARKS** | TS Treated | TS Untreated | WT Treated |
| TS Untreated | 0.0557 |  |  |
| WT Treated | **0.0163** | N/A |  |
| WT Untreated | **0.0181** | 0.8250 | 0.3925 |
| **NEUROMOTOR** | TS Treated | TS Untreated | WT Treated |
| TS Untreated | **0.0065** |  |  |
| WT Treated | **0.0002** | N/A |  |
| WT Untreated | **0.0025** | 0.0555 | 0.3376 |
| **REFLEXES** | TS Treated | TS Untreated | WT Treated |
| TS Untreated | **0.1642** |  |  |
| WT Treated | **0.0001** | N/A |  |
| WT Untreated | **0.0001** | **0.0423** | **0.0001** |

**Supplementary Table S8.**
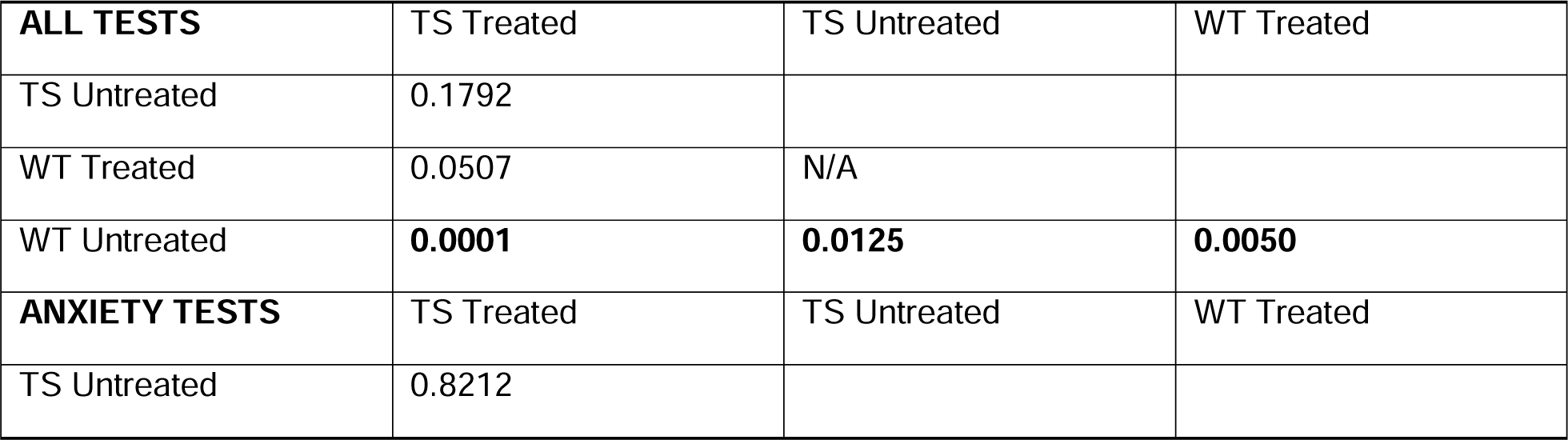
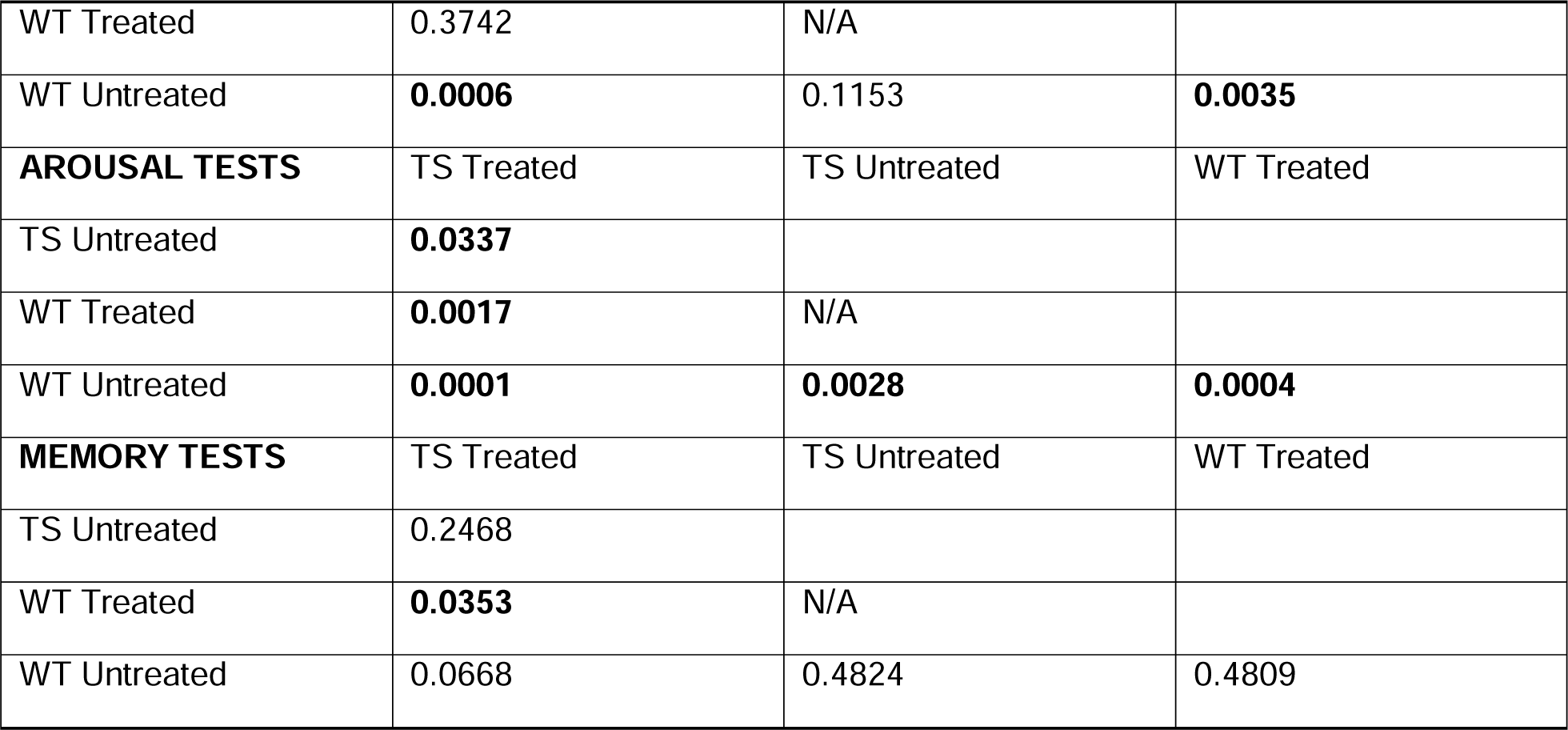
*P*-values resulting from the pairwise tests after a one-way PERMANOVA (9,999 permutation rounds) based on Mahalanobis distances for adult cognitive tests before treatment discontinuation at Cog._1_. Bold font indicates statistically significant values. Pairwise comparisons marked as N/A were not calculated since they did not evaluate any relevant scientific question.

**Supplementary Table S9.**
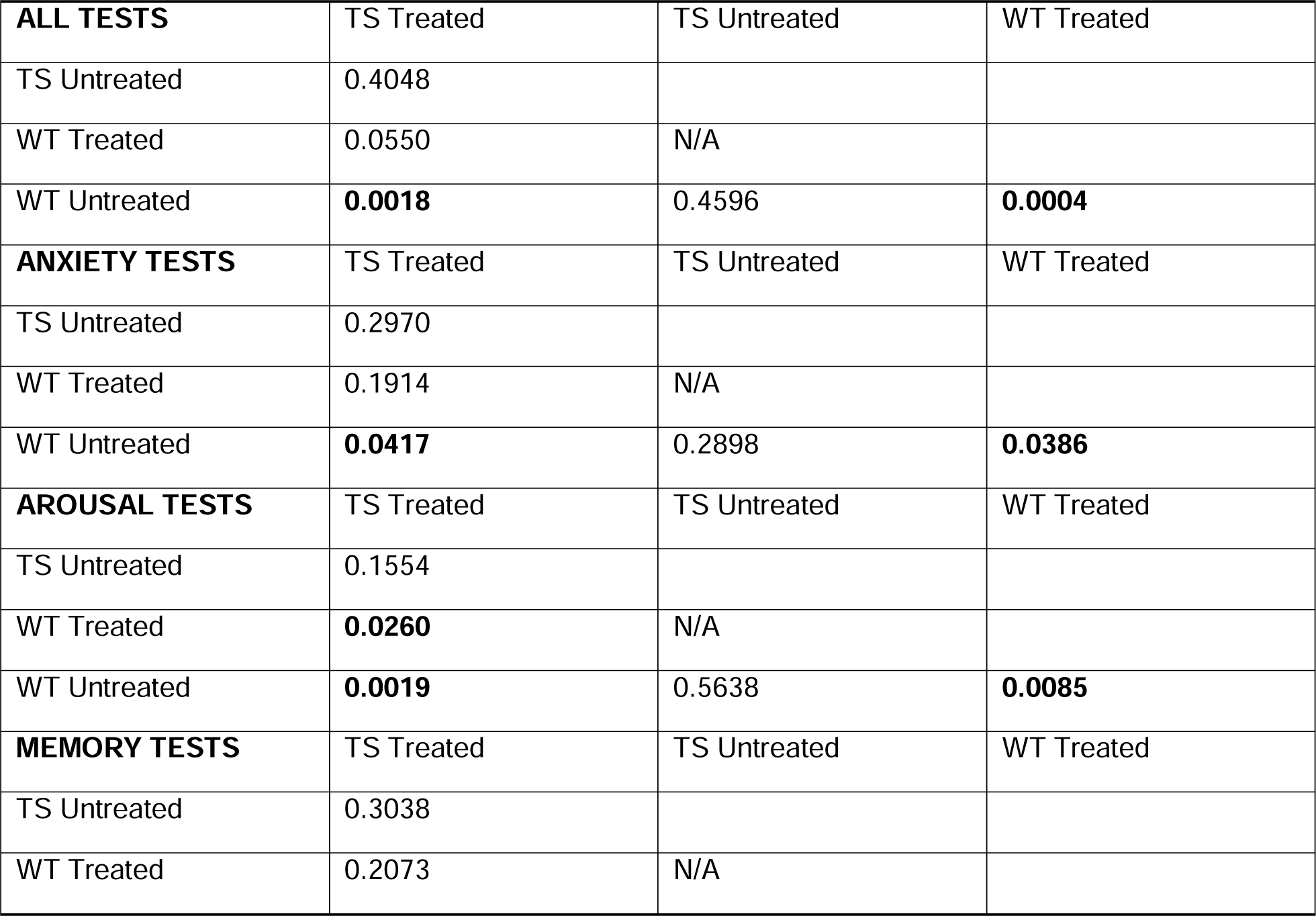
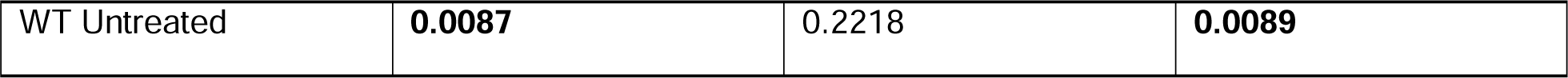
*P*-values resulting from the pairwise tests after a one-way PERMANOVA (9,999 permutation rounds) based on Mahalanobis distances for adult cognitive tests after treatment discontinuation at Cog._2_. Bold font indicates statistically significant values. Pairwise comparisons marked as N/A were not calculated since they did not evaluate any relevant scientific question.

|  |  |  |  |
| --- | --- | --- | --- |
| **ALL TESTS** | TS Treated | TS Untreated | WT Treated |
| TS Untreated | 0.4048 |  |  |
| WT Treated | 0.0550 | N/A |  |
| WT Untreated | **0.0018** | 0.4596 | **0.0004** |
| **ANXIETY TESTS** | TS Treated | TS Untreated | WT Treated |
| TS Untreated | 0.2970 |  |  |
| WT Treated | 0.1914 | N/A |  |
| WT Untreated | **0.0417** | 0.2898 | **0.0386** |
| **AROUSAL TESTS** | TS Treated | TS Untreated | WT Treated |
| TS Untreated | 0.1554 |  |  |
| WT Treated | **0.0260** | N/A |  |
| WT Untreated | **0.0019** | 0.5638 | **0.0085** |
| **MEMORY TESTS** | TS Treated | TS Untreated | WT Treated |
| TS Untreated | 0.3038 |  |  |
| WT Treated | 0.2073 | N/A |  |
| WT Untreated | **0.0087** | 0.2218 | **0.0089** |

**Supplementary Table S10.** Most concentrated metabolites contributing to the respective spectral regions in MRS. ala – alanine, arg – arginine, cre – creatine, cys – cysteine, Hα of AA – α hydrogen of amino acids, ile – iso leucine, leu – leucine, lys – lysine, gaba – γ-amino butyrate, gln – glutamine, glu – glutamane, gly – glycine, GPC – glycerol phosphocholine, his – histidine, NAA – N-acetyl aspartate, PChol – phosphocholine, PCr – phosphocreatine,tyr – tyrosine, val – valine

| Region number | Metabolite name | Integration Range [ppm] |
| --- | --- | --- |
| 1 | lactate, myo-inositol | 4.00 – 4.20 |
| 2 | glucose, Hα of AA, cre, PCr | 3.81 – 4.00 |
| 3 | myo-inositol, glucose, Hα of AA | 3.68 – 3.79 |
| 4 | myo-inositol, gly, glucose | 3.45 – 3.62 |
| 5 | taurine, glucose | 3.30 – 3.45 |
| 6 | PChol, GPC, choline, taurine | 3.19 – 3.30 |
| 7 | cre, PCr, lys, his, cys, tyr | 2.91 – 3.10 |
| 8 | gln, glu, NAA, val, pro | 2.18 – 2.45 |
| 9 | gln, glu, NAA, lys, gaba | 1.88 – 2.18 |
| 10 | ala, leu, lys, arg | 1.45 – 1.70 |
| 11 | lipids, lactate | 1.25 – 1.38 |
| 12 | lipids, ile, leu, val | 0.70 – 0.98 |

**Supplementary Table S11.** *P*-values resulting from the pairwise tests after a one-way PERMANOVA (9,999 permutation rounds) based on Mahalanobis distances for MRS spectra before treatment discontinuation. Bold font indicates statistically significant values. Pairwise comparisons marked as N/A were not calculated since they did not evaluate any relevant scientific question.

| ALL TESTS | TS Treated | TS Untreated | WT Treated |
| --- | --- | --- | --- |
| TS Untreated | **0.0361** |  |  |
| WT Treated | **0.0163** | N/A |  |
| WT Untreated | **0.0496** | **0.0136** | 0.4779 |

**Supplementary Table S12.**
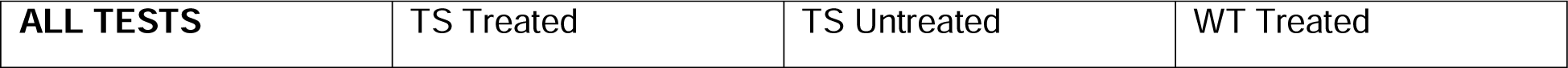
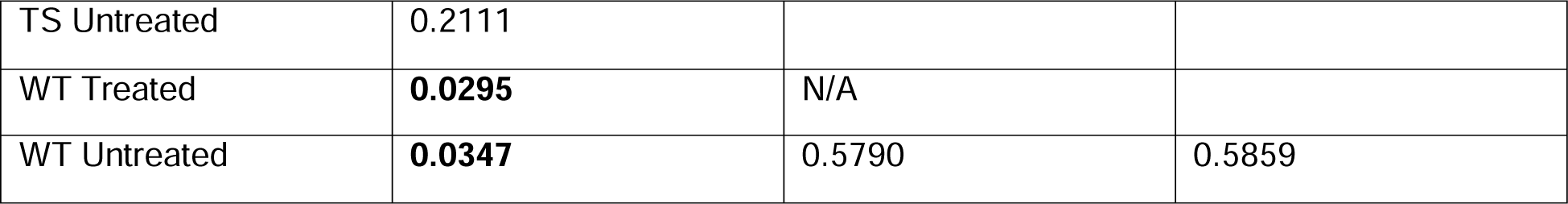
*P*-values resulting from the pairwise tests after a one-way PERMANOVA (9,999 permutation rounds) based on Mahalanobis distances for MRS spectra after treatment discontinuation. Bold font indicates statistically significant values. Pairwise comparisons marked as N/A were not calculated since they did not evaluate any relevant scientific question.

| ALL TESTS | TS Treated | TS Untreated | WT Treated |
| --- | --- | --- | --- |
| TS Untreated | 0.2111 |  |  |
| WT Treated | **0.0295** | N/A |  |
| WT Untreated | **0.0347** | 0.5790 | 0.5859 |

**Supplementary Table S13.** *P*-values resulting from the pairwise tests after a one-way PERMANOVA (9,999 permutation rounds) based on Euclidean distances for normalized gene expression data at endpoint. Bold font indicates statistically significant values. Pairwise comparisons marked as N/A were not calculated since they did not evaluate any relevant scientific question.

|  |  |  |  |
| --- | --- | --- | --- |
| **ALL TESTS** | TS Treated | TS Untreated | WT Treated |
| TS Untreated | 0.2356 |  |  |
| WT Treated | 0.1184 | N/A |  |
| WT Untreated | 0.4473 | 0.1630 | 0.5594 |

**Supplementary Table S15.** Litter information. For each mouse, the litter number, genotype, treatment received and sex are provided.

| Mouse number | Litter | Genotype | Treatment | Sex |
| --- | --- | --- | --- | --- |
| 1 | 1 | TS | Not treated | F |
| 2 | 1 | WT | Not treated | F |
| 3 | 1 | WT | Not treated | M |
| 4 | 1 | TS | Not treated | M |
| 5 | 1 | WT | Not treated | M |
| 6 | 1 | TS | Not treated | F |
| 7 | 2 | TS | Not treated | M |
| 8 | 2 | TS | Not treated | F |
| 9 | 2 | TS | Not treated | M |
| 10 | 2 | TS | Not treated | F |
| 11 | 3 | TS | Not treated | M |
| 12 | 3 | WT | Not treated | F |
| 13 | 4 | WT | Not treated | M |
| 14 | 4 | TS | Not treated | F |
| 15 | 4 | WT | Not treated | F |
| 16 | 4 | WT | Not treated | M |
| 17 | 4 | TS | Not treated | F |
| 18 | 4 | TS | Not treated | F |
| 19 | 5 | WT | Not treated | F |
| 20 | 5 | WT | Not treated | F |
| 21 | 5 | WT | Not treated | F |
| 22 | 5 | WT | Not treated | F |
| 23 | 6 | WT | Not treated | F |
| 24 | 6 | WT | Not treated | M |
| 25 | 6 | TS | Not treated | M |
| 26 | 6 | WT | Not treated | M |
| 27 | 6 | TS | Not treated | F |
| 28 | 6 | TS | Not treated | M |
| 29 | 6 | TS | Not treated | M |
| 30 | 7 | WT | 0.09 mg EGCG/mL | F |
| 31 | 7 | WT | 0.09 mg EGCG/mL | M |
| 32 | 7 | WT | 0.09 mg EGCG/mL | F |
| 33 | 7 | WT | 0.09 mg EGCG/mL | M |
| 34 | 7 | TS | 0.09 mg EGCG/mL | M |
| 35 | 7 | TS | 0.09 mg EGCG/mL | F |
| 36 | 8 | TS | 0.09 mg EGCG/mL | F |
| 37 | 8 | WT | 0.09 mg EGCG/mL | F |
| 38 | 8 | WT | 0.09 mg EGCG/mL | F |
| 39 | 8 | WT | 0.09 mg EGCG/mL | M |
| 40 | 9 | TS | 0.09 mg EGCG/mL | M |
| 41 | 9 | WT | 0.09 mg EGCG/mL | M |
| 42 | 9 | TS | 0.09 mg EGCG/mL | F |
| 43 | 9 | TS | 0.09 mg EGCG/mL | F |
| 44 | 9 | TS | 0.09 mg EGCG/mL | F |
| 45 | 9 | TS | 0.09 mg EGCG/mL | F |
| 46 | 10 | TS | 0.09 mg EGCG/mL | F |
| 47 | 10 | WT | 0.09 mg EGCG/mL | F |
| 48 | 10 | WT | 0.09 mg EGCG/mL | M |
| 49 | 10 | TS | 0.09 mg EGCG/mL | M |
| 50 | 10 | WT | 0.09 mg EGCG/mL | F |
| 51 | 10 | TS | 0.09 mg EGCG/mL | F |
| 52 | 10 | TS | 0.09 mg EGCG/mL | F |
| 53 | 11 | WT | 0.09 mg EGCG/mL | M |
| 54 | 11 | WT | 0.09 mg EGCG/mL | M |
| 55 | 11 | WT | 0.09 mg EGCG/mL | F |
| 56 | 11 | WT | 0.09 mg EGCG/mL | M |
| 57 | 11 | WT | 0.09 mg EGCG/mL | M |
| 58 | 11 | TS | 0.09 mg EGCG/mL | M |
| 59 | 11 | TS | 0.09 mg EGCG/mL | F |
| 60 | 12 | TS | 0.09 mg EGCG/mL | M |
| 61 | 12 | TS | 0.09 mg EGCG/mL | M |
| 62 | 12 | WT | 0.09 mg EGCG/mL | F |
| 63 | 12 | TS | 0.09 mg EGCG/mL | M |
| 64 | 12 | TS | 0.09 mg EGCG/mL | F |
| 65 | 13 | TS | 0.09 mg EGCG/mL | F |
| 66 | 13 | WT | 0.09 mg EGCG/mL | F |
| 67 | 13 | WT | 0.09 mg EGCG/mL | F |
| 68 | 13 | WT | 0.09 mg EGCG/mL | M |
| 69 | 13 | TS | 0.09 mg EGCG/mL | F |
| 70 | 13 | TS | 0.09 mg EGCG/mL | M |
| 71 | 13 | TS | 0.09 mg EGCG/mL | M |

**Supplementary Table S16.** µCT scanning parameters used at each stage.

| Stage | Source energy (Kv) | Filter | Current (μA) | Exposure time (ms) | Average s | Step increment (°) | Total angle (°) | Time | Voxel size ($\mu\text{m}^3$) |
| --- | --- | --- | --- | --- | --- | --- | --- | --- | --- |
| $\mu\text{CT}_1$ | 35 | Al 0.5 mm | 500 | 180 | 3 | 1 | 180° | 3 min | 51.7 |
| $\mu\text{CT}_{2-4}$ | 55 | Al 1 mm | 700 | 80 | 2 | 1 | 360° | 3 min | 51.7 |
| Tibia (*ex vivo*) | 60 | Al 0.5 mm | 80 | 3000 | 0 | 0.4 | 180° | 23 min | 5 |

**Supplementary Table S17.** Anatomical definition of craniofacial landmarks. N/A indicates that the landmarks were not acquired at this stage.

| Landmark number at $\mu\text{CT}_1$ | Landmark number at $\mu\text{CT}_2$, $\mu\text{CT}_3$ and $\mu\text{CT}_4$ | Anatomical definition |
| --- | --- | --- |
| 1 | 1 | Tip of the nasal bone |
| N/A | 2 | Intersection of nasal and frontal bones |
| N/A | 3 | Intersection of frontal and parietal bones |
| N/A | 4 | Intersection of parietal and interparietal bones |
| N/A | 5 | Intersection of interparietal and occipital bones |
| 13 | 6 | Anterior-most point on intersection of premaxillae and nasal bones (left) |
| 14 | 7 | Center of alveolar ridge over maxillary incisor (left) |
| 15 | 8 | Most inferior point on premaxilla-maxilla suture (left) |
| 16 | 9 | Anterior notch on frontal process lateral to infraorbital fissure (left) |
| 17 | 10 | Intersection of frontal process of maxilla with frontal and lacrimal bones (left) |
| 18 | 11 | Intersection of zygomatic process of maxilla with zygoma (left) |
| 19 | 12 | Frontal-squamosal intersection at temporal crest (left) |
| 20 | 13 | Intersection of maxilla and sphenoid on inferior alveolar (left) |
| 21 | 14 | Intersection of zygoma with zygomatic process of temporal (left) |
| N/A | 15 | Intersection of squamosal body to zygomatic process of squamosal (left) |
| N/A | 16 | Intersection of parietal, temporal and occipital bones (left) |
| 24 | 17 | Anterior-most point on intersection of premaxillae and nasal bones (right) |
| 25 | 18 | Center of alveolar ridge over maxillary incisor (right) |
| 26 | 19 | Most inferior point on premaxilla-maxilla suture (right) |
| 27 | 20 | Anterior notch on frontal process lateral to infraorbital fissure (right) |
| 28 | 21 | Intersection of frontal process of maxilla with frontal and lacrimal bones (right) |
| 29 | 22 | Intersection of zygomatic process of maxilla with zygoma (right) |
| 30 | 23 | Frontal-squamosal intersection at temporal crest (right) |
| 31 | 24 | Intersection of maxilla and sphenoid on inferior alveolar (right) |
| 32 | 25 | Intersection of zygoma with zygomatic process of temporal (right) |
| N/A | 26 | Intersection of squamosal body to zygomatic process of squamosal (right) |
| N/A | 27 | Intersection of parietal, temporal and occipital bones (right) |
| 2 | N/A | Posterior-most point on the nasal bone at the midline |
| 3 | N/A | Anterior-most point on the frontal bone at the midline (left) |
| 4 | N/A | Anterior-most point on the frontal bone at the midline (right) |
| 5 | N/A | Posterior-most point on the frontal bone at the midline (left) |
| 6 | N/A | Posterior-most point on the frontal bone at the midline (right) |
| 7 | N/A | Anterior-most point on the parietal bone at the midline (left) |
| 8 | N/A | Anterior-most point on the parietal bone at the midline (right) |
| 9 | N/A | Maximum curvature point on the posterior part of the parietal bone (left) |
| 10 | N/A | Maximum curvature point on the posterior part of the parietal bone (right) |
| 11 | N/A | Anterior-most point on the interparietal bone at the midline |
| 12 | N/A | Anterior-most point on the occipital bone at the midline |
| 22 | N/A | Joining of squamosal body to zygomatic process of squamosal (left) |
| 23 | N/A | Tip of the post-tympanic hook (left) |
| 33 | N/A | Joining of squamosal body to zygomatic process of squamosal (right) |
| 34 | N/A | Tip of the post-tympanic hook (right) |

**Supplementary Table S18.** Trabecular bone parameters measured from *ex vivo* micro-CT images at 8M.

| Parameter | Abbreviation | Description | Unit |
| --- | --- | --- | --- |
| Percentage bone volume | BV/TV | Ratio of bone volume to total volume inside the Region of interest | % |
| Trabecular number | Tb.N | Average number of trabeculae per length unit of bone | 1/mm |
| Trabecular thickness | Tb.Th | Average thickness of the trabecular bone | mm |
| Trabecular separation | Tb.Sp | Average distance between the trabecular bone | mm |
| Bone mineral density | BMD | Volumetric density of hydroxyapatite in the trabecular bone | g/cm<sup>3</sup> |

**Supplementary Table S19.**
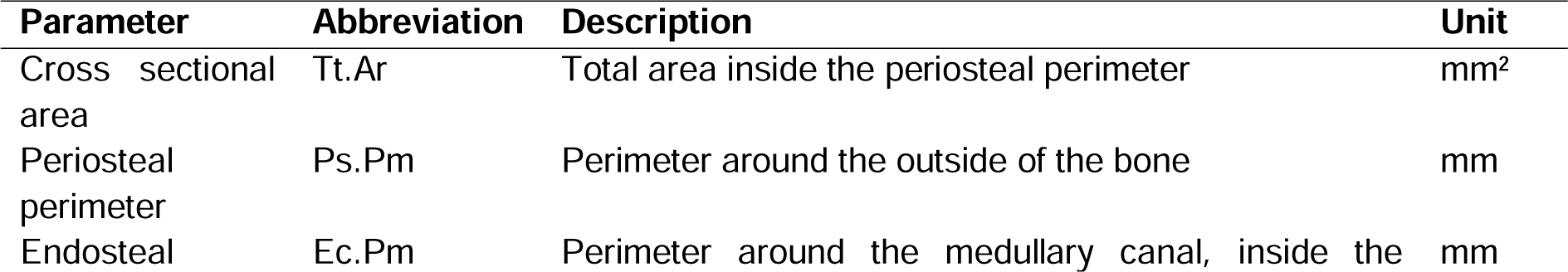
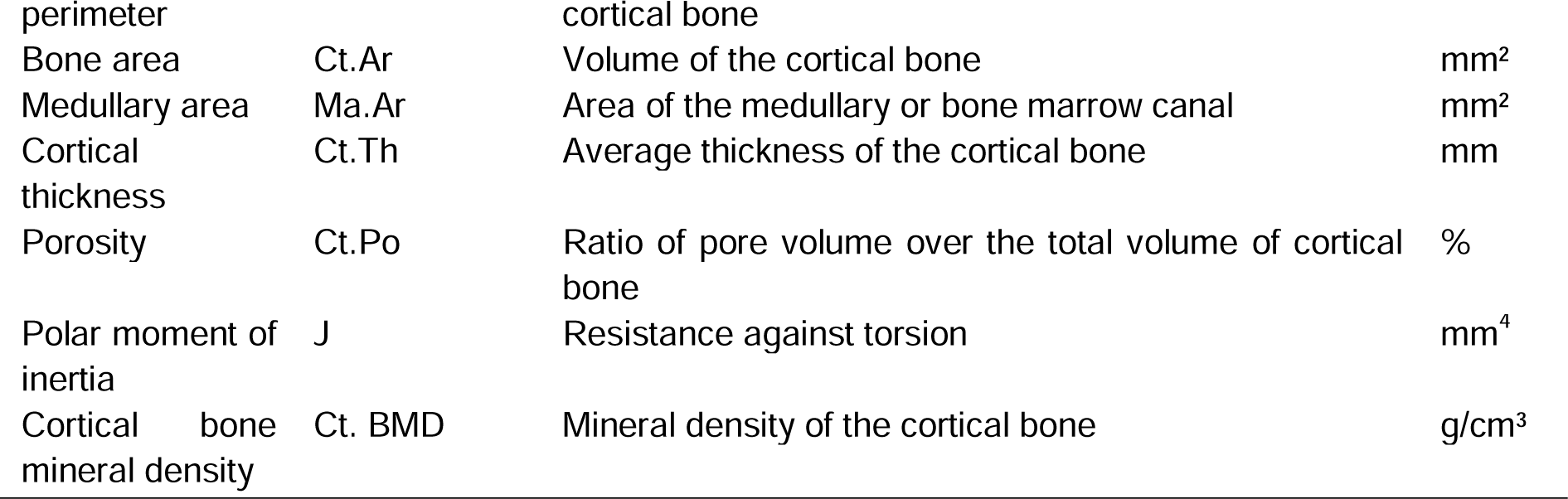
Cortical bone parameters measured from *ex vivo* micro-CT images at 8M.

**Supplementary Table S20.**
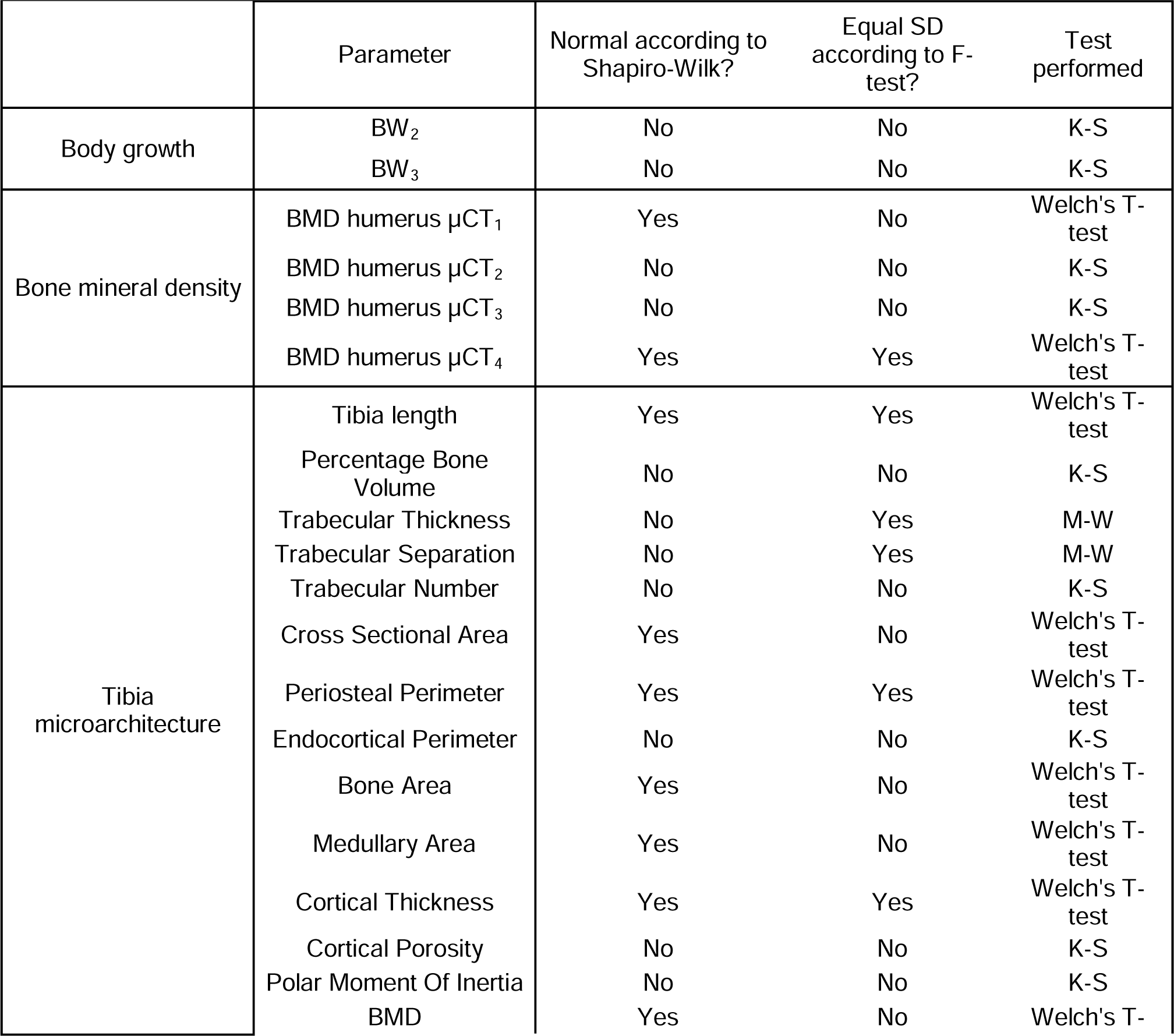
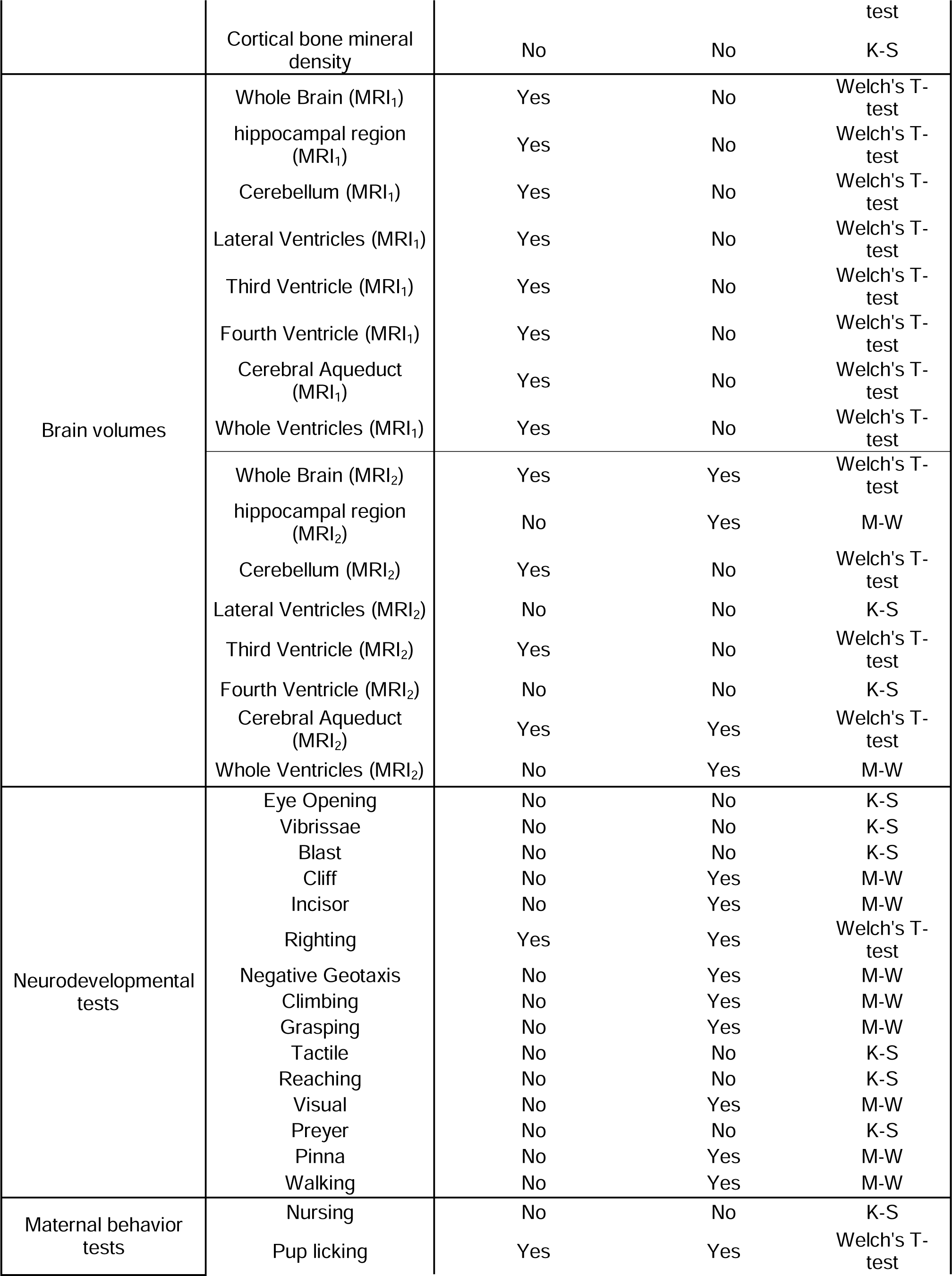
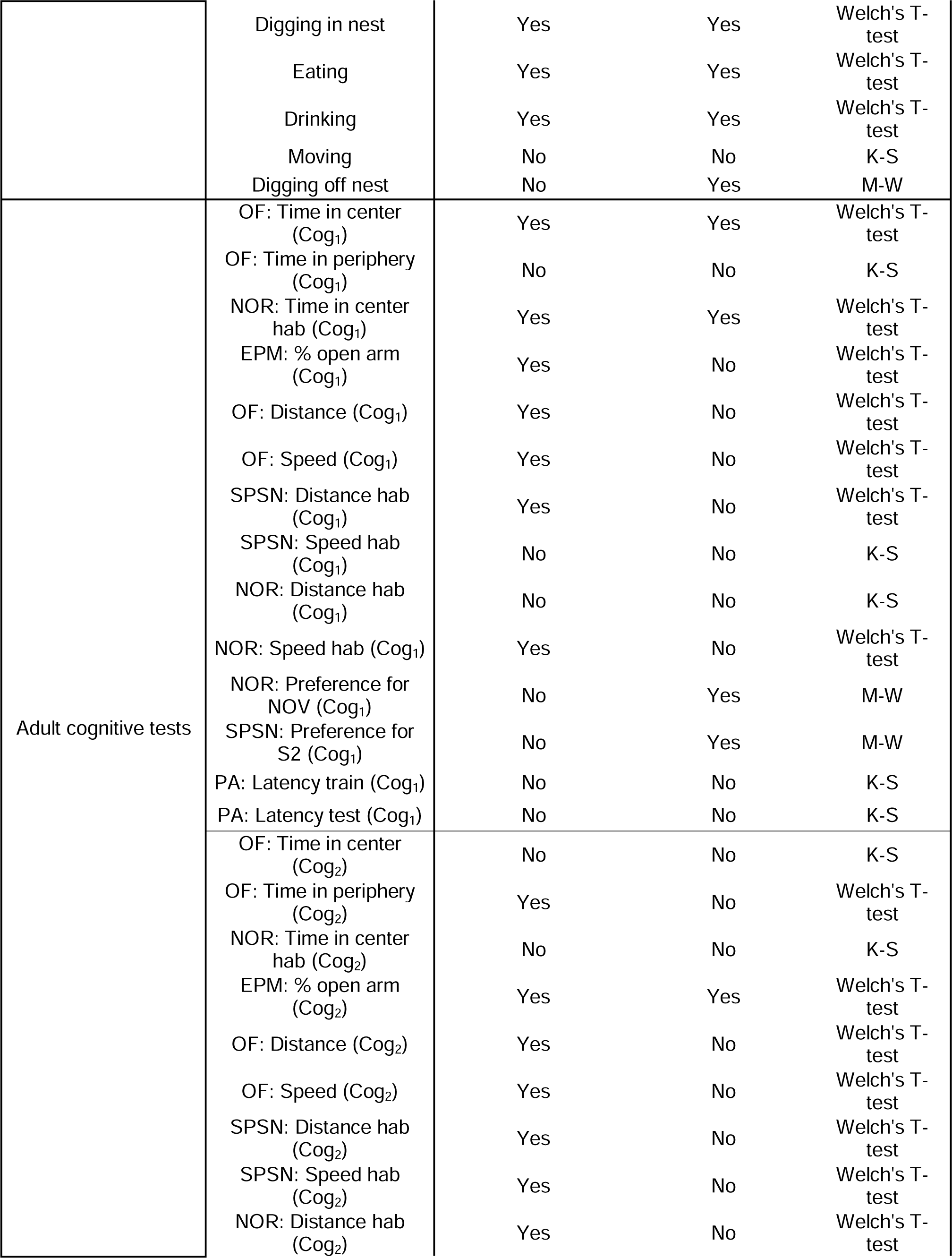
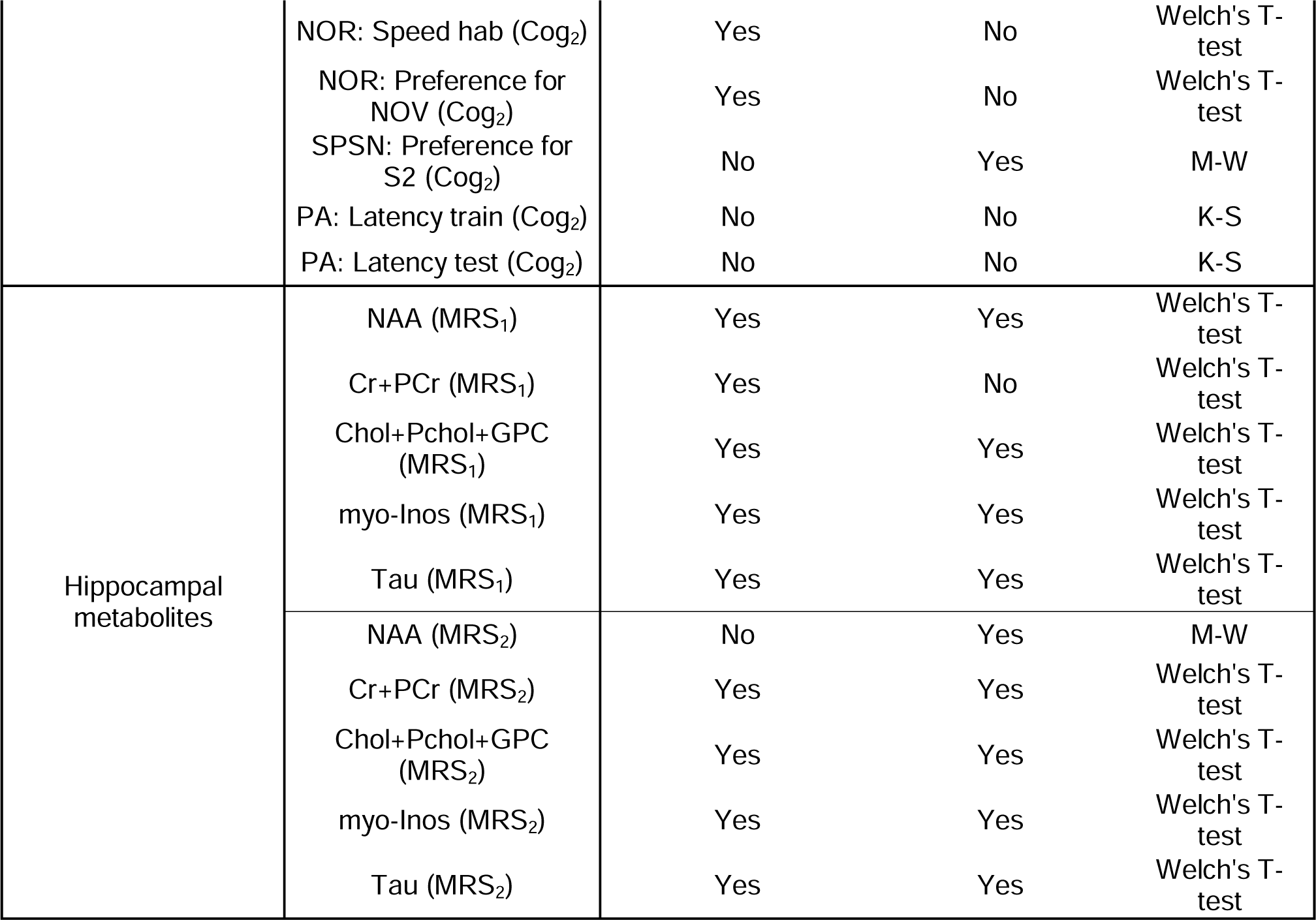
Normality, homoscedasticity, and statistical tests performed per parameter. If one of the four mice groups was not normally distributed or one pairwise comparison was not homoscedastic, the variable was considered as not normally distributed and/or not homoscedastic.

## Author Contributions

Conceptualization, SL,NMA,GVV; methodology, LVe,WG,GVV; formal analysis, SL,ES,MA,AC,MP,KW,VVB,LVa,NGF,LGF; investigation, SL,BT,ES; resources, NMA,GVV,ZCV,CA,UH,HP; data curation, SL,NMA, GVV; writing—original draft preparation, SL,NMA,GVV; writing—review and editing, SL,NMA,GVV; visualization, SL,NMA,GVV; supervision, NMA,GVV,ZCV,CA; project administration, GVV; funding acquisition, NMA and GVV. All authors have read and agreed to the published version of the manuscript.

## Institutional Review Board Statement

The animal study protocol was approved by the Animal Ethics Committee of KU Leuven (ECD approval number P120/2019).

## Data Availability Statement

Data is available upon reasonable request to the corresponding authors.

## Acknowledgments

Imaging data was acquired in the Molecular Small Animal Imaging Center (MoSAIC), a core facility of Dept. Imaging and Pathology, Group Biomedical Sciences, KU Leuven. RNAseq was pefomed in the Genomics Core facility of KU Leuven. The authors acknowledge the Laboratory Animal Centre core facility of KU Leuven for support with animal care.

## REFERENCES

Abeysekera, I., Thomas, J., Georgiadis, T. M., Berman, A. G., Hammond, M. A., Dria, K. J., Roper, R. J. (2016). Differential effects of Epigallocatechin-3-gallate containing supplements on correcting skeletal defects in a Down syndrome mouse model. Mol Nutr Food Res, 60(4), 717–726. 10.1002/mnfr.201500781

Aït Yahya-Graison, E., Aubert, J., Dauphinot, L., Rivals, I., Prieur, M., Golfier, G., Potier, M. C. (2007). Classification of human chromosome 21 gene-expression variations in Down syndrome: impact on disease phenotypes. Am J Hum Genet, 81(3), 475–491. 10.1086/520000

Aldridge, K., Reeves, R. H., Olson, L. E., & Richtsmeier, J. T. (2007). Differential effects of trisomy on brain shape and volume in related aneuploid mouse models. American journal of medical genetics. Part A, 143A(10), 1060–1070. 10.1002/ajmg.a.31721

Anders, S., Pyl, P. T., & Huber, W. (2014). HTSeq—a Python framework to work with high-throughput sequencing data. Bioinformatics, 31(2), 166–169. 10.1093/bioinformatics/btu638

Andrews, S. (2010). FASTQC. A quality control tool for high throughput sequence data. In.

Antonarakis, S. E., Lyle, R., Dermitzakis, E. T., Reymond, A., & Deutsch, S. (2004). Chromosome 21 and Down syndrome: from genomics to pathophysiology. Nature Reviews Genetics, 5(10), 725–738. 10.1038/nrg1448

Antonarakis, S. E., Skotko, B. G., Rafii, M. S., Strydom, A., Pape, S. E., Bianchi, D. W., Reeves, R. H. (2020). Down syndrome. Nature Reviews Disease Primers, 6(1), 9. 10.1038/s41572-019-0143-7

Aronesty, E. (2011). ea-utils: Command-line tools for processing biological sequencing data. In: Durham, NC.

Arron, J. R., Winslow, M. M., Polleri, A., Chang, C.-P., Wu, H., Gao, X., Crabtree, G. R. (2006). NFAT dysregulation by increased dosage of DSCR1 and DYRK1A on chromosome 21. Nature, 441(7093), 595–600. 10.1038/nature04678

Atas-Ozcan, H., Brault, V., Duchon, A., & Herault, Y. (2021). Dyrk1a from Gene Function in Development and Physiology to Dosage Correction across Life Span in Down Syndrome. Genes, 12(11), 1833. https://www.mdpi.com/2073-4425/12/11/1833

Aylward, E. H., Habbak, R., Warren, A. C., Pulsifer, M. B., Barta, P. E., Jerram, M., & Pearlson, G. D. (1997). Cerebellar volume in adults with Down syndrome. Arch Neurol, 54(2), 209–212. 10.1001/archneur.1997.00550140077016

Aziz, N. M., Guedj, F., Pennings, J. L. A., Olmos-Serrano, J. L., Siegel, A., Haydar, T. F., & Bianchi, D. W. (2018). Lifespan analysis of brain development, gene expression and behavioral phenotypes in the Ts1Cje, Ts65Dn and Dp(16)1/Yey mouse models of Down syndrome. Disease Models &&& Mechanisms, 11(6), dmm031013. 10.1242/dmm.031013

Beacher, F., Simmons, A., Daly, E., Prasher, V., Adams, C., Margallo-Lana, M. L., Murphy, D. G. M. (2005). Hippocampal Myo-inositol and Cognitive Ability in Adults With Down Syndrome: An In Vivo Proton Magnetic Resonance Spectroscopy Study. Archives of General Psychiatry, 62(12), 1360–1365. 10.1001/archpsyc.62.12.1360

Becker, W., Soppa, U., & Tejedor, F. J. (2014). DYRK1A: a potential drug target for multiple Down syndrome neuropathologies. CNS Neurol Disord Drug Targets, 13(1), 26–33. 10.2174/18715273113126660186

Beqaj, S., Jusaj, N., & Živković, V. (2017). Attainment of gross motor milestones in children with Down syndrome in Kosovo - developmental perspective. Med Glas (Zenica), 14(2), 189–198. 10.17392/917-17

Billingsley, C. N., Allen, J. R., Baumann, D. D., Deitz, S. L., Blazek, J. D., Newbauer, A., Roper, R. J. (2013). Non-trisomic homeobox gene expression during craniofacial development in the Ts65Dn mouse model of Down syndrome. Am J Med Genet A, 161a(8), 1866–1874. 10.1002/ajmg.a.36006

Blazek, J. D., Abeysekera, I., Li, J., & Roper, R. J. (2015). Rescue of the abnormal skeletal phenotype in Ts65Dn Down syndrome mice using genetic and therapeutic modulation of trisomic Dyrk1a. Hum Mol Genet, 24(20), 5687–5696. 10.1093/hmg/ddv284

Blazek, J. D., Gaddy, A., Meyer, R., Roper, R. J., & Li, J. (2011). Disruption of bone development and homeostasis by trisomy in Ts65Dn Down syndrome mice. Bone, 48(2), 275–280. 10.1016/j.bone.2010.09.028

Blazek, J. D., Malik, A. M., Tischbein, M., Arbones, M. L., Moore, C. S., & Roper, R. J. (2015). Abnormal mineralization of the Ts65Dn Down syndrome mouse appendicular skeleton begins during embryonic development in a Dyrk1a-independent manner. Mechanisms of Development, 136, 133–142. 10.1016/j.mod.2014.12.004

Caputo, C., Wood, E., & Jabbour, L. (2016). Impact of fetal alcohol exposure on body systems: A systematic review. Birth Defects Res C Embryo Today, 108(2), 174–180. 10.1002/bdrc.21129

Carfì, A., Liperoti, R., Fusco, D., Giovannini, S., Brandi, V., Vetrano, D. L., Onder, G. (2017). Bone mineral density in adults with Down syndrome. Osteoporos Int, 28(10), 2929–2934. 10.1007/s00198-017-4133-x

Catuara-Solarz, S., Espinosa-Carrasco, J., Erb, I., Langohr, K., Gonzalez, J. R., Notredame, C., & Dierssen, M. (2016). Combined Treatment With Environmental Enrichment and (-)-Epigallocatechin-3-Gallate Ameliorates Learning Deficits and Hippocampal Alterations in a Mouse Model of Down Syndrome. eNeuro, 3(5). 10.1523/eneuro.0103-16.2016

Catuara-Solarz, S., Espinosa-Carrasco, J., Erb, I., Langohr, K., Notredame, C., Gonzalez, J. R., & Dierssen, M. (2015). Principal Component Analysis of the Effects of Environmental Enrichment and (-)- epigallocatechin-3-gallate on Age-Associated Learning Deficits in a Mouse Model of Down Syndrome [Original Research]. Frontiers in Behavioral Neuroscience, 9(330). 10.3389/fnbeh.2015.00330

Chang, K. T., Shi, Y. J., & Min, K. T. (2003). The Drosophila homolog of Down’s syndrome critical region 1 gene regulates learning: implications for mental retardation. Proc Natl Acad Sci U S A, 100(26), 15794–15799. 10.1073/pnas.2536696100

Chang, P., Bush, D., Schorge, S., Good, M., Canonica, T., Shing, N., Fisher, E. M. C. (2020). Altered Hippocampal-Prefrontal Neural Dynamics in Mouse Models of Down Syndrome. Cell Reports, 30(4), 1152–1163.e1154. 10.1016/j.celrep.2019.12.065

Chrast, R., Scott, H. S., Papasavvas, M. P., Rossier, C., Antonarakis, E. S., Barras, C., Antonarakis, S. E. (2000). The Mouse Brain Transcriptome by SAGE: Differences in Gene Expression between P30 Brains of the Partial Trisomy 16 Mouse Model of Down Syndrome (Ts65Dn) and Normals. Genome Research, 10(12), 2006–2021. http://genome.cshlp.org/content/10/12/2006.abstract

Chu, K. O., Wang, C. C., Chu, C. Y., Chan, K. P., Rogers, M. S., Choy, K. W., & Pang, C. P. (2006). Pharmacokinetic studies of green tea catechins in maternal plasma and fetuses in rats. J Pharm Sci, 95(6), 1372–1381. 10.1002/jps.20594

Chu, K. O., Wang, C. C., Chu, C. Y., Choy, K. W., Pang, C. P., & Rogers, M. S. (2006). Uptake and distribution of catechins in fetal organs following in utero exposure in rats. Human Reproduction, 22(1), 280–287. 10.1093/humrep/del353

Cieuta-Walti, C., Cuenca-Royo, A., Langohr, K., Rakic, C., López-Vílchez, M., Lirio, J., Fornell, R. T. (2022). Safety and preliminary efficacy on cognitive performance and adaptive functionality of epigallocatechin gallate (EGCG) in children with Down syndrome. A randomized phase Ib clinical trial (PERSEUS study). Genet Med, 24(10), 2004–2013. 10.1016/j.gim.2022.06.011

Costa, A. C., Stasko, M. R., Schmidt, C., & Davisson, M. T. (2010). Behavioral validation of the Ts65Dn mouse model for Down syndrome of a genetic background free of the retinal degeneration mutation Pde6b(rd1). Behav Brain Res, 206(1), 52–62. 10.1016/j.bbr.2009.08.034

Coussons-Read, M. E., & Crnic, L. S. (1996). Behavioral assessment of the Ts65Dn mouse, a model for Down syndrome: altered behavior in the elevated plus maze and open field. Behav Genet, 26(1), 7–13.

Davisson, M. T., Schmidt, C., & Akeson, E. C. (1990). Segmental trisomy of murine chromosome 16: a new model system for studying Down syndrome. Prog Clin Biol Res, 360, 263–280.

de la Torre, R., de Sola, S., Hernandez, G., Farré, M., Pujol, J., Rodriguez, J., Dierssen, M. (2016). Safety and efficacy of cognitive training plus epigallocatechin-3-gallate in young adults with Down’s syndrome (TESDAD): a double-blind, randomised, placebo-controlled, phase 2 trial. The Lancet Neurology, 15(8), 801–810. 10.1016/S1474-4422(16)30034-5

De la Torre, R., De Sola, S., Pons, M., Duchon, A., de Lagran, M. M., Farré, M., Dierssen, M. (2014). Epigallocatechin-3-gallate, a DYRK1A inhibitor, rescues cognitive deficits in Down syndrome mouse models and in humans. Mol Nutr Food Res, 58(2), 278–288. 10.1002/mnfr.201300325

de Moraes, M. E., Tanaka, J. L., de Moraes, L. C., Filho, E. M., & de Melo Castilho, J. C. (2008). Skeletal age of individuals with Down syndrome. Spec Care Dentist, 28(3), 101–106. 10.1111/j.1754-4505.2008.00020.x

De Toma, I., Sierra, C., & Dierssen, M. (2021). Meta-analysis of transcriptomic data reveals clusters of consistently deregulated gene and disease ontologies in Down syndrome. PLoS Comput Biol, 17(9), e1009317. 10.1371/journal.pcbi.1009317

Dierssen, M., & de Lagrán, M. M. (2006). DYRK1A (dual-specificity tyrosine-phosphorylated and - regulated kinase 1A): a gene with dosage effect during development and neurogenesis. ScientificWorldJournal, 6, 1911–1922. 10.1100/tsw.2006.319

Dierssen, M., Fotaki, V., Martínez de Lagrán, M., Gratacós, M., Arbonés, M., Fillat, C., & Estivill, X. (2002). Neurobehavioral development of two mouse lines commonly used in transgenic studies. Pharmacol Biochem Behav, 73(1), 19–25. 10.1016/s0091-3057(02)00792-x

Dryden, I. L., & Mardia, K. V. (1998). Statistical Shape Analysis. Wiley.

Duchon, A., del Mar Muniz Moreno, M., Martin Lorenzo, S., Silva de Souza, M. P., Chevalier, C., Nalesso, V., Herault, Y. (2021). Multi-influential genetic interactions alter behaviour and cognition through six main biological cascades in Down syndrome mouse models. Hum Mol Genet, 30(9), 771–788. 10.1093/hmg/ddab012

Duchon, A., Raveau, M., Chevalier, C., Nalesso, V., Sharp, A. J., & Herault, Y. (2011). Identification of the translocation breakpoints in the Ts65Dn and Ts1Cje mouse lines: relevance for modeling Down syndrome. Mammalian genome : official journal of the International Mammalian Genome Society, 22(11-12), 674–684. 10.1007/s00335-011-9356-0

Escorihuela, R. M., Fernández-Teruel, A., Vallina, I. F., Baamonde, C., Lumbreras, M. A., Dierssen, M., Flórez, J. (1995). A behavioral assessment of Ts65Dn mice: a putative Down syndrome model. Neuroscience Letters, 199(2), 143–146. 10.1016/0304-3940(95)12052-6

Escorihuela, R. M., Vallina, I. F., Martínez-Cué, C., Baamonde, C., Dierssen, M., Tobeña, A., Fernández-Teruel, A. (1998). Impaired short- and long-term memory in Ts65Dn mice, a model for Down syndrome. Neuroscience Letters, 247(2), 171–174. 10.1016/S0304-3940(98)00317-6

Fedorov, A., Beichel, R., Kalpathy-Cramer, J., Finet, J., Fillion-Robin, J. C., Pujol, S., Kikinis, R. (2012). 3D Slicer as an image computing platform for the Quantitative Imaging Network. Magn Reson Imaging, 30(9), 1323–1341. 10.1016/j.mri.2012.05.001

Fernandes, M. B., Maximino, L. P., Perosa, G. B., Abramides, D. V., Passos-Bueno, M. R., & Yacubian-Fernandes, A. (2016). Apert and Crouzon syndromes-Cognitive development, brain abnormalities, and molecular aspects. Am J Med Genet A, 170(6), 1532–1537. 10.1002/ajmg.a.37640

Ferreira-Vasques, A. T., & Lamônica, D. A. C. (2015). Motor, linguistic, personal and social aspects of children with Down syndrome. Journal of applied oral science : revista FOB, 23(4), 424–430. 10.1590/1678-775720150102

Fischer-Brandies, H., Schmid, R. G., & Fischer-Brandies, E. (1986). Craniofacial development in patients with Down’s syndrome from birth to 14 years of age. European Journal of Orthodontics, 8(1), 35–42. 10.1093/ejo/8.1.35

Frank, K., & Esbensen, A. J. (2015). Fine motor and self-care milestones for individuals with Down syndrome using a Retrospective Chart Review. Journal of Intellectual Disability Research, 59(8), 719–729. 10.1111/jir.12176

García-Cerro, S., Martínez, P., Vidal, V., Corrales, A., Flórez, J., Vidal, R., Martínez-Cué, C. (2014). Overexpression of Dyrk1A Is Implicated in Several Cognitive, Electrophysiological and Neuromorphological Alterations Found in a Mouse Model of Down Syndrome. PLoS One, 9(9), e106572. 10.1371/journal.pone.0106572

Goodlett, C. R., Stringer, M., LaCombe, J., Patel, R., Wallace, J. M., & Roper, R. J. (2020). Evaluation of the therapeutic potential of Epigallocatechin-3-gallate (EGCG) via oral gavage in young adult Down syndrome mice. Scientific Reports, 10(1), 10426–10426. 10.1038/s41598-020-67133-z

Grieco, J., Pulsifer, M., Seligsohn, K., Skotko, B., & Schwartz, A. (2015). Down syndrome: Cognitive and behavioral functioning across the lifespan. Am J Med Genet C Semin Med Genet, 169(2), 135–149. 10.1002/ajmg.c.31439

Griffey, R. H., & P. Flamig, D. (1990). VAPOR for solvent-suppressed, short-echo, volume-localized proton spectroscopy. Journal of Magnetic Resonance (1969), 88(1), 161-166. 10.1016/0022-2364(90)90120-X

Guedj, F., Sébrié, C., Rivals, I., Ledru, A., Paly, E., Bizot, J. C., Delabar, J. M. (2009). Green tea polyphenols rescue of brain defects induced by overexpression of DYRK1A. PLoS One, 4(2), e4606. 10.1371/journal.pone.0004606

Guidi, S., Ciani, E., Bonasoni, P., Santini, D., & Bartesaghi, R. (2011). Widespread proliferation impairment and hypocellularity in the cerebellum of fetuses with down syndrome. Brain Pathol, 21(4), 361–373. 10.1111/j.1750-3639.2010.00459.x

Gupta, M., Dhanasekaran, A. R., & Gardiner, K. J. (2016). Mouse models of Down syndrome: gene content and consequences. Mammalian Genome, 27(11), 538–555. 10.1007/s00335-016-9661-8

Hallgrimsson, B., Percival, C. J., Green, R., Young, N. M., Mio, W., & Marcucio, R. (2015). Morphometrics, 3D Imaging, and Craniofacial Development. Curr Top Dev Biol, 115, 561–597. 10.1016/bs.ctdb.2015.09.003

Hammer, O., Harper, D., & Ryan, P. (2001). PAST: Paleontological Statistics Software Package for Education and Data Analysis. Palaeontologia Electronica, 4, 1–9.

Hamner, T., Udhnani, M. D., Osipowicz, K. Z., & Lee, N. R. (2018). Pediatric Brain Development in Down Syndrome: A Field in Its Infancy. J Int Neuropsychol Soc, 24(9), 966–976. 10.1017/s1355617718000206

Heinen, M., Hettich, M. M., Ryan, D. P., Schnell, S., Paesler, K., & Ehninger, D. (2012). Adult-Onset Fluoxetine Treatment Does Not Improve Behavioral Impairments and May Have Adverse Effects on the Ts65Dn Mouse Model of Down Syndrome. Neural Plasticity, 2012, 467251. 10.1155/2012/467251

Herault, Y., Delabar, J. M., Fisher, E. M. C., Tybulewicz, V. L. J., Yu, E., & Brault, V. (2017). Rodent models in Down syndrome research: impact and future opportunities. Dis Model Mech, 10(10), 1165–1186. 10.1242/dmm.029728

Hoeffer, C. A., Dey, A., Sachan, N., Wong, H., Patterson, R. J., Shelton, J. M., Rothermel, B. A. (2007). The Down syndrome critical region protein RCAN1 regulates long-term potentiation and memory via inhibition of phosphatase signaling. J Neurosci, 27(48), 13161–13172. 10.1523/jneurosci.3974-07.2007

Holtzman, D. M., Santucci, D., Kilbridge, J., Chua-Couzens, J., Fontana, D. J., Daniels, S. E., Mobley, W. C. (1996). Developmental abnormalities and age-related neurodegeneration in a mouse model of1Down1syndrome. Proceedings of the National Academy of Sciences, 93(23), 13333–13338. 10.1073/pnas.93.23.13333

Huang, H. T., Cheng, T. L., Lin, S. Y., Ho, C. J., Chyu, J. Y., Yang, R. S., Shen, C. L. (2020). Osteoprotective Roles of Green Tea Catechins. Antioxidants (Basel, Switzerland), 9(11). 10.3390/antiox9111136

Huang, W., Galdzicki, Z., van Gelderen, P., Balbo, A., Chikhale, E. G., Schapiro, M. B., & Rapoport, S. I. (2000). Brain myo-inositol level is elevated in Ts65Dn mouse and reduced after lithium treatment. NeuroReport, 11(3), 445–448. https://journals.lww.com/neuroreport/Fulltext/2000/02280/Brain_myo_inositol_level_is_elevated_in_Ts65Dn.4.aspx

Inc., T. M. (2022). MATLAB 2020b. In The MathWorks Inc. https://www.mathworks.com

Insausti, A. M., Megías, M., Crespo, D., Cruz-Orive, L. M., Dierssen, M., Vallina, I. F., Flórez, J. (1998). Hippocampal volume and neuronal number in Ts65Dn mice: a murine model of Down syndrome. Neurosci Lett, 253(3), 175–178. 10.1016/s0304-3940(98)00641-7

Ishihara, K., Amano, K., Takaki, E., Shimohata, A., Sago, H., Epstein, C. J., & Yamakawa, K. (2010). Enlarged brain ventricles and impaired neurogenesis in the Ts1Cje and Ts2Cje mouse models of Down syndrome. Cereb Cortex, 20(5), 1131–1143. 10.1093/cercor/bhp176

Jamal, R., LaCombe, J., Patel, R., Blackwell, M., Thomas, J. R., Sloan, K., Roper, R. J. (2022). Increased dosage and treatment time of Epigallocatechin-3-gallate (EGCG) negatively affects skeletal parameters in normal mice and Down syndrome mouse models. PLoS One, 17(2), e0264254. 10.1371/journal.pone.0264254

James Rohlf, F., & Marcus, L. F. (1993). A revolution morphometrics. Trends Ecol Evol, 8(4), 129–132. 10.1016/0169-5347(93)90024-J

Jarhad, D. B., Mashelkar, K. K., Kim, H.-R., Noh, M., & Jeong, L. S. (2018). Dual-Specificity Tyrosine Phosphorylation-Regulated Kinase 1A (DYRK1A) Inhibitors as Potential Therapeutics. Journal of Medicinal Chemistry, 61(22), 9791–9810. 10.1021/acs.jmedchem.8b00185

Ji, J., Lee, H., Argiropoulos, B., Dorrani, N., Mann, J., Martinez-Agosto, J. A., Quintero-Rivera, F. (2015). DYRK1A haploinsufficiency causes a new recognizable syndrome with microcephaly, intellectual disability, speech impairment, and distinct facies. European Journal of Human Genetics, 23, 1473–1481.

Kao, C. H., Chen, C. C., Wang, S. J., & Yeh, S. H. (1992). Bone mineral density in children with Down’s syndrome detected by dual photon absorptiometry. Nucl Med Commun, 13(10), 773–775.

Kazemi, M., Salehi, M., & Kheirollahi, M. (2016). Down Syndrome: Current Status, Challenges and Future Perspectives. Int J Mol Cell Med, 5(3), 125–133.

Kazuki, Y., Gao, F. J., Li, Y., Moyer, A. J., Devenney, B., Hiramatsu, K., Reeves, R. H. (2020). A non-mosaic transchromosomic mouse model of down syndrome carrying the long arm of human chromosome 21. eLife, 9. 10.7554/eLife.56223

Kazuki, Y., Gao, F. J., Yamakawa, M., Hirabayashi, M., Kazuki, K., Kajitani, N., Reeves, R. H. (2022). A transchromosomic rat model with human chromosome 21 shows robust Down syndrome features. The American Journal of Human Genetics, 109(2), 328–344. 10.1016/j.ajhg.2021.12.015

Keeling, J. W., Hansen, B. F., & Kjaer, I. (1997). Pattern of malformations in the axial skeleton in human trisomy 21 fetuses. Am J Med Genet, 68(4), 466–471.

Kim, D., Paggi, J. M., Park, C., Bennett, C., & Salzberg, S. L. (2019). Graph-based genome alignment and genotyping with HISAT2 and HISAT-genotype. Nat Biotechnol, 37(8), 907–915. 10.1038/s41587-019-0201-4

Kim, H. I., Kim, S. W., Kim, J., Jeon, H. R., & Jung, D. W. (2017). Motor and Cognitive Developmental Profiles in Children With Down Syndrome. Annals of rehabilitation medicine, 41(1), 97–103. 10.5535/arm.2017.41.1.97

Kleschevnikov, A. M., Yu, J., Kim, J., Lysenko, L. V., Zeng, Z., Yu, Y. E., & Mobley, W. C. (2017). Evidence that increased Kcnj6 gene dose is necessary for deficits in behavior and dentate gyrus synaptic plasticity in the Ts65Dn mouse model of Down syndrome. Neurobiol Dis, 103, 1–10. 10.1016/j.nbd.2017.03.009

Klingenberg, C. P. (2010). Evolution and development of shape: integrating quantitative approaches. Nature Reviews Genetics, 11(9), 623–635. 10.1038/nrg2829

Klingenberg, C. P. (2011). MorphoJ: an integrated software package for geometric morphometrics. Mol Ecol Resour, 11(2), 353–357. 10.1111/j.1755-0998.2010.02924.x

LaCombe, J. M., & Roper, R. J. (2020). Skeletal dynamics of Down syndrome: A developing perspective. Bone, 133, 115215. 10.1016/j.bone.2019.115215

Lamar, M., Foy, C. M. L., Beacher, F., Daly, E., Poppe, M., Archer, N., Murphy, D. G. M. (2011). Down syndrome with and without dementia: An in vivo proton Magnetic Resonance Spectroscopy study with implications for Alzheimer’s disease. NeuroImage, 57(1), 63–68. 10.1016/j.neuroimage.2011.03.073

Lana-Elola, E., Watson-Scales, S. D., Fisher, E. M. C., & Tybulewicz, V. L. J. (2011). Down syndrome: searching for the genetic culprits. Disease Models & Mechanisms, 4(5), 586–595. 10.1242/dmm.008078

Lee, Y., Ha, J., Kim, H. J., Kim, Y.-S., Chang, E.-J., Song, W.-J., & Kim, H.-H. (2009). Negative Feedback Inhibition of NFATc1 by DYRK1A Regulates Bone Homeostasis * Journal of Biological Chemistry, 284(48), 33343–33351. 10.1074/jbc.M109.042234

Letourneau, A., Santoni, F. A., Bonilla, X., Sailani, M. R., Gonzalez, D., Kind, J., Antonarakis, S. E. (2014). Domains of genome-wide gene expression dysregulation in Down’s syndrome. Nature, 508(7496), 345–350. 10.1038/nature13200

Li, H., Handsaker, B., Wysoker, A., Fennell, T., Ruan, J., Homer, N., Durbin, R. (2009). The Sequence Alignment/Map format and SAMtools. Bioinformatics, 25(16), 2078–2079. 10.1093/bioinformatics/btp352

Lin, A. L., Powell, D., Caban-Holt, A., Jicha, G., Robertson, W., Gold, B. T., Head, E. (2016). (1)H-MRS metabolites in adults with Down syndrome: Effects of dementia. Neuroimage Clin, 11, 728–735. 10.1016/j.nicl.2016.06.001

Llambrich, S., González-Colom, R., Wouters, J., Roldán, J., Salassa, S., Wouters, K., Martínez-Abadías, N. (2022). Green Tea Catechins Modulate Skeletal Development with Effects Dependent on Dose, Time, and Structure in a Down Syndrome Mouse Model. Nutrients, 14(19), 4167. 10.3390/nu14194167

Llambrich, S., González, R., Albaigès, J., Wouters, J., Marain, F., Himmelreich, U., Vande Velde, G. (2022). Multimodal in vivo Imaging of the Integrated Postnatal Development of Brain and Skull and Its Co-modulation With Neurodevelopment in a Down Syndrome Mouse Model [Original Research]. Frontiers in Medicine, 9. 10.3389/fmed.2022.815739

Locatelli, C., Onnivello, S., Antonaros, F., Feliciello, A., Filoni, S., Rossi, S., Lanfranchi, S. (2021). Is the Age of Developmental Milestones a Predictor for Future Development in Down Syndrome? Brain Sci, 11(5). 10.3390/brainsci11050655

Lott, I. T., & Dierssen, M. (2010). Cognitive deficits and associated neurological complications in individuals with Down’s syndrome. The Lancet Neurology, 9(6), 623–633. 10.1016/S1474-4422(10)70112-5

Love, M. I., Huber, W., & Anders, S. (2014). Moderated estimation of fold change and dispersion for RNA-seq data with DESeq2. Genome Biology, 15(12), 550. 10.1186/s13059-014-0550-8

Lyle, R., Gehrig, C., Neergaard-Henrichsen, C., Deutsch, S., & Antonarakis, S. E. (2004). Gene expression from the aneuploid chromosome in a trisomy mouse model of down syndrome. Genome Res, 14(7), 1268–1274. 10.1101/gr.2090904

Malak, R., Kostiukow, A., Krawczyk-Wasielewska, A., Mojs, E., & Samborski, W. (2015). Delays in Motor Development in Children with Down Syndrome. Medical science monitor : international medical journal of experimental and clinical research, 21, 1904–1910. 10.12659/MSM.893377

McCarron, M., McCallion, P., Reilly, E., Dunne, P., Carroll, R., & Mulryan, N. (2017). A prospective 20-year longitudinal follow-up of dementia in persons with Down syndrome. J Intellect Disabil Res, 61(9), 843–852. 10.1111/jir.12390

McElyea, S. D., Starbuck, J. M., Tumbleson-Brink, D. M., Harrington, E., Blazek, J. D., Ghoneima, A., Roper, R. J. (2016). Influence of prenatal EGCG treatment and Dyrk1a dosage reduction on craniofacial features associated with Down syndrome. Hum Mol Genet, 25(22), 4856–4869. 10.1093/hmg/ddw309

Même, S., Joudiou, N., Yousfi, N., Szeremeta, F., Lopes-Pereira, P., Beloeil, J., Herault, Y. and Même, W. (2014). In Vivo 9.4T MRI and 1H MRS for Evaluation of Brain Structural and Metabolic Changes in the Ts65Dn Mouse Model for Down Syndrome. World Journal of Neuroscience, 4, 152–163. 10.4236/wjns.2014.42018

Monteagudo, A. (2020). Holoprosencephaly. American Journal of Obstetrics & Gynecology, 223(6), B13–B16. 10.1016/j.ajog.2020.08.178

Motulsky, H. J., & Brown, R. E. (2006). Detecting outliers when fitting data with nonlinear regression a new method based on robust nonlinear regression and the false discovery rate. BMC Bioinformatics, 7, 123. 10.1186/1471-2105-7-123

Movsas, T. Z., Spitzer, A. R., & Gewolb, I. H. (2016). Ventriculomegaly in very-low-birthweight infants with Down syndrome. Developmental Medicine & Child Neurology, 58(11), 1167–1171. 10.1111/dmcn.13191

Muñiz Moreno, M. d. M., Brault, V., Birling, M.-C., Pavlovic, G., & Herault, Y. (2020). Chapter 4 Modeling Down syndrome in animals from the early stage to the 4.0 models and next. In M. Dierssen (Ed.), Prog Brain Res (Vol. 251, pp 91–143). Elsevier. 10.1016/bs.pbr.2019.08.001

Noll, C., Kandiah, J., Moroy, G., Gu, Y., Dairou, J., & Janel, N. (2022). Catechins as a Potential Dietary Supplementation in Prevention of Comorbidities Linked with Down Syndrome. Nutrients, 14(10). 10.3390/nu14102039

Nopoulos, P., Langbehn, D. R., Canady, J., Magnotta, V., & Richman, L. (2007). Abnormal Brain Structure in Children With Isolated Clefts of the Lip or Palate. Archives of Pediatrics & Adolescent Medicine, 161(8), 753–758. 10.1001/archpedi.161.8.753

Olmos-Serrano, J. L., Kang, H. J., Tyler, W. A., Silbereis, J. C., Cheng, F., Zhu, Y., Sestan, N. (2016). Down Syndrome Developmental Brain Transcriptome Reveals Defective Oligodendrocyte Differentiation and Myelination. Neuron, 89(6), 1208–1222. 10.1016/j.neuron.2016.01.042

Olmos-Serrano, J. L., Tyler, W. A., Cabral, H. J., & Haydar, T. F. (2016). Longitudinal measures of cognition in the Ts65Dn mouse: Refining windows and defining modalities for therapeutic intervention in Down syndrome. Experimental Neurology, 279, 40–56. 10.1016/j.expneurol.2016.02.005

Olson, L. E., Richtsmeier, J. T., Leszl, J., & Reeves, R. H. (2004). A Chromosome 21 Critical Region Does Not Cause Specific Down Syndrome Phenotypes. Science, 306(5696), 687–690. doi:10.1126/science.1098992

Pallast, N., Diedenhofen, M., Blaschke, S., Wieters, F., Wiedermann, D., Hoehn, M., Aswendt, M. (2019). Processing Pipeline for Atlas-Based Imaging Data Analysis of Structural and Functional Mouse Brain MRI (AIDAmri) [Original Research]. Frontiers in Neuroinformatics, 13. 10.3389/fninf.2019.00042

Patkee, P. A., Baburamani, A. A., Kyriakopoulou, V., Davidson, A., Avini, E., Dimitrova, R., Rutherford, M. A. (2020). Early alterations in cortical and cerebellar regional brain growth in Down Syndrome: An in vivo fetal and neonatal MRI assessment. Neuroimage Clin, 25, 102139. 10.1016/j.nicl.2019.102139

Pearlson, G. D., Breiter, S. N., Aylward, E. H., Warren, A. C., Grygorcewicz, M., Frangou, S., Pulsifer, M. B. (1998). MRI brain changes in subjects with Down syndrome with and without dementia. Dev Med Child Neurol, 40(5), 326–334.

Pinter, J. D., Eliez, S., Schmitt, J. E., Capone, G. T., & Reiss, A. L. (2001). Neuroanatomy of Down’s syndrome: a high-resolution MRI study. Am J Psychiatry, 158(10), 1659–1665. 10.1176/appi.ajp.158.10.1659

Pirozzi, F., Nelson, B., & Mirzaa, G. (2018). From microcephaly to megalencephaly: determinants of brain size. Dialogues Clin Neurosci, 20(4), 267–282. 10.31887/DCNS.2018.20.4/gmirzaa

Quinzi, F., Vannozzi, G., Camomilla, V., Piacentini, M. F., Boca, F., Bortels, E., Sbriccoli, P. (2022). Motor Competence in Individuals with Down Syndrome: Is an Improvement Still Possible in Adulthood? Int J Environ Res Public Health, 19(4). 10.3390/ijerph19042157

Ratiney, H., Coenradie, Y., Cavassila, S., van Ormondt, D., & Graveron-Demilly, D. (2004). Time-domain quantitation of 1H short echo-time signals: background accommodation. Magma, 16(6), 284–296. 10.1007/s10334-004-0037-9

Raveau, M., Nakahari, T., Asada, S., Ishihara, K., Amano, K., Shimohata, A., Yamakawa, K. (2017). Brain ventriculomegaly in Down syndrome mice is caused by Pcp4 dose-dependent cilia dysfunction. Hum Mol Genet, 26(5), 923–931. 10.1093/hmg/ddx007

Real de Asua, D., Quero, M., Moldenhauer, F., & Suarez, C. (2015). Clinical profile and main comorbidities of Spanish adults with Down syndrome. Eur J Intern Med, 26(6), 385–391. 10.1016/j.ejim.2015.05.003

Reeves, R. H., Irving, N. G., Moran, T. H., Wohn, A., Kitt, C., Sisodia, S. S., Davisson, M. T. (1995). A mouse model for Down syndrome exhibits learning and behaviour deficits. Nat Genet, 11(2), 177–184. 10.1038/ng1095-177

Reinholdt, L. G., Ding, Y., Gilbert, G. J., Czechanski, A., Solzak, J. P., Roper, R. J., Davisson, M. T. (2011). Molecular characterization of the translocation breakpoints in the Down syndrome mouse model Ts65Dn. Mammalian genome : official journal of the International Mammalian Genome Society, 22(11-12), 685–691. 10.1007/s00335-011-9357-z

Richtsmeier, J. T., Baxter, L. L., & Reeves, R. H. (2000). Parallels of craniofacial maldevelopment in down syndrome and Ts65Dn mice. Dev Dyn, 217(2), 137–145. 10.1002/(SICI)1097-0177(200002)217:2<137::AID-DVDY1>3.0.CO;2-N

Roberts, R. M., Mathias, J. L., & Wheaton, P. (2012). Cognitive Functioning in Children and Adults With Nonsyndromal Cleft Lip and/or Palate: A Meta-analysis. Journal of Pediatric Psychology, 37(7), 786–797. 10.1093/jpepsy/jss052

Rodrigues, M., Nunes, J., Figueiredo, S., Martins de Campos, A., & Geraldo, A. F. (2019). Neuroimaging assessment in Down syndrome: a pictorial review. Insights into Imaging, 10(1), 52. 10.1186/s13244-019-0729-3

Rondal, J. A. (2020). Down syndrome: A curative prospect? AIMS Neuroscience, 7(2), 168–193. 10.3934/Neuroscience.2020012

Roper, R. J., St John, H. K., Philip, J., Lawler, A., & Reeves, R. H. (2006). Perinatal loss of Ts65Dn Down syndrome mice. Genetics, 172(1), 437–443. 10.1534/genetics.105.050898

Roper, R. J., VanHorn, J. F., Cain, C. C., & Reeves, R. H. (2009). A neural crest deficit in Down syndrome mice is associated with deficient mitotic response to Sonic hedgehog. Mechanisms of Development, 126(3), 212–219. 10.1016/j.mod.2008.11.002

Rouillard, A. D., Gundersen, G. W., Fernandez, N. F., Wang, Z., Monteiro, C. D., McDermott, M. G., & Ma’ayan, A. (2016). The harmonizome: a collection of processed datasets gathered to serve and mine knowledge about genes and proteins. Database, 2016. 10.1093/database/baw100

Ruparelia, A., Pearn, M. L., & Mobley, W. C. (2012). Cognitive and pharmacological insights from the Ts65Dn mouse model of Down syndrome. Current Opinion in Neurobiology, 22(5), 880–886. 10.1016/j.conb.2012.05.002

Santin, M. D., Valabrègue, R., Rivals, I., Pénager, R., Paquin, R., Dauphinot, L., Potier, M. C. (2014). In vivo 1H MRS study in microlitre voxels in the hippocampus of a mouse model of Down syndrome at 11.71T. NMR Biomed, 27(10), 1143–1150. 10.1002/nbm.3155

Saran, N. G., Pletcher, M. T., Natale, J. E., Cheng, Y., & Reeves, R. H. (2003). Global disruption of the cerebellar transcriptome in a Down syndrome mouse model. Hum Mol Genet, 12(16), 2013–2019. 10.1093/hmg/ddg217

Shaw, P. R., Klein, J. A., Aziz, N. M., & Haydar, T. F. (2020). Longitudinal neuroanatomical and behavioral analyses show phenotypic drift and variability in the Ts65Dn mouse model of Down syndrome. Disease Models & Mechanisms, 13(9). 10.1242/dmm.046243

Smigielska-Kuzia, J., Boćkowski, L., Sobaniec, W., Sendrowski, K., Olchowik, B., Cholewa, M., Lebkowska, U. (2011). A volumetric magnetic resonance imaging study of brain structures in children with Down syndrome. Neurol Neurochir Pol, 45(4), 363–369. 10.1016/s0028-3843(14)60107-9

Smigielska-Kuzia, J., & Sobaniec, W. (2007). Brain metabolic profile obtained by proton magnetic resonance spectroscopy HMRS in children with Down syndrome. Adv Med Sci, 52 Suppl 1, 183–187.

Souchet, B., Duchon, A., Gu, Y., Dairou, J., Chevalier, C., Daubigney, F., Delabar, J. M. (2019). Prenatal treatment with EGCG enriched green tea extract rescues GAD67 related developmental and cognitive defects in Down syndrome mouse models. Scientific Reports, 9(1), 3914. 10.1038/s41598-019-40328-9

Souchet, B., Guedj, F., Penke-Verdier, Z., Daubigney, F., Duchon, A., Herault, Y., Delabar, J. M. (2015). Pharmacological correction of excitation/inhibition imbalance in Down syndrome mouse models [Original Research]. Frontiers in Behavioral Neuroscience, 9. https://www.frontiersin.org/articles/10.3389/fnbeh.2015.00267

Stagni, F., & Bartesaghi, R. (2022). The Challenging Pathway of Treatment for Neurogenesis Impairment in Down Syndrome: Achievements and Perspectives. Front Cell Neurosci, 16, 903729. 10.3389/fncel.2022.903729

Stagni, F., Giacomini, A., Emili, M., Guidi, S., & Bartesaghi, R. (2018). Neurogenesis impairment: An early developmental defect in Down syndrome. Free Radic Biol Med, 114, 15–32. 10.1016/j.freeradbiomed.2017.07.026

Stagni, F., Giacomini, A., Emili, M., Trazzi, S., Guidi, S., Sassi, M., Bartesaghi, R. (2016). Short- and long-term effects of neonatal pharmacotherapy with epigallocatechin-3-gallate on hippocampal development in the Ts65Dn mouse model of Down syndrome. Neuroscience, 333, 277–301. 10.1016/j.neuroscience.2016.07.031

Stagni, F., Giacomini, A., Guidi, S., Ciani, E., & Bartesaghi, R. (2015). Timing of therapies for Down syndrome: the sooner, the better [Review]. Frontiers in Behavioral Neuroscience, 9(265). 10.3389/fnbeh.2015.00265

Starbuck, J. M., Llambrich, S., Gonzàlez, R., Albaigès, J., Sarlé, A., Wouters, J., Martínez-Abadías, N. (2021). Green tea extracts containing epigallocatechin-3-gallate modulate facial development in Down syndrome. Scientific Reports, 11(1), 4715. 10.1038/s41598-021-83757-1

Starčuk Jr, Z., Štrbák, O., Starčuková, J., & Graveron-Demilly, D. (2009). Simulation of steady state free precession acquisition mode in coupled spin systems for fast MR spectroscopic imaging. Meas Sci Technol, 20.

Stefan, D., Di Cesare, F., Andrasescu, A., Popa, E., Lazariev, A., Vescovo, E., Graveron-Demilly, D. (2009). Quantitation of magnetic resonance spectroscopy signals: the jMRUI software package. Measurement Science and Technology, 20, 104035. 10.1088/0957-0233/20/10/104035

Steingass, K. J., Chicoine, B., McGuire, D., & Roizen, N. J. (2011). Developmental Disabilities Grown Up: Down Syndrome. Journal of Developmental & Behavioral Pediatrics, 32(7), 548–558. 10.1097/DBP.0b013e31822182e0

Stringer, M., Abeysekera, I., Dria, K. J., Roper, R. J., & Goodlett, C. R. (2015). Low dose EGCG treatment beginning in adolescence does not improve cognitive impairment in a Down syndrome mouse model. Pharmacol Biochem Behav, 138, 70–79. 10.1016/j.pbb.2015.09.002

Stringer, M., Abeysekera, I., Thomas, J., LaCombe, J., Stancombe, K., Stewart, R. J., Roper, R. J. (2017). Epigallocatechin-3-gallate (EGCG) consumption in the Ts65Dn model of Down syndrome fails to improve behavioral deficits and is detrimental to skeletal phenotypes. Physiology & Behavior, 177, 230–241. 10.1016/j.physbeh.2017.05.003

Sunkin, S. M., Ng, L., Lau, C., Dolbeare, T., Gilbert, T. L., Thompson, C. L., Dang, C. (2013). Allen Brain Atlas: an integrated spatio-temporal portal for exploring the central nervous system. Nucleic Acids Res, 41(Database issue), D996-d1008. 10.1093/nar/gks1042

Suri, S., Tompson, B. D., & Cornfoot, L. (2010). Cranial base, maxillary and mandibular morphology in Down syndrome. The Angle Orthodontist, 80(5), 861–869. 10.2319/111709-650.1

Tallino, S., Winslow, W., Bartholomew, S. K., & Velazquez, R. (2022). Temporal and brain region-specific elevations of soluble Amyloid-β40-42 in the Ts65Dn mouse model of Down syndrome and Alzheimer’s disease. Aging Cell, 21(4), e13590. 10.1111/acel.13590

Thomas, J. R., LaCombe, J., Long, R., Lana-Elola, E., Watson-Scales, S., Wallace, J. M., Roper, R. J. (2020). Interaction of sexual dimorphism and gene dosage imbalance in skeletal deficits associated with Down syndrome. Bone, 136, 115367. 10.1016/j.bone.2020.115367

Thomas, J. R., & Roper, R. J. (2021). Current Analysis of Skeletal Phenotypes in Down Syndrome. Current Osteoporosis Reports, 19(3), 338–346. 10.1007/s11914-021-00674-y

Thomas, J. R., Sloan, K., Cave, K., Wallace, J. M., & Roper, R. J. (2021). Skeletal Deficits in Male and Female down Syndrome Model Mice Arise Independent of Normalized Dyrk1a Expression in Osteoblasts. Genes, 12(11), 1729. https://www.mdpi.com/2073-4425/12/11/1729

Treit, S., Zhou, D., Chudley, A. E., Andrew, G., Rasmussen, C., Nikkel, S. M., Beaulieu, C. (2016). Relationships between Head Circumference, Brain Volume and Cognition in Children with Prenatal Alcohol Exposure. PLoS One, 11(2), e0150370. 10.1371/journal.pone.0150370

van den Boogaart, A., van Ormondt, D., Pijnappel, W. W. F., de Beer, R. and Ala Korpel, M. (1994). Removal of the residual water resonance from 1H magnetic resonance spectra. Mathematics of Signal Processing III,, 175–195.

Vanherp, L., Poelmans, J., Weerasekera, A., Hillen, A., Croitor-Sava, A. R., Sorrell, T. C., Himmelreich, U. (2021). Trehalose as quantitative biomarker for in vivo diagnosis and treatment follow-up in cryptococcomas. Transl Res, 230, 111–122. 10.1016/j.trsl.2020.11.001

Vicente, A., Bravo-González, L.-A., López-Romero, A., Muñoz, C. S., & Sánchez-Meca, J. (2020). Craniofacial morphology in down syndrome: a systematic review and meta-analysis. Scientific Reports, 10(1), 19895. 10.1038/s41598-020-76984-5

Vilardell, M., Rasche, A., Thormann, A., Maschke-Dutz, E., Pérez-Jurado, L. A., Lehrach, H., & Herwig, R. (2011). Meta-analysis of heterogeneous Down Syndrome data reveals consistent genome-wide dosage effects related to neurological processes. BMC Genomics, 12, 229. 10.1186/1471-2164-12-229

Vogels, A., & Fryns, J. P. (2006). Pfeiffer syndrome. Orphanet J Rare Dis, 1, 19. 10.1186/1750-1172-1-19

Weerasekera, A., Sima, D. M., Dresselaers, T., Van Huffel, S., Van Damme, P., & Himmelreich, U. (2018). Non-invasive assessment of disease progression and neuroprotective effects of dietary coconut oil supplementation in the ALS SOD1(G93A) mouse model: A (1)H-magnetic resonance spectroscopic study. Neuroimage Clin, 20, 1092–1105. 10.1016/j.nicl.2018.09.011

Weisfeld-Adams, J. D., Tkachuk, A. K., Maclean, K. N., Meeks, N. L., & Scott, S. A. (2016). A de novo 2.78-Mb duplication on chromosome 21q22.11 implicates candidate genes in the partial trisomy 21 phenotype. NPJ Genom Med, 1. 10.1038/npjgenmed.2016.3

Wilhoit, L. F., Scott, D. A., & Simecka, B. A. (2017). Fetal Alcohol Spectrum Disorders: Characteristics, Complications, and Treatment. Community Ment Health J, 53(6), 711–718. 10.1007/s10597-017-0104-0

Wozniak, J. R., Riley, E. P., & Charness, M. E. (2019). Clinical presentation, diagnosis, and management of fetal alcohol spectrum disorder. Lancet Neurol, 18(8), 760–770. 10.1016/s1474-4422(19)30150-4

Xicota, L., Rodríguez, J., Langohr, K., Fitó, M., Dierssen, M., & de la Torre, R. (2020). Effect of epigallocatechin gallate on the body composition and lipid profile of down syndrome individuals: Implications for clinical management. Clinical Nutrition, 39(4), 1292–1300. 10.1016/j.clnu.2019.05.028

Yin, X., Jin, N., Shi, J., Zhang, Y., Wu, Y., Gong, C.-X., Liu, F. (2017). Dyrk1A overexpression leads to increase of 3R-tau expression and cognitive deficits in Ts65Dn Down syndrome mice. Scientific Reports, 7(1), 619. 10.1038/s41598-017-00682-y

Zis, P., & Strydom, A. (2018). Clinical aspects and biomarkers of Alzheimer’s disease in Down syndrome. Free Radic Biol Med, 114, 3–9. 10.1016/j.freeradbiomed.2017.08.024

